# Inverted genomic regions between reference genome builds in humans impact imputation accuracy and decrease the power of association testing

**DOI:** 10.1101/2022.02.19.481174

**Authors:** Xin Sheng, Lucy Xia, David V. Conti, Christopher A. Haiman, Linda Kachuri, Charleston W. K. Chiang

## Abstract

Over the last two decades, the human reference genome has undergone multiple updates as we complete a linear representation of our genome. There are two versions of human references currently used in the biomedical literature, GRCh37/hg19 and GRCh38, and conversions between these versions are critical for quality control, imputation, and association analysis. In the present study we show that genomic coordinates for single nucleotide variants (SNVs) in regions inverted between different builds of the reference genome are erroneously converted by the TOPMed imputation server. Depending on the array type, we estimate the inappropriate conversions of variant coordinates would occur in about 2-5 Mb of the genome. Independent of the imputation server, however, in-house conversions of variant coordinates are routine yet impact of mapped variants to these inverted regions and its downstream analytical consequences are underrecognized. We observed that errors for palindromic variants in these inverted regions cannot be detected by standard quality control procedures and destabilize the local haplotype structure, leading to loss of imputation accuracy and power in association analyses. Though only a small proportion of the genome is affected, we show that these regions include important disease susceptibility variants that would be lost due to poor imputation. For example, we show that a known locus associated with prostate cancer on chr10 would have its association P-value drop from 2.86×10^-7^ to 0.0011 in a case-control analysis of 20,286 Africans and African Americans (10,643 cases and 9,643 controls). We devise a straight-forward heuristic based on the popular tool *liftOver* that can easily detect and correct these variants in the inverted regions between genome builds to locally improve imputation accuracy.

## Introduction

In the ensuing 13 years since the completion of the human genome project, the human reference genome assembly has undergone at least 18 major updates and numerous patches[1,2]. The latest genome build from the Genome Reference Consortium (GRC), GRCh38, was released in 2013[1], and was most recently patched in 2019. The human reference genome assembly plays an essential part in etiologic and translational research by providing a common roadmap for deciphering the location of genes and functional regions of the human genome and discovering genetic variation that affects disease susceptibility. Reference genome build and variant position are the most fundamental pieces of information that are reported in genetic association studies, providing researchers with the proper context when trying to interpret, replicate, or meta-analyze reported associations. However, at the time of this writing, some 8 years since the last major update of the human reference genome assembly, both GRCh38 and the previous prevailing reference build (GRCh37/hg19, released in 2009) continue to co-exist in literature. For example, the Genome Aggregation Database (gnomAD[3]) continues to maintain GRCh37 and GRCh38 versions of the database. In fact, polygenic score models curated in the Polygenic Score Catalog[4] are largely still in GRCh37/hg19. For example, for height (EFO: 0004339) and BMI (EFO: 0004340), 18 out of 19 released polygenic score models with explicit genome build information at the time of writing were provided in GRCh37/hg19, with the one remaining model in GRCh36. While there is a movement towards utilizing GRCh38 as the main reference genome assembly, many datasets remained in GRCh37/hg19 as there exists a wealth of information generated in this coordinate system and a continued reliance on GRCh37 by numerous computational tools for downstream analysis[5]. As such, it is a constant need to update the coordinate systems of these legacy datasets in order to harmonize them with the genetic data produced by the latest genotyping platform or sequencing technology.

Re-mapping or re-alignment of genetic datasets into a different reference build is computationally expensive, therefore bioinformatic conversions between assembly builds, using tools such as *liftover* from the UCSC Genome Browser ([6]; https://genome.ucsc.edu/cgi-bin/hgLiftOver), were developed to harmonize different datasets and enable analyses essential for genetic discovery. In the case of *liftover*, the conversion process utilizes a chain file that provides a mapping of contiguous positions from one genome build to another. Other similar tools also exist, such as *CrossMap* and *Remap* ([*7,8];* https://www.ncbi.nlm.nih.gov/genome/tools/remap). To facilitate standardizing the coordinate system of genetic datasets and enable seamless downstream meta-analysis, the TOPMed imputation server[9,10] (https://imputation.biodatacatalyst.nhlbi.nih.gov/) can internally convert GRCh37 input dataset into the GRCh38 coordinate system. Once standardized on the same coordinate system between input dataset and reference dataset, genotype imputation, a process of estimating unobserved genotypes in an input dataset (typically genome-wide single nucleotide variant, or SNV, data from genotyping microarrays) from the haplotypes of a reference panel, can proceed. Imputation is now an essential tool to improve the coverage and power of a genome-wide association study (GWAS), facilitate downstream fine-mapping of a target region, and enable meta-analysis in consortiums when multiple datasets were genotyped on different array platforms[11]. Because imputation relies on a reference panel of haplotypes, it is essential that the input data is coded in the same genomic coordinates with forward strand alleles as that of the imputation reference. (Note that we use forward strand as synonymous as the “plus” strand, defined as the strand with its 5’ end at the tip of the short arm of a chromosome. This is different from the forward strand in the context of dbSNP submissions[12].)

In the current study, we observed an error in the conversion between reference genome builds by the TOPMed imputation server. This error is localized specifically to regions that are inverted between GRCh37/hg19 and GRCh38/hg38, causing directly genotyped SNVs to be dropped and then imputed at lower quality. However, our observed issue is beyond a potential bioinformatic bug in the TOPMed imputation server. In-house conversions of genomic coordinates using tools like *liftOver* are routine, and often done to conform the coordinate systems across multiple datasets for GWAS meta-analysis. In this context, we further found that even manually converting input files to GRCh38 prior to imputation does not completely resolve this issue. Since the conversion involves inverted sequences between genome builds, the flipped alleles, particularly for palindromic SNVs (i.e. A/T transversion or C/G transversion variants), often escape detection. Attempts to impute such GWAS dataset would be hampered by destabilized local haplotype structure, leading to poorer quality and decreased power in association testing. To overcome this underrecognized analysis problem, we developed a heuristic based on converting the basepair (bp) immediately before and after the focal SNV to deduce whether the SNV is found in an inverted region. We showed that our approach can detect and correct all impacted SNVs in the array platforms we tested. In empirical analysis using prostate cancer as an example, our approach would identify important associations that would otherwise be missed due to poor imputation quality.

## Results

As of freeze 8 (r2) of the TOPMed imputation server (last accessed on 12/20/2021), the server allows internal conversion of genome build so users can submit imputation-ready GWAS datasets for imputation without explicitly converting the coordinate from GRCh37 to GRCh38. We observed that this practice results in a number of directly genotyped SNVs on the array being labeled “typed only” after the server converts the genome build, suggesting that these SNVs are not found in the TOPMed imputation reference panel despite being common in the population. This also created instances of imputed variants immediately adjacent to the “typed only” variant in the output with complementary alleles (**Table 1**).

**Table 1:** Examples of unexpected typed-only SNVs observed from the imputation using server liftOver. These directly genotyped SNVs become “typed-only” SNVs, meanwhile a SNV with complementary alleles is imputed at the immediate adjacent location to the “typed-only” variant.

| Impute SNV information (hg19/GRCh37) |  |  |  |  | Records in TOPMed imputation output (hg38/GRCh38) |  |  |  |
| --- | --- | --- | --- | --- | --- | --- | --- | --- |
| Chr | Pos | A1 | A2 | SNPID | SNV entry | Type | REF | ALT |
| 1 | 145095477 | A | G | rs28549707 | Chr1:120177450:G:A | typedOnly | -- | -- |
|  |  |  |  |  | Chr1:120177451:C:T | Imputed | C | T |
| 1 | 144474542 | C | T | rs10907360 | chr1:120959671:T:C | typedOnly | -- | -- |
|  |  |  |  |  | chr1:120959672:A:G | Imputed | A | G |
| 1 | 145755813 | C | A | rs10157535 | chr1:145679248:A:C | typedOnly | -- | -- |
|  |  |  |  |  | chr1:145679249:T:G | Imputed | T | G |

If the genome build is updated using UCSC’s tool *liftOver* (https://genome.ucsc.edu/cgi-bin/hgLiftOver) prior to submission to the imputation server, we found that *liftOver* maps these SNVs to a coordinate that is off by 1 bp compared to the mapped coordinate from the imputation server. Therefore, because of the 1-bp shift in the implemented build conversion in TOPMed server, these SNVs would need to be imputed back by the server.

Genome-wide examinations of this occurrence using several GWAS datasets indicated that this is not a general phenomenon across the genome. The vast majority (> 99.8%) of the SNVs found on the GWAS arrays that we examined would be correctly mapped from GRCh37 to GRCh38 by the imputation server. However, we noticed when the conversion does fail, these SNVs are not randomly scattered along the genome, but tend to cluster into regions (**Supplemental Figure 1**; **Supplemental Table 1**). Since the imputed versions of these SNVs tend to have complementary alleles (e.g. rs28549707 from **Table 1**), we suspected that these are the regions in which the orientations are reversed between GRCh37 and GRCh38. Specifically, the reference alleles for these SNVs reside on the forward (plus) strands in GRCh37 but reverse (minus) strands in GRCh38. Previous genome-wide alignments of GRCh38 to GRCh37 reference revealed 11 Mb (0.37% of total length) of inverted sequences. Though we could not locate a complete record of all inversions, the ten longest inverted regions previously reported in the supplement of Schneider et al.[1] coincided with our mapping of SNVs erroneously converted by the imputation server. We thus termed these regions as Between-Builds Inverted Sequence (BBIS) regions.

In the BBIS regions, information from these directly genotyped SNVs could not be used to inform the local haplotype structure. We thus expected that the lack of genotype information would lower the imputation accuracy locally around BBIS regions. We first assessed whether this issue could be resolved by manual conversion of input data from GRCh37 to GRCh38 prior to imputation, a common practice in quality control pipelines for GWAS datasets. However, because it is underrecognized that regions of the genome can be inverted between genome builds, changing the positions of SNVs in BBIS regions without mapping the alleles to the complementary strand would still be taken as an allelic mismatch for non-palindromic SNVs (they will be flagged as strand flip in “snps-excluded.txt” file in the imputation server). This resulted in the same fate that these SNVs are dropped from imputation. These challenges are exacerbated for palindromic SNVs, which would have been retained for imputation even though their alleles are reversed relative to the reference panel, thereby disrupting the local haplotype structure. In principle, one approach to fix the strand flip issue for non-palindromic SNVs is running quality control checks in the imputation server first, then inspecting and correcting as needed SNVs with strand flip. However, palindromic SNVs could not be reliably detected, and for ancestrally diverse populations with limited reference samples, it would not be advisable to assign alleles for palindromic SNVs based on alternate allele frequency. Even with available population-specific frequencies, accurately inferring palindromic alleles for variants with intermediate (>40%) frequencies becomes challenging.

Having recognized this specific issue, one can devise a solution using multiple approaches. We solved this problem by devising a heuristic that we termed *triple*-*liftOver* that could identify SNVs that fall within BBIS regions. In essence, we convert the genome build using *liftOver* not just for the bp of the SNV of interest (*i*), but also for the bp before (*i*-1) and after the SNV (*i*+1). If the sequence is inverted around this SNV location (*j*) in the new build, the succeeding base will become the preceding base and the preceding base will become the succeeding base (**Figure 1**). To account for the rare event that the three-bp sequence is no longer contiguous in the new genome reference build, we lessen the constraint to flag a site as inverted if either of the neighboring base shows the orientation change.

**Figure 1:**
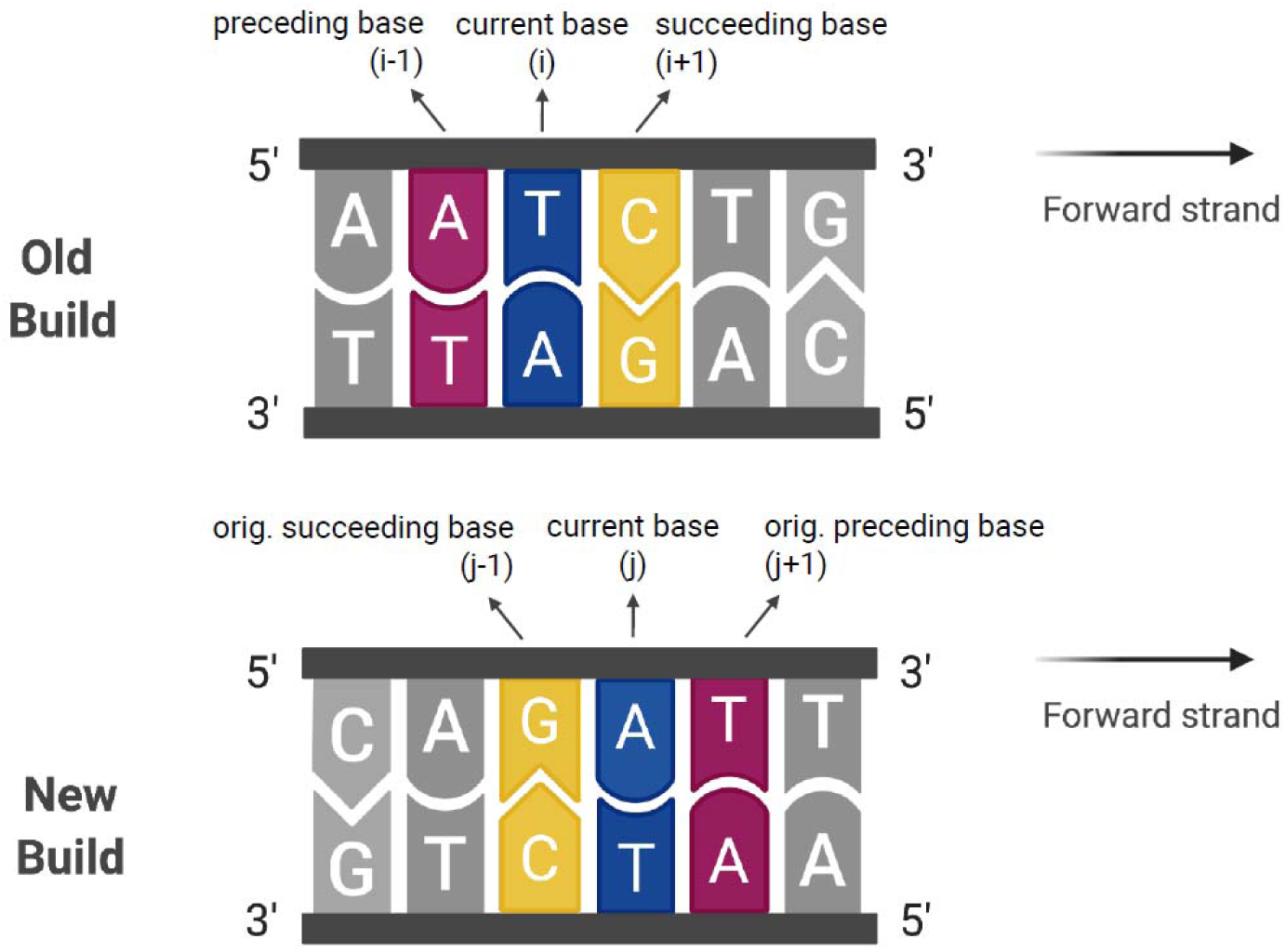
schematic of the *triple-liftOver* heuristic. To illustrate how the *triple-liftOver* approach works, a SNV is assumed to reside at base i with reference allele T on the forward strand in the old build. Its preceding base (*i-1*) is A and succeeding base (*i+1*) is C. When this segment of the sequence falls into a BBIS region, the prior forward strand would become the reverse strand in the new build. The corresponding SNV site would be at base (*j*) with reference allele A (complementary to T) in the new build. Its new preceding base G (*j-1)* is the original succeeding base C (*i+1*) on the opposite strand. Similarly, its new succeeding base T (*j+1*) is the original preceding base A (*i-1*) on the opposite strand. Our heuristic relies on the exact inversion of either adjacent bp to the focal position to identify SNVs found in the BBIS region.

We validated the *triple*-*liftOver* approach using three GWAS datasets for Prostate Cancer[13] that were genotyped on three different Illumina platforms: Human1M-Duo (AAPC1M), Consortium-OncoArray (ONCO-AAPC) and H3Africa (AAPC-H3) (**Method**). Using all non-palindromic SNVs on each array platform, we aimed to identify the proportion of server-identified strand flips that would be detected using the *triple*-*liftOver* approach. Across the three Illumina arrays our approach identified all non-palindromic SNVs (733 for Human1M-Duo, 410 for Consortium-OncoArray and 1325 for H3Africa) flagged by the imputation server as strand flips. Given the 100% sensitivity in identifying non-palindromic SNVs residing in the BBIS regions, we used our approach to further identify 33, 53, and 59 palindromic SNVs for the Human1M-Duo, Consortium-OncoArray and H3Africa array, respectively, that would escape detection by the imputation server. The detected palindromic and non-palindromic SNVs in BBIS regions cluster into 36, 25, and 51 stretches spanned by consecutive markers on each of Human1M-Duo, Consortium-OncoArray, and H3Africa, respectively, together covering approximately 2.7Mb to 5.4Mb in length, consisting of 501 to 1578 consecutive markers found on an array (**Supplemental Table 2; Method**).

We further validated the detected BBIS-region SNVs by checking the reference allele in the human reference GRCh37 and GRCh38. Indeed, for all 1927 unique SNV sites found in BBIS region across the three arrays, either the reference allele or the alternative allele of the SNV in GRCh37 is complementary to the reference allele in GRCh38. Illumina array designs are biased towards non-palindromic SNVs to avoid strand confusions, hence we detected relatively fewer palindromic SNVs compared to the non-palindromic SNVs that were localized in the BBIS regions. We expect our approach to identify greater number of palindromic SNVs for other array platforms without this bias.

The *triple*-*liftOver* approach takes a variant-centric approach: an input list of SNV coordinates is used to identify a subset exhibiting behaviors consistent with the SNV falling within the BBIS regions. To evaluate how many SNVs across the genome may fall into BBIS regions, we interrogated each biallelic SNV from IMPUTE2 1000 genomes phase3 legend file (https://mathgen.stats.ox.ac.uk/impute/1000GP_Phase3.html) with *triple*-*liftOver*. Out of 80,978,435 SNVs with unique chromosome location in GRCh37 that can be lifted over to the same chromosome in GRCh38, our program identified 208,930 as inverted sites (0.258%). Among the 208,930 inverted sites, 208,164 sites (99.63%) have both adjacent bp in reverse order in GRCh38, while 766 have one of the adjacent bp in reverse order. 80,767,799 out of 80,978,435 SNVs (99.74%) sites would maintain its relative order with both neighboring bases between GRCh37 and GRCh38. For the remaining 1,706 SNVs (0.0021%), all except one have one adjacent bp in the same orientation as in GRCh37, suggesting that they are not in a BBIS region, but the position immediately upstream or downstream of the SNV site may be deleted or translocated between genome builds. The one exception is a SNV disjointed with both of its adjacent bp in GRCh38. We further validated these 208,930 inverted sites by comparing the reference allele between GRCh37 and GRCh38 human reference sequences. In 99.99% of the sites (208,903 out of 208,930), one of the two alleles of the SNV in GRCh37 is complementary to the reference allele in GRCh38. For the remaining 27 SNVs (0.01%), neither the reference allele nor the alternative allele of the SNV is complementary to the reference allele in GRCh38, but BLAT of nearby sequences suggest that the sequences are indeed inverted, suggesting an annotation error for the SNV allele or a sequencing error in one of the reference genome build. The genomic locations of these inverted sites together with the ones detected from our three GWAS arrays are shown in **Supplemental Figure 1 and Supplemental Table 3**.

We next examined the impact on imputation quality due to these SNVs falling within BBIS regions. BBIS regions are sparse and the proportion of SNVs falling in these regions are low (∼0.26%-0.37% of the human genome[1]), therefore uncorrected SNVs that are inverted between genome builds do not result in an appreciable impact on the overall imputation quality (**Supplemental Figure 2**). Instead, their impact on imputation is confined more locally. To better visualize the local impact on imputation accuracy, we grouped nearby (not necessarily contiguous) inverted SNVs that were detected by *triple*-*liftOver* into approximately 18, 15, and 27 regions across the genome for each of the three arrays we examined, spanning ∼2.2Mb to 3.8Mb and comprised of 475-1375 SNVs (**Method**). We then extended each merged region by 500kb to systematically examine the imputation quality locally within the BBIS region and the surrounding non-inverted regions. In general, we observed that correcting the strand for both palindromic and non-palindromic SNVs will improve imputation accuracy locally, particularly for larger BBIS regions.

Using the region around 46Mb on chr10 from OncoArray (ONCO-AAPC) as an illustrative example, we describe the impact of three approaches to dealing with SNVs in BBIS regions when imputing a dataset in GRCh37. In the first approach, we submit the array data in GRCh37 for imputation, relying on using the TOPMed imputation server to convert the coordinates into GRCh38 prior to imputation (**Figure 2A**). In the second approach, we manually converted the coordinates into GRCh38 using *liftOver*, and manually corrected the detectable strand issues for all non-palindromic SNVs (**Figure 2B**). In the last approach, we used *triple*-*liftOver* to systematically identify SNVs falling in BBIS regions and flipped the strand for all detected SNVs, including both palindromic and non-palindromic SNVs (**Figure 2C**).

**Figure 2:**
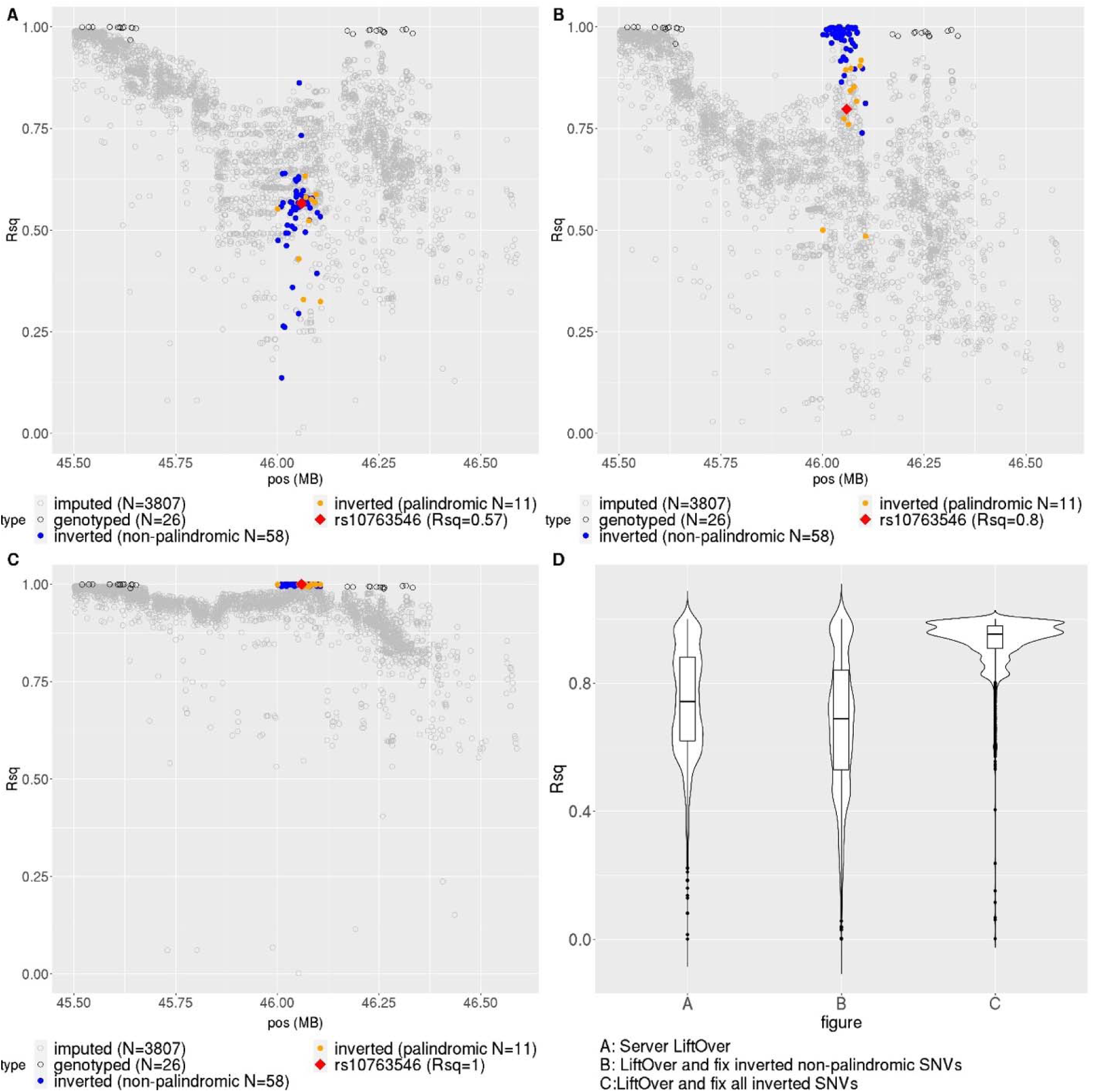
Three scenarios of imputation for the 46Mb region (GRCh38) on chr10 in ONCO-AAPC. We compared three reasonable approaches to impute this 1.1 Mb region (chr10:45,500,727-46,605,935) where the MSMB gene (chr10:46,033,313-46,046,269) resides: (A) rely on the imputation server to convert input array data in GRCh37 to GRCh38 prior to imputation; (B) manually convert the GRCh37 coordinates to GRCh38 and fix strands for all non-palindromic SNVs prior to imputation; (C) manually convert GRCh37 coordinates to GRCh38 and fix strands for all SNVs detected by *triple-liftOver* prior to imputation. Note that without *triple-liftOver* or other similar approaches, only non-palindromic SNVs in BBIS regions would be identified. In (D) we show violin plot of Rsq distribution as a measure of imputation accuracy for each scenario. Only common variants (MAF >= 0.01) are shown in plot A through C.

In the first scenario, all genotyped SNVs (N=69) falling within this chr10 BBIS region were dropped and then re-imputed with relatively poor Rsq (**Figure 2A**), as the TOPMed server genome build conversion in these regions would result in a 1-bp shift, as described earlier (**Table 1**). Correcting for strand flips at non-palindromic SNVs (N=58) improved imputation quality for directly genotyped SNVs as they became recognized by the imputation server. However, the imputation quality for other SNVs in the region remained poor (**Figure 2B**). The first approach yielded a mean Rsq 0.74 compared to 0.67 in the second approach (*P* < 2.2×10^-16^ by Wilcoxon Rank Sum Test), which failed to correct palindromic SNVs (N = 12; **Figure 2D**). The lower Rsq in the second approach suggests that the uncorrected palindromic SNVs disrupt local haplotype structures due to their incompatible genotypes for these SNVs. After fixing the strand issue for these 12 SNVs, the overall imputation quality is increased to mean Rsq of 0.93 across the 3903 common variants in this 1.1Mb region surrounding a BBIS region (*P* < 2.2×10^-16^ compared to both approach 1 and approach 2 by Wilcoxon Rank Sum Test; **Figure 2C**).

Improvement in imputation accuracy was generally observed across all 28 regions and the three array platforms that we tested, though the magnitude of the improvement depends on the number of palindromic and non-palindromic SNVs falling within an inverted region and how well the imputation algorithm can impute these SNVs without their actual genotypes. We also observed qualitatively similar pattern using OncoArray dataset from Latinos (ONCO-LAPC), suggesting that the adverse effect caused by these inverted SNVs is not a phenomenon unique to the African and African American populations we examined (**Supplemental Figure 3-6; Method**).

Despite the low prevalence of SNVs found in BBIS regions their impact on phenotypic associations is appreciable. The BBIS region on chr10 includes the MSMB gene (chr10:46033313-46046269 on GRCh38), which contains a consistently replicated susceptibility signal rs10993994 for prostate cancer [11] (**Figure 2**). We tested a palindromic SNV within this locus, rs10763546, for its association with prostate cancer in 10,643 cases and 9,643 controls. This variant was directly genotyped on one of the three Illumina arrays (ONCO-AAPC), but had to be imputed on the other two array platforms. When submitted for imputation in hg19, rs10763546 would have to be imputed in all three array platforms because of the 1-bp error in genome build conversion. In this case, the local region as well as the SNV were poorly imputed (Rsq ∼0.57-0.66 for rs10763546). As a result, we found only little evidence of association (OR = 1.11, P_meta_ = 0.0011 after meta-analysis across ONCO-AAPC, AAPC1M, and AAPC-H3; Supplemental Table 4), which would not be declared genome-wide significant in a standard GWAS. If we converted the genome build manually using *liftOver* prior to submission for imputation, the palindromic SNV rs10763546 would not be detected as a strand flip. Instead, the incompatible haplotypic pattern in the region due to uncorrected palindromic SNVs would destabilize the haplotype models and result in relatively poor imputation, even for a directly genotyped variant on the OncoArray (Rsq = 0.80). Association testing in ONCO-AAPC would result in effectively a null association for rs10763546 (OR = 1.04, P = 0.42; **Supplemental Table 4**), and only a marginally improved meta-analysis association signal (OR = 1.10, *P*_meta_ = 0.00020; **Supplemental Table 4**). Finally, when we applied *triple*-*liftOver* to identify and correct all SNVs requiring strand flips in this region, the imputation accuracy for this SNV and the local region is improved (Rsq = 1 for rs10763546), resulting in a meta-analysis result of OR = 1.14 and P = 2.86×10^-7^, nearly genome-wide significant. We repeated the same analysis for this SNV in ONCO-LAPC (1192 cases and 1052 controls). The version with fixing all strand issue yields an OR = 1.41 and *P* = 6.52×10^-8^ compared to OR = 1.19 and *P* = 0.045 from the server *liftOver* version (**Supplemental Table 4**). Taken together, our proof-of-principle analysis demonstrated the ability to detect or further fine-map this associated locus would be hampered simply due to a change in reference genome build.

We also examined all of the trait-associated SNVs found in the GWAS catalog[14] and the Global Biobank Engine[15], which is primarily based on UK Biobank data (**Method**). For the GWAS catalog, we examined 162K unique SNVs. Among which, 151 SNVs are found in the BBIS regions (∼0.09%, in a total of 277 variant-trait pairs, **Supplemental Table 5**). In the Global Biobank Engine, we obtained 2.1M SNV variant-trait pairs that would collapse into 300K unique SNVs, and 409 SNVs would be in the BBIS regions (∼0.14%, in a total of 3385 variant-trait pairs, **Supplemental Table 6**). These estimates are broadly consistent with the estimated proportion of SNVs falling into the BBIS regions (∼0.26%-0.37% of the human genome[1]). Functional annotation of SNVs in BBIS regions revealed 1961 nonsynonymous variants and 678 substitutions in 54 genes that were predicted to be deleterious by Sift and PolyPhen. Among non-coding SNVs, 187 variants had CADD scores greater than 20, corresponding to the top 1% most deleterious substitutions in the genome (**Supplemental Table 7, Supplemental Figure 7**). Taken together, we demonstrate that BBIS regions harbor variants of significance to multiple phenotypes and that ignoring allelic errors within these regions could lead to missed opportunities in identifying a genetic association.

## Discussion

Our current investigation was motivated by an anomaly where directly genotyped common SNVs on the array appeared to be absent from the imputation reference panel using the TOPMed imputation server. Our deep dive into this anomaly revealed that the problem is more general than a minor bioinformatic error in the genome build conversion protocol implemented by the TOPMed server. It is ultimately caused by genomic regions that are inverted between different builds of the human reference, which is largely overlooked in standard bioinformatic analysis. We termed these regions Between-Build Inverted Sequence (BBIS) regions. Without knowing that a set of SNVs fall within the BBIS regions, imputation of the region could be impacted as genotyped SNVs could be removed from analysis due to errors in genome build conversion or incompatible alleles due to the inversion, or destabilized local haplotype structure due to undetected palindromic SNVs. Since SNVs in BBIS regions represent a small proportion of the genome, these minor inconsistencies are often ignored. For example, a few hundred SNVs identified as potential strand flips are often regarded as potential annotation errors, rather than a signature of pervasive strand-flip issues. However, as we have shown, ignoring these issues when preparing a dataset for imputation and GWAS analysis could also cause functional, trait-associated, variants to be undetected.

We devised a simple algorithm that we call *triple*-*liftOver* to detect SNV sites in BBIS regions with high sensitivity. Once detected, records for these SNVs can be modified to properly reflect the strand and alleles before imputation. This program can also help identify inverted SNVs in meta-analysis where individual results came from different genome build versions (such as phase3-imputed vs. TOPMed-imputed results). Failing to detect allele incompatibility for non-palindromic SNVs could also result in SNVs being meta-analyzed as multi-allelic variants and palindromic SNVs being meta-analyzed incorrectly.

We focused our evaluation on the TOPMed imputation server because it is the current state-of-the-art imputation server. However, we expect our findings to be relevant for users of other imputation services[16–18] or if they perform custom imputations[19–21]. In fact, given the routine use of *liftOver* for in-house conversions of genomic coordinates, analytical intricacies due to regions inverted between genome builds have likely gone generally unrecognized. Our simply algorithm is light-weight and built on-top of *liftOver*, and thus could easily be slipped into bioinformatic pipelines already utilizing *liftOver* and its chain files, though other approaches to convert genomic coordinates are possible[7,8]. Another way to avoid the issues caused by BBIS regions is to define alleles invariant across genome build, such as the TOP and BOT strand definitions for variants in the Illumina platforms[12]. However this does not apply to other genotyping platforms, such annotation are not routinely associated with individual genetic datasets, and a change would require also systematic updates of the imputation reference panel.

There are a few issues that our heuristic has yet to address. For one, even though the field has not completely converted to using GRCh38 as the reference, a new human genome reference T2T-CHM13 has been released[22]. A detailed comparison of overlapping regions between GRCh38 and T2T-CHM13 is needed to reveal the extent of the BBIS regions between these two builds. But given a chain file between the two builds, our method should readily identify variants in the BBIS regions. Moreover, we restricted our analyses and demonstrations to SNVs, although indels and other structural variants may also be affected. In principle, if the boundaries of the structural variants are called reliably, the same heuristic can be used to determine if the structural variants also need to be inverted onto the complementary strand, but this has not been thoroughly tested. Moreover, the changes in the human genome reference assembly across builds could be far more complex than what we have realized in some regions. Our variant-based *triple*-*liftOver* approach relies on *liftOver* to convert the coordinates of basepairs around the focal SNV site to infer whether the focal SNV is found in an inverted region. If the SNV resides in a region which has changed dramatically across genome builds, our approach may not yet be able to tackle those complicated scenarios. Finally, even though in our demonstrations we focused on the conversion between GRCh37 to GRCh38 in humans. In principle, our heuristic would be equally effective for other genome builds and in other species, as long as there is a uniform way to convert coordinates across genome builds. Therefore, we expect the application of our heuristic to go beyond the human species. Given the simplicity of our heuristic, there is little computational cost to apply this heuristic in standard quality-control pipelines.

## Methods

### GWAS datasets and statistical analysis

We used four in-house prostate cancer GWAS datasets to examine the effect of imputation accuracy affected by SNV sites that fall within the BBIS regions. These four datasets were ELLIPSE OncoArray (African ancestry, 4231 cases/3953 controls on Illumina Consortium-OncoArray, abbreviated as “ONCO-AAPC”), AAPC GWAS (4822 cases/4642 controls on Illumina Human1M-Duo BeadChip, abbreviated as “AAPC1M”), California and Uganda Prostate Cancer Study (1590 cases/1048 controls on Illumina H3Africa consortium array, abbreviated as “AAPC-H3”) and ELLIPSE OncoArray (Hispanic Ethnicity, 1192 cases/1052 controls on Illumina Consortium-OncoArray, abbreviated as “ONCO-LAPC”). Details of the study description, data processing and quality control filtering, and models for association testing can be found in previous publication[13]. Briefly, association with prostate cancer risk was estimated in each dataset using logistic regression adjusting for sub-study, age and top 10 principal components. Per-allele odds ratios and standard errors from three AAPC association results were meta-analyzed using fixed-effect inverse-variance weighting.

### Imputation via TOPMed imputation server

Imputation-ready GWAS datasets in either GRCh37 or GRCh38 genome build were submitted to the TOPMed imputation server (https://imputation.biodatacatalyst.nhlbi.nih.gov/) for QC and imputation. Imputation pipeline used was michigan-imputationserver-1.5.7, imputation software was minimac4-1.0.2 and phasing software was eagle-2.4.

### *triple*-*liftOver* script

*triple*-*liftOver* is a PERL script which takes an input file in PLINK bim format and converts the genomic coordinate for three consecutive bases at each chromosomal position between genome builds using UCSC’s tool *liftOver* (https://genome.ucsc.edu/cgi-bin/hgLiftOver). It outputs the new positions in the destination build with a category column indicating whether a variant is found in a BBIS region. Variants that cannot be lifted over (in the unmapped output of *liftOver*) or are no longer on the same chromosome are excluded from the output file. For a focal SNV to qualify as an inverted site, the heuristic requires either the succeeding base in the old build to become the preceding base in the new build or the preceding base in the old build to become the succeeding base in the new build.

### Merging SNVs inverted between genome builds into contiguous stretches

After the *triple*- *liftOver* was applied to each of the three GWAS PLINK .bim file to identify all sites that are inverted between genome builds, consecutive inverted SNV sites on each chromosome that are within 250kb of each other were merged into stretches (**Supplemental Table 2**). The genomic coordinates for each stretch span the first to the last SNV. A stretch defined by a single SNV flanked by two SNVs that are not found to be in regions inverted between genome builds would be assigned a length of 1 bp. Note that the span from these stretches calculated in this manner is likely to be underestimated due to potential errors near the boundaries, and would differ across array platforms due to differences in SNV content.

### Imputation Rsq by position plots for BBIS regions

*triple*-*liftOver* was applied to each GWAS PLINK bim file to identify all the inverted sites. Three approaches to imputations were conducted for each dataset (scenarios are described in detail in **Results**). Imputation results were downloaded from the server and imputation info files were inspected to assess the imputation quality for common variants in each scenario. To avoid the confusion in a situation in which the SNV falling in the BBIS regions had been dropped before the imputation (thus has no impact on the imputation result), the inverted sites identified by *triple*-*liftOver* were compared to the server QC files first. Any variants that failed the QC check due to being monomorphic, allele mismatch and not being in the reference (thus “typed only”) in imputation scenario C (**Figure 2**) were excluded. The remaining inverted sites were then merged into regions if they are within 250KB from each other. Singleton inverted sites were excluded from these region plots (**Figure 2, Supplemental Figure 3-6**). The region was expanded for another 500Kb upstream and downstream for a better overview of the region although it may cause a few regions to overlap with each other. To improve clarity of the presentation in plots, we restricted the variants in each plot to be either the common variants (MAF >= 0.01) in imputation scenario C (**Figure 2**) or an inverted site. The variants are assigned to four categories in the following order: 1) inverted palindromic SNV site, 2) inverted non-palindromic SNV site, 3) genotyped SNV and 4) imputed variant. Due to the one bp shift caused by server *liftOver*, inverted SNV sites are imputed in scenario A but genotyped in scenario B and C.

### Trait-associated SNVs

The GRCh37 build files from GWAS catalog (http://hgdownload.soe.ucsc.edu/goldenPath/hg19/database/gwasCatalog.txt.gz) and the Global Biobank Engine (https://biobankengine.stanford.edu/downloads) were used to identify the inverted sites for trait-association SNVs. For GWAS catalog file, we downloaded 331M variant-trait pairs and examined 162,001 unique single nucleotide chromosomal locations on Chr1 to 22, X and Y (chromEnd – chromStart = 1) with a unique variant name. For the Global Biobank Engine, there were genome-wide summary statistics for ∼3800 traits, and we retained all SNVs that had P < 1×10^-8^ for any trait. In total, there are 2,912 traits with at least 1 SNV associated with P < 1×10^-8^, for a total of ∼2.1M SNV-trait pairs and 298,556 unique chromosomal locations.

### Data Availability

The *triple*-*liftOver* script is publicly available at https://github.com/GraceSheng/triple-*liftOver*. Individual level African American and Latinos prostate cancer cases and controls data are available through dbGaP (accession number phs000306.v4.p1 and phs001391.v1.p1)

## Acknowledgement

This work was supported by research grants from the National Institute of Health (R35GM142783, to C.W.K.C.; R01CA257328, U19CA214253 and U01CA164973 to C.H.; K99CA246076 to L.K.). Computation for this work was supported by USC’s Center for Advanced Research Computing (CARC; https://carc.usc.edu).

