## Supplemental Figures for "Inverted genomic regions between reference genome builds in humans impact imputation accuracy and decrease the power of association testing"

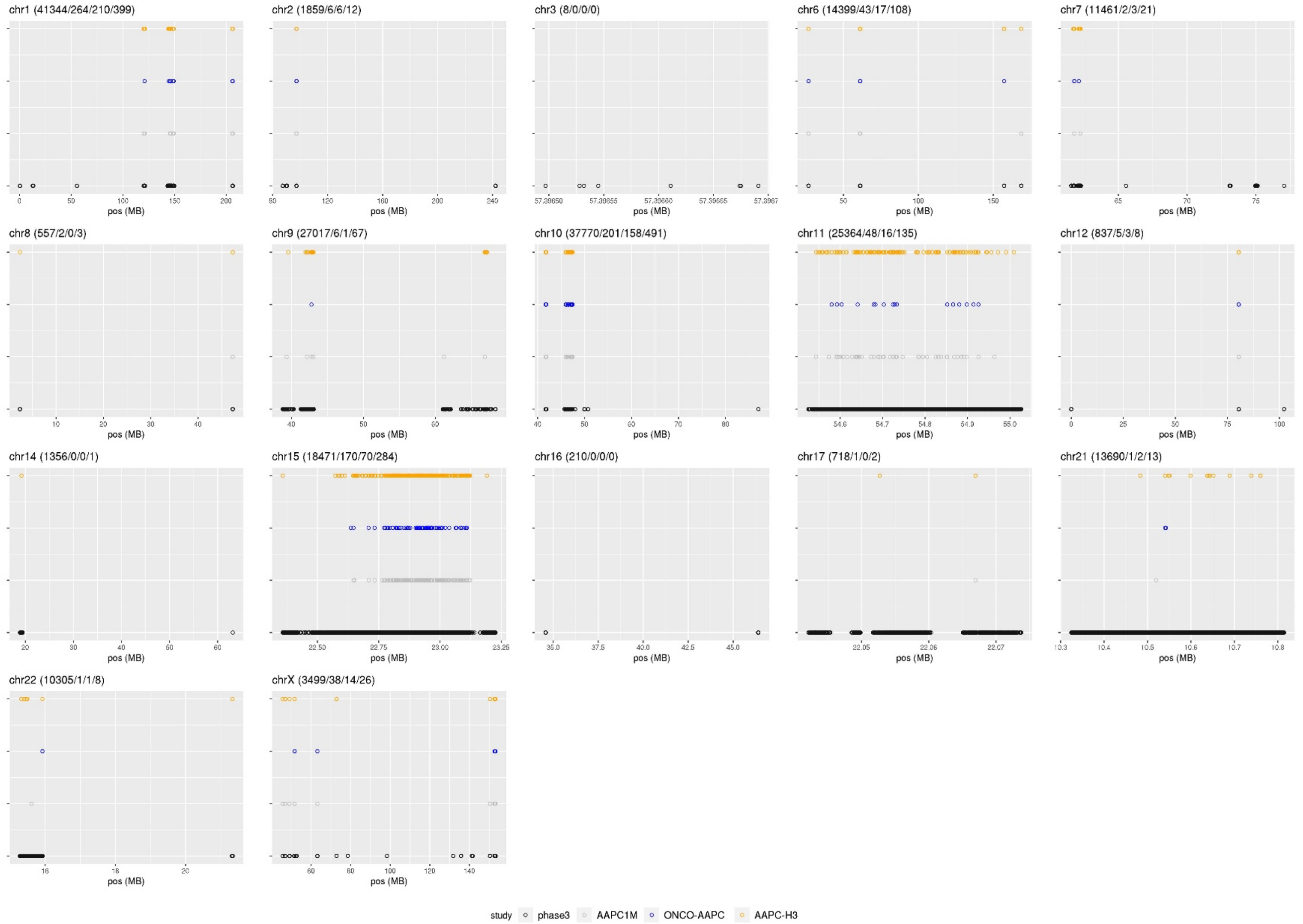

**Supplemental Figure 1.** The genomic locations (GRCh38) of BBIS-region SNVs detected by *triple-liftOver* in 1000 genomes phase3 biallelic SNVs and three GWAS arrays (AAPC1M, ONCO-AAPC and AAPC-H3). The SNV counts in the title of each plot are referring to phase3, AAPC1M, ONCO-AAPC and AAPC-H3 in this order.

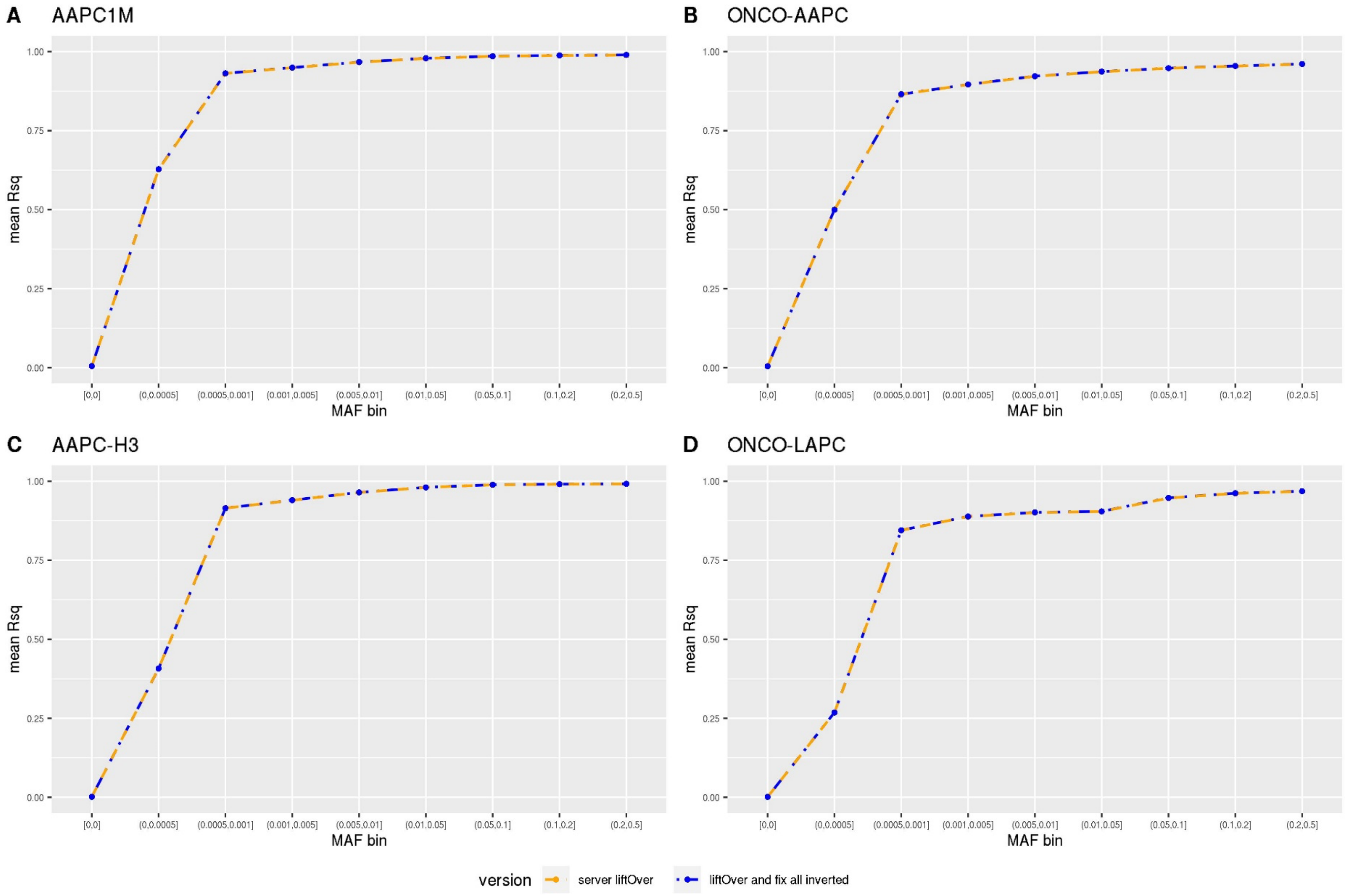


**Supplemental Figure 2**. Comparison of overall imputation quality (mean Rsq) by MAF bin between leaving GRCh38 positions and strands for BBIS-region SNVs uncorrected (server liftOver) and corrected (liftOver and fix strands for all inverted SNVs prior to submission for imputation) in four GWAS datasets.

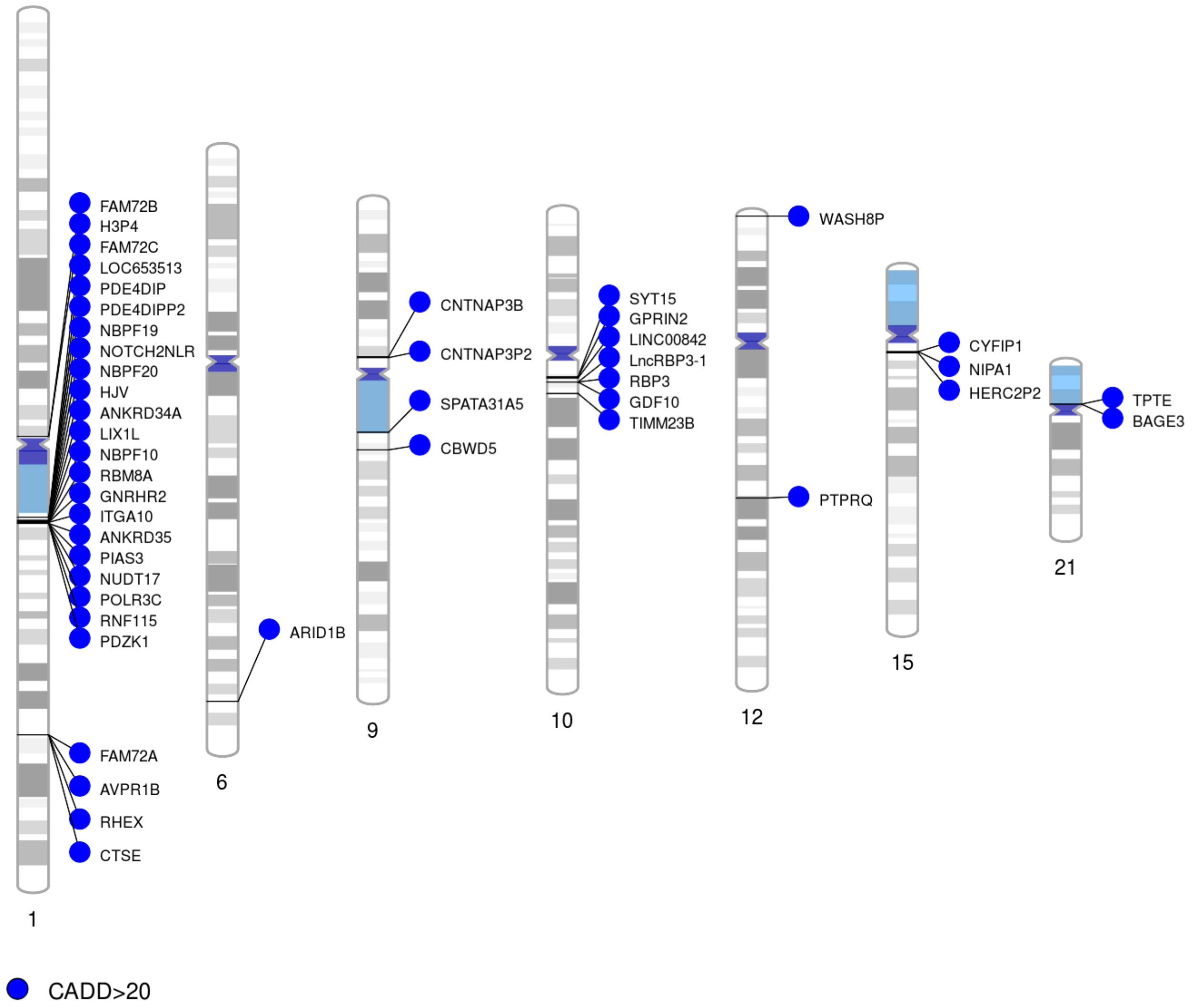


**Supplemental Figure 7:** Phenogram plot showing the BBIS regions that harbor variants with CADD PHRED-scales scores >20, corresponding to the top 1% most deleterious substitutions genome wide.
