## Supplemental Figure 3 for "Inverted genomic regions between reference genome builds in humans impact imputation accuracy and decrease the power of association testing"

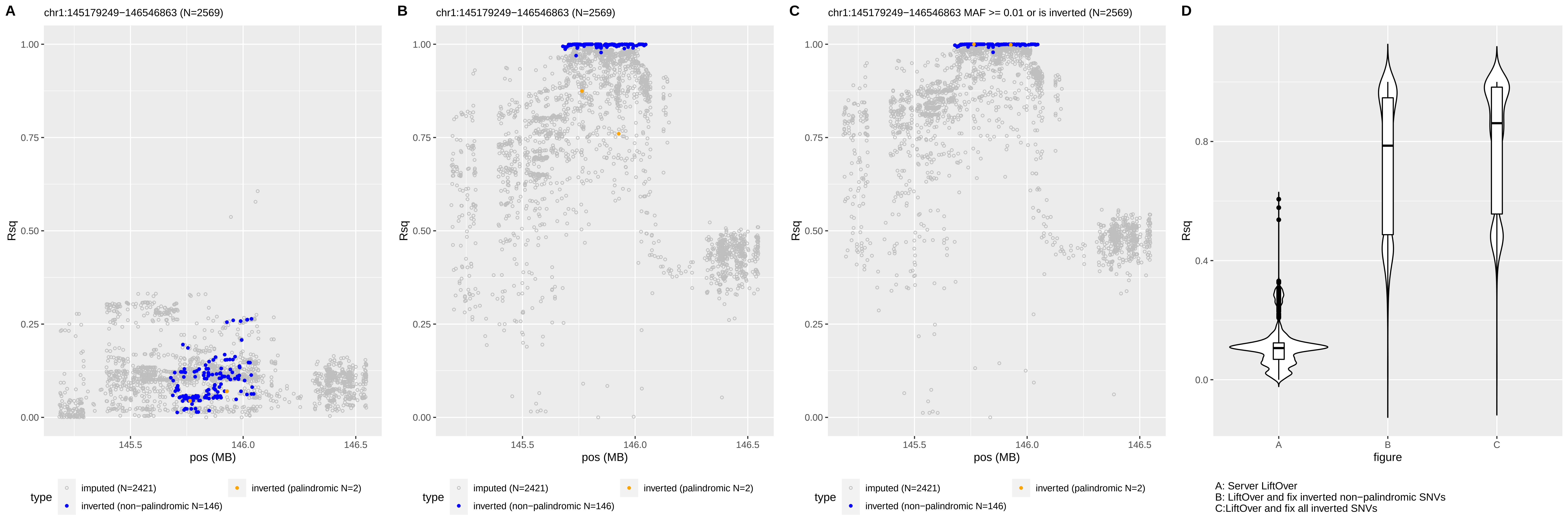
A
chr1:145179249−146546863 (N=2569)
B
chr1:145179249−146546863 (N=2569)
C
chr1:145179249−146546863 MAF >= 0.01 or is inverted (N=2569)
D
Rsq
pos (MB)
figure
type
imputed (N=2421)
inverted (palindromic N=2)
inverted (non−palindromic N=146)
A: Server LiftOver
B: LiftOver and fix inverted non−palindromic SNVs
C:LiftOver and fix all inverted SNVs

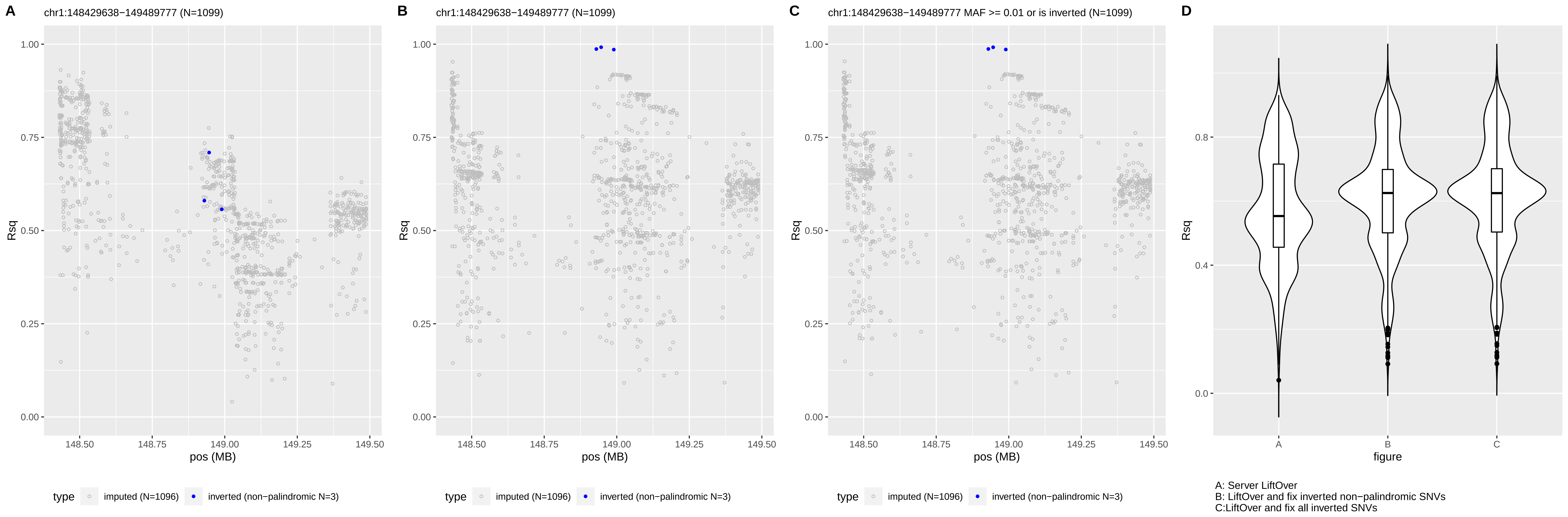
A
chr1:148429638−149489777 (N=1099)
B
chr1:148429638−149489777 (N=1099)
C
chr1:148429638−149489777 MAF >= 0.01 or is inverted (N=1099)
D
Rsq
pos (MB)
figure
type
imputed (N=1096)
inverted (non−palindromic N=3)
A: Server LiftOver
B: LiftOver and fix inverted non−palindromic SNVs
C:LiftOver and fix all inverted SNVs

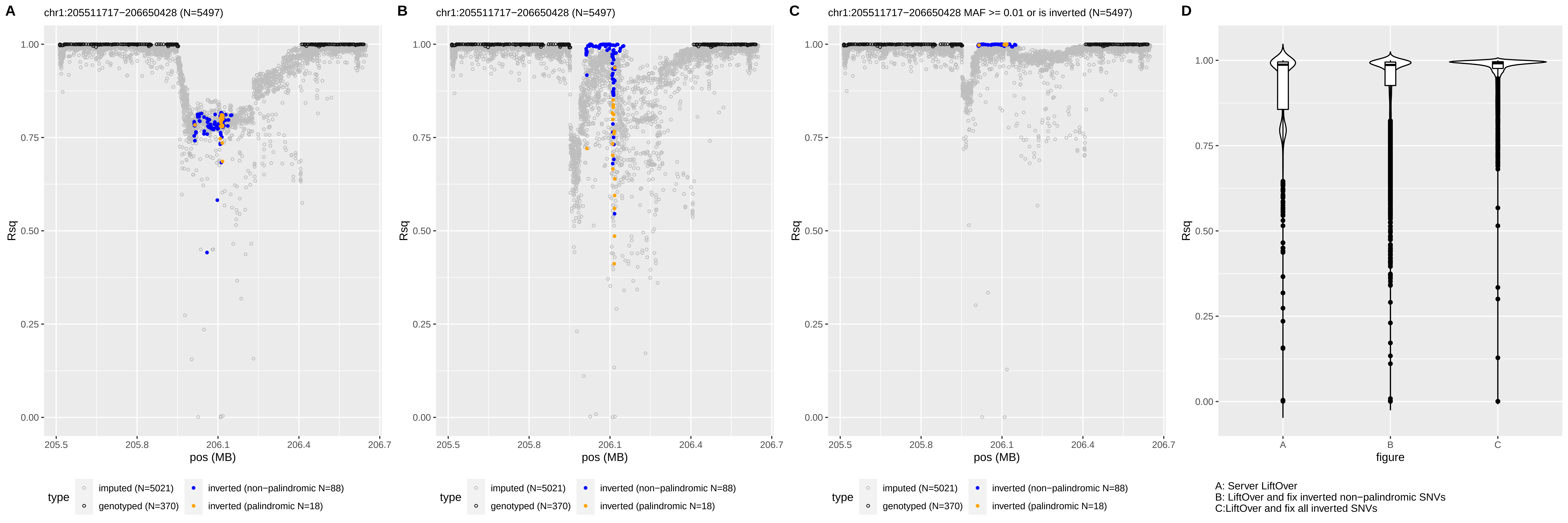

A
chr1:205511717−206650428 (N=5497)
B
chr1:205511717−206650428 (N=5497)
C
chr1:205511717−206650428 MAF >= 0.01 or is inverted (N=5497)
D
Rsq
pos (MB)
figure
type
imputed (N=5021)
genotyped (N=370)
inverted (non−palindromic N=88)
inverted (palindromic N=18)
A: Server LiftOver
B: LiftOver and fix inverted non−palindromic SNVs
C:LiftOver and fix all inverted SNVs

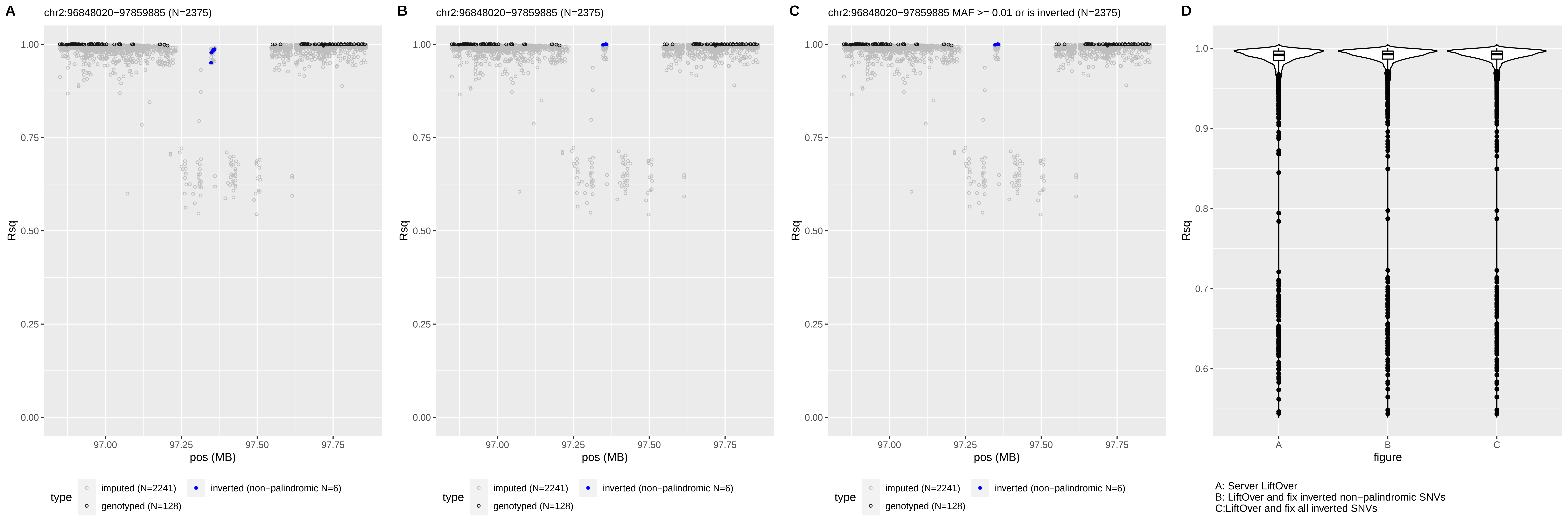
A
chr2:96848020−97859885 (N=2375)
B
chr2:96848020−97859885 (N=2375)
C
chr2:96848020−97859885 MAF >= 0.01 or is inverted (N=2375)
D
Rsq
pos (MB)
type
imputed (N=2241)
genotyped (N=128)
inverted (non−palindromic N=6)
figure
A: Server LiftOver
B: LiftOver and fix inverted non−palindromic SNVs
C:LiftOver and fix all inverted SNVs

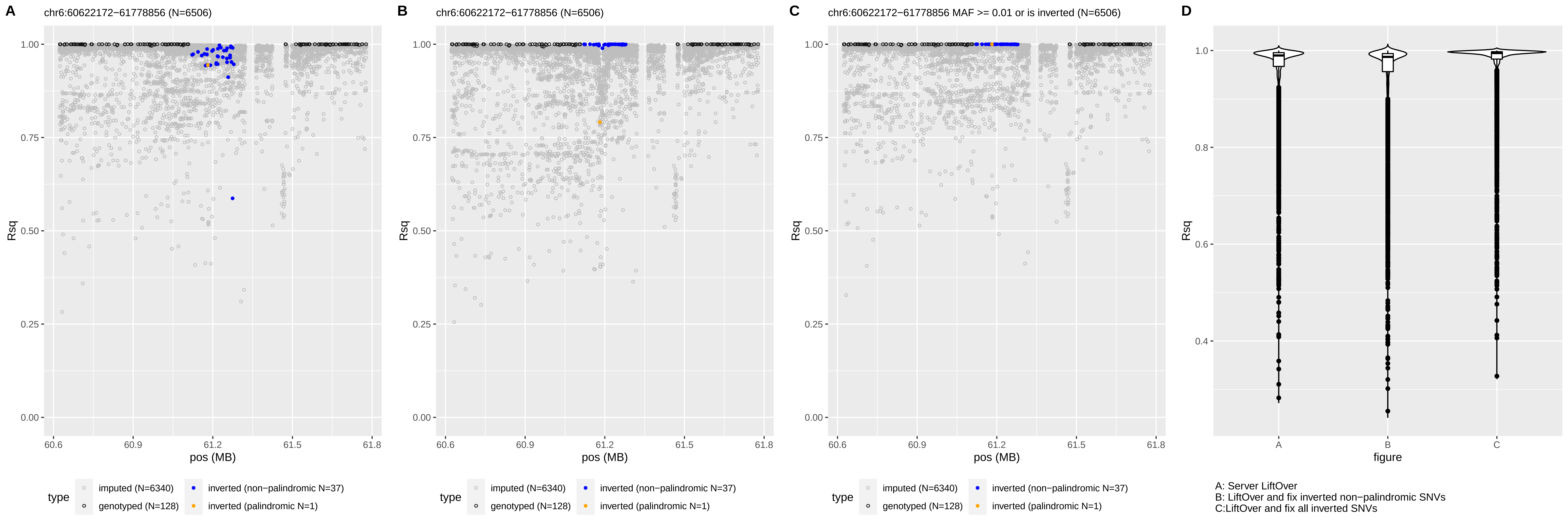
A
chr6:60622172−61778856 (N=6506)
B
chr6:60622172−61778856 (N=6506)
C
chr6:60622172−61778856 MAF >= 0.01 or is inverted (N=6506)
D
Rsq
pos (MB)
figure
type
imputed (N=6340)
genotyped (N=128)
inverted (non−palindromic N=37)
inverted (palindromic N=1)
A: Server LiftOver
B: LiftOver and fix inverted non−palindromic SNVs
C:LiftOver and fix all inverted SNVs

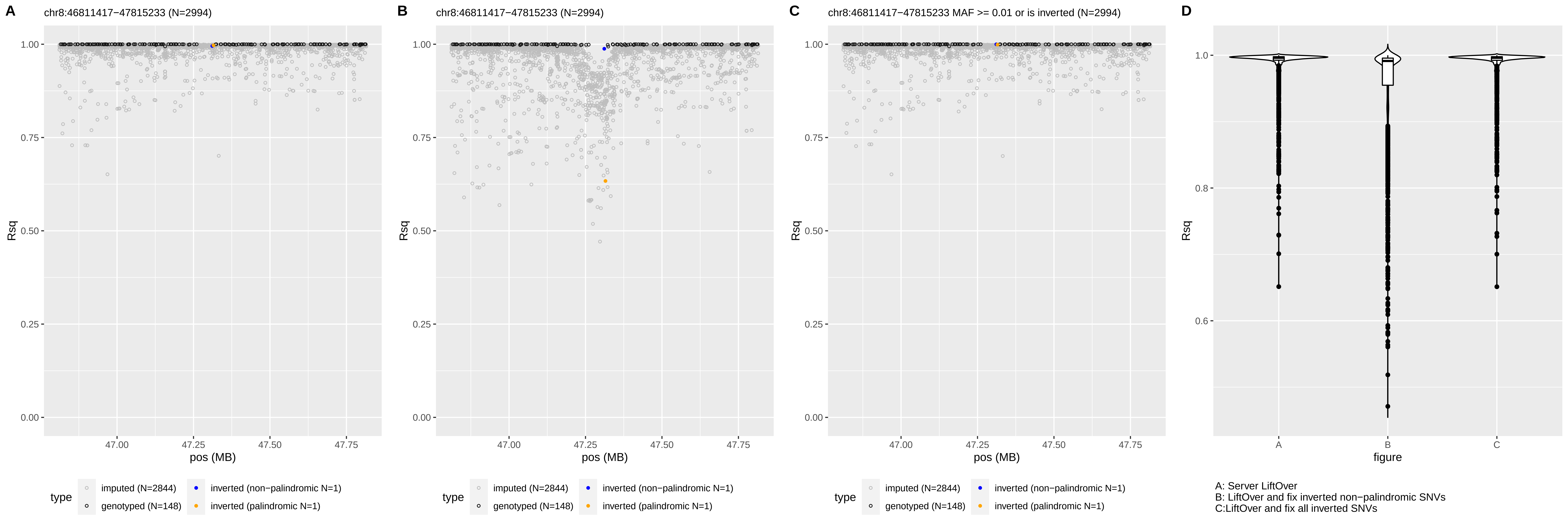
A
chr8:46811417−47815233 (N=2994)
B
chr8:46811417−47815233 (N=2994)
C
chr8:46811417−47815233 MAF >= 0.01 or is inverted (N=2994)
D
Rsq
pos (MB)
1.00
0.75
0.50
0.25
0.00
47.00
47.25
47.50
47.75
type
imputed (N=2844)
genotyped (N=148)
inverted (non−palindromic N=1)
inverted (palindromic N=1)
1.0
0.8
0.6
figure
A: Server LiftOver
B: LiftOver and fix inverted non−palindromic SNVs
C:LiftOver and fix all inverted SNVs

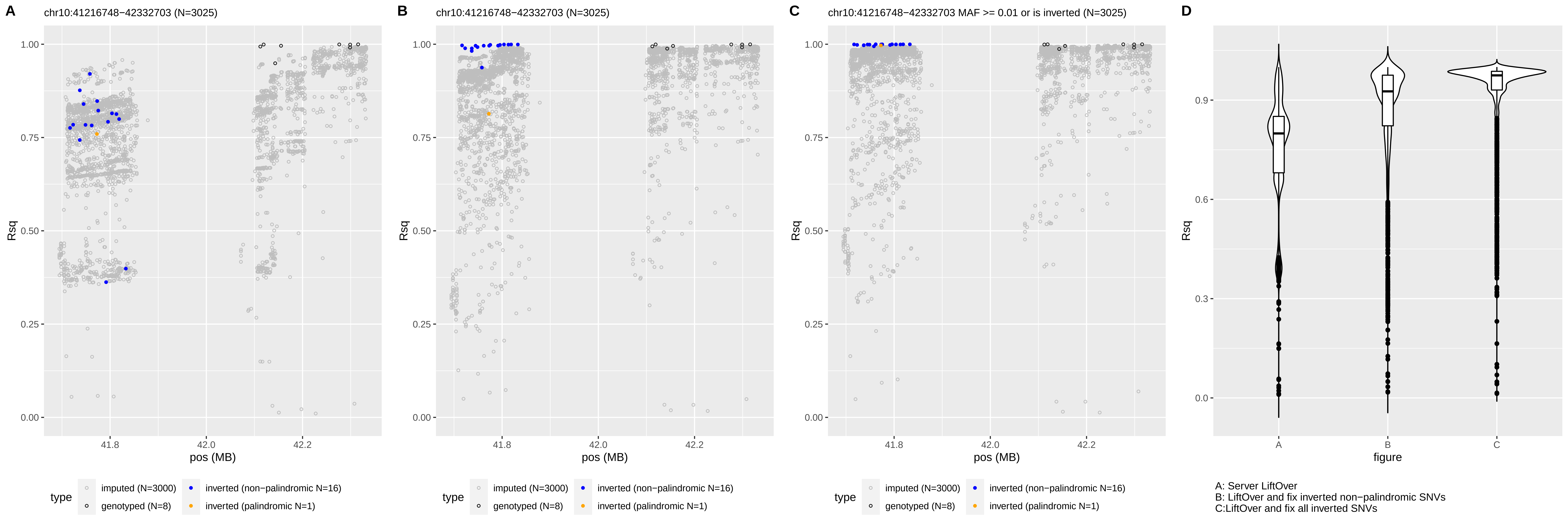
A
chr10:41216748−42332703 (N=3025)
B
chr10:41216748−42332703 (N=3025)
C
chr10:41216748−42332703 MAF >= 0.01 or is inverted (N=3025)
D
Rsq
pos (MB)
figure
type
imputed (N=3000)
genotyped (N=8)
inverted (non−palindromic N=16)
inverted (palindromic N=1)
A: Server LiftOver
B: LiftOver and fix inverted non−palindromic SNVs
C:LiftOver and fix all inverted SNVs

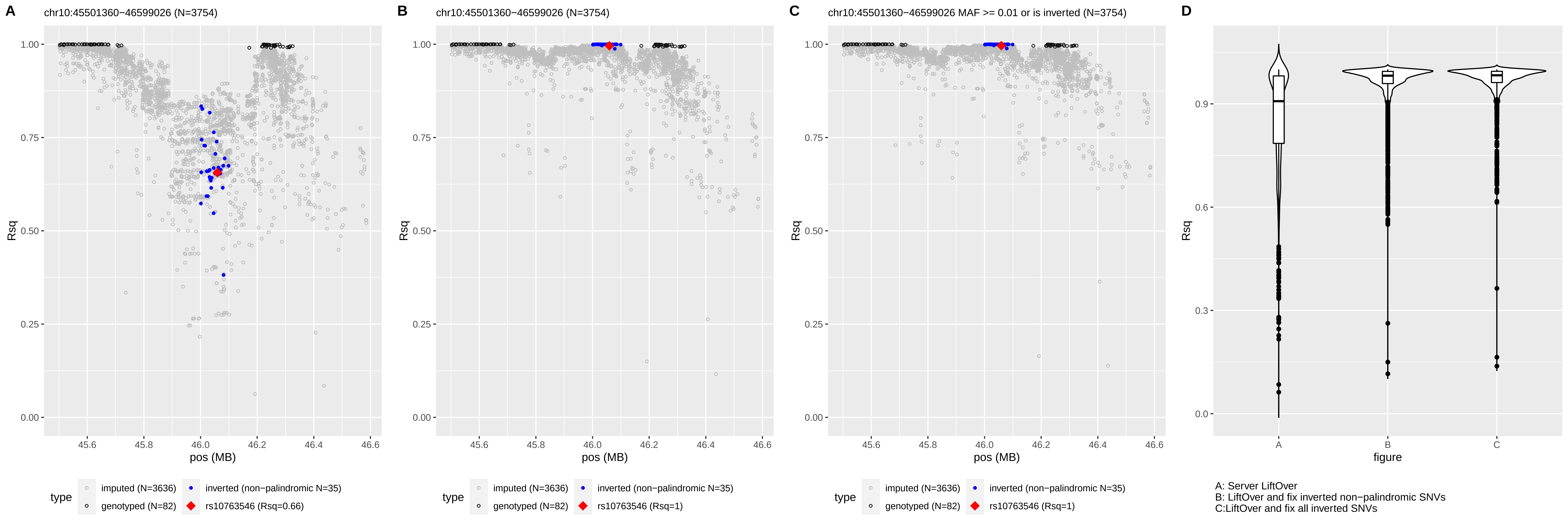

A
chr10:45501360−46599026 (N=3754)
B
chr10:45501360−46599026 (N=3754)
C
chr10:45501360−46599026 MAF >= 0.01 or is inverted (N=3754)
D
Rsq
pos (MB)
type
imputed (N=3636)
genotyped (N=82)
inverted (non−palindromic N=35)
rs10763546 (Rsq=0.66)
rs10763546 (Rsq=1)
figure
A: Server LiftOver
B: LiftOver and fix inverted non−palindromic SNVs
C:LiftOver and fix all inverted SNVs

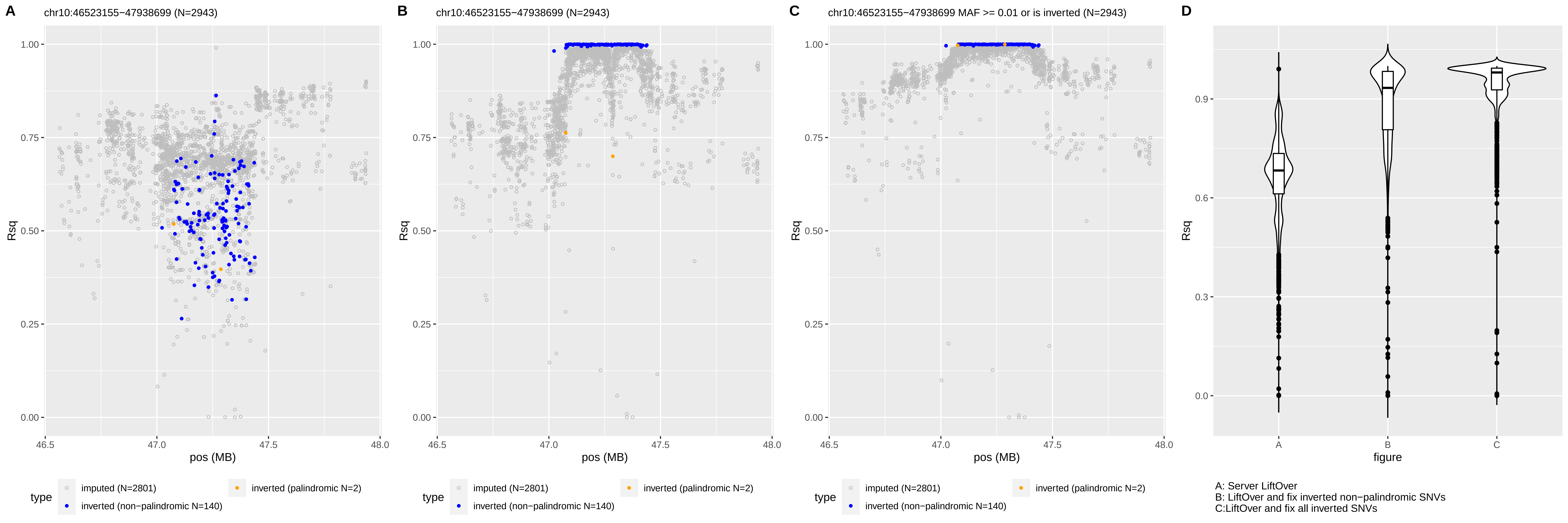

A
chr10:46523155−47938699 (N=2943)
B
chr10:46523155−47938699 (N=2943)
C
chr10:46523155−47938699 MAF >= 0.01 or is inverted (N=2943)
D
Rsq
pos (MB)
figure
type
imputed (N=2801)
inverted (non−palindromic N=140)
inverted (palindromic N=2)
A: Server LiftOver
B: LiftOver and fix inverted non−palindromic SNVs
C:LiftOver and fix all inverted SNVs

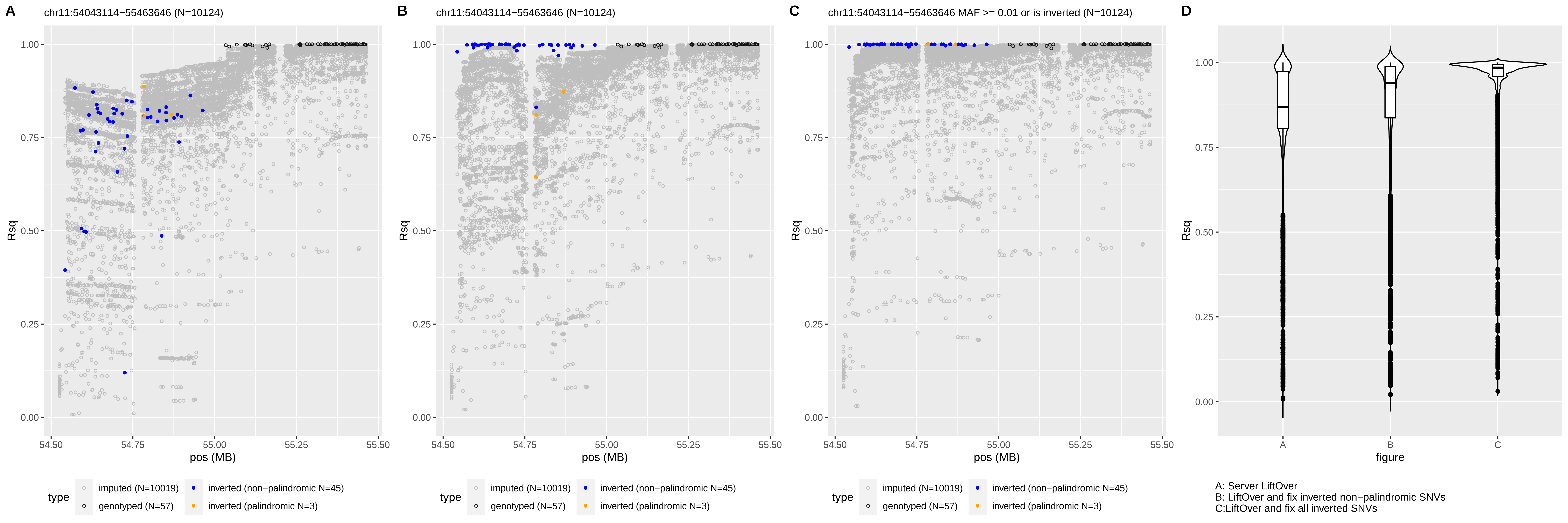
A
chr11:54043114−55463646 (N=10124)
B
chr11:54043114−55463646 (N=10124)
C
chr11:54043114−55463646 MAF >= 0.01 or is inverted (N=10124)
D
Rsq
pos (MB)
type
imputed (N=10019)
genotyped (N=57)
inverted (non−palindromic N=45)
inverted (palindromic N=3)
figure
A: Server LiftOver
B: LiftOver and fix inverted non−palindromic SNVs
C:LiftOver and fix all inverted SNVs

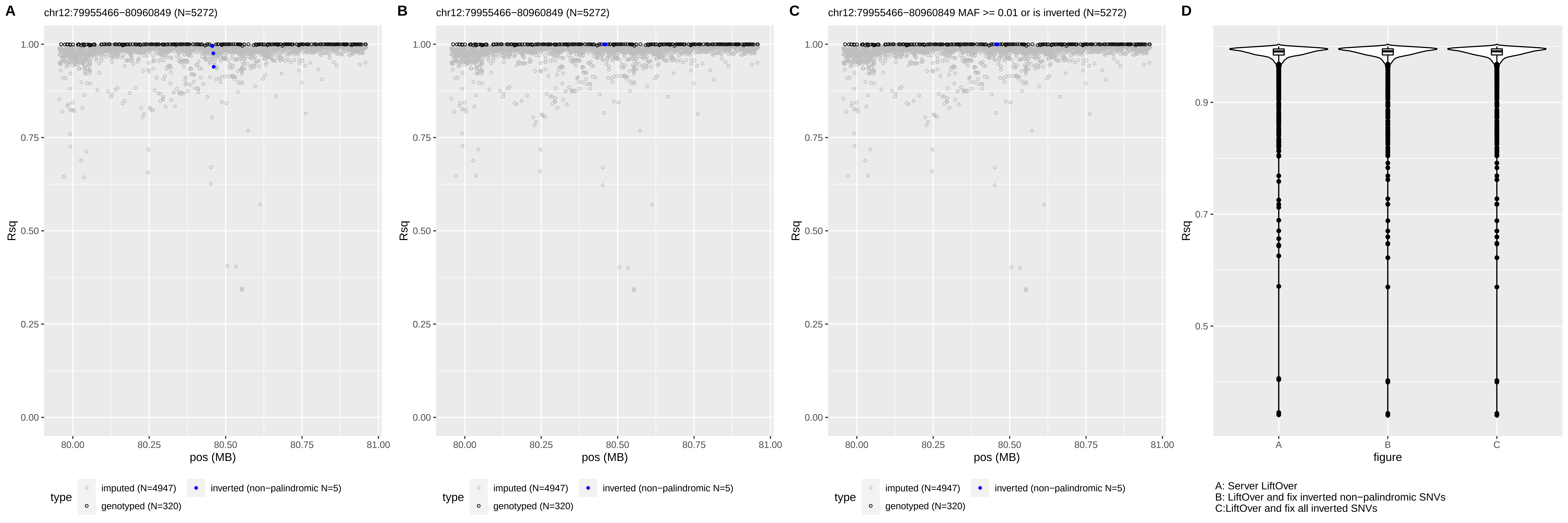
A
chr12:79955466−80960849 (N=5272)
B
chr12:79955466−80960849 (N=5272)
C
chr12:79955466−80960849 MAF >= 0.01 or is inverted (N=5272)
D
Rsq
pos (MB)
type
imputed (N=4947)
genotyped (N=320)
inverted (non−palindromic N=5)
figure
A: Server LiftOver
B: LiftOver and fix inverted non−palindromic SNVs
C:LiftOver and fix all inverted SNVs

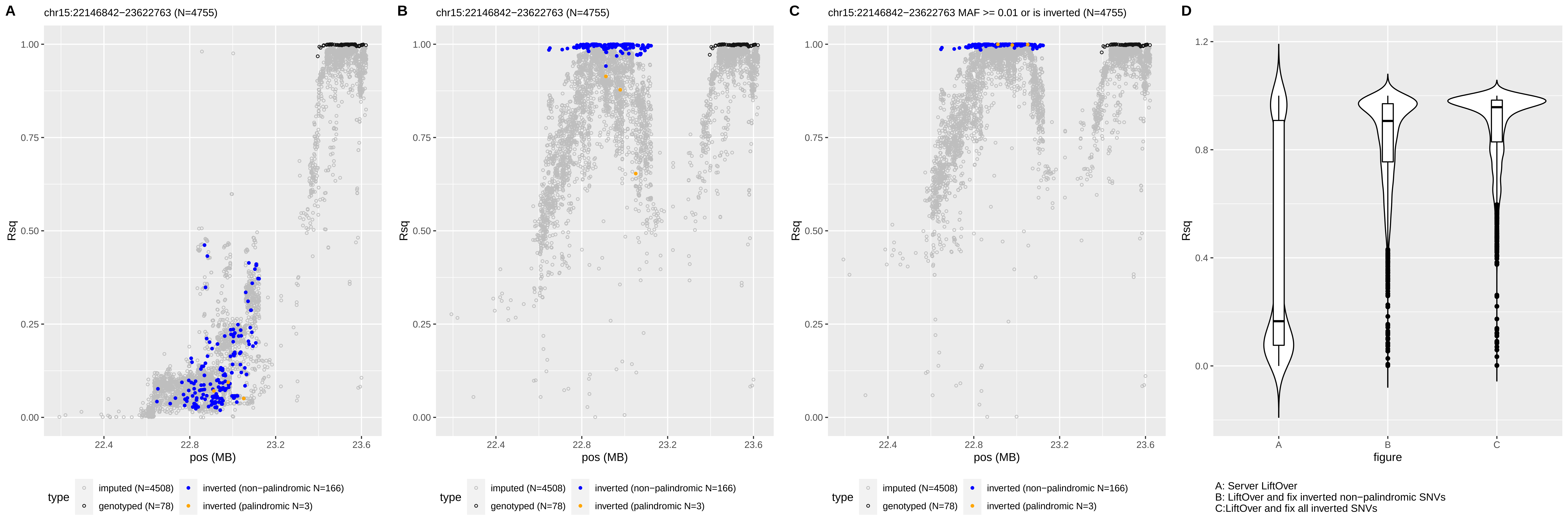

A
chr15:22146842−23622763 (N=4755)
B
chr15:22146842−23622763 (N=4755)
C
chr15:22146842−23622763 MAF >= 0.01 or is inverted (N=4755)
D
Rsq
pos (MB)
figure
type
imputed (N=4508)
genotyped (N=78)
inverted (non−palindromic N=166)
inverted (palindromic N=3)
A: Server LiftOver
B: LiftOver and fix inverted non−palindromic SNVs
C:LiftOver and fix all inverted SNVs

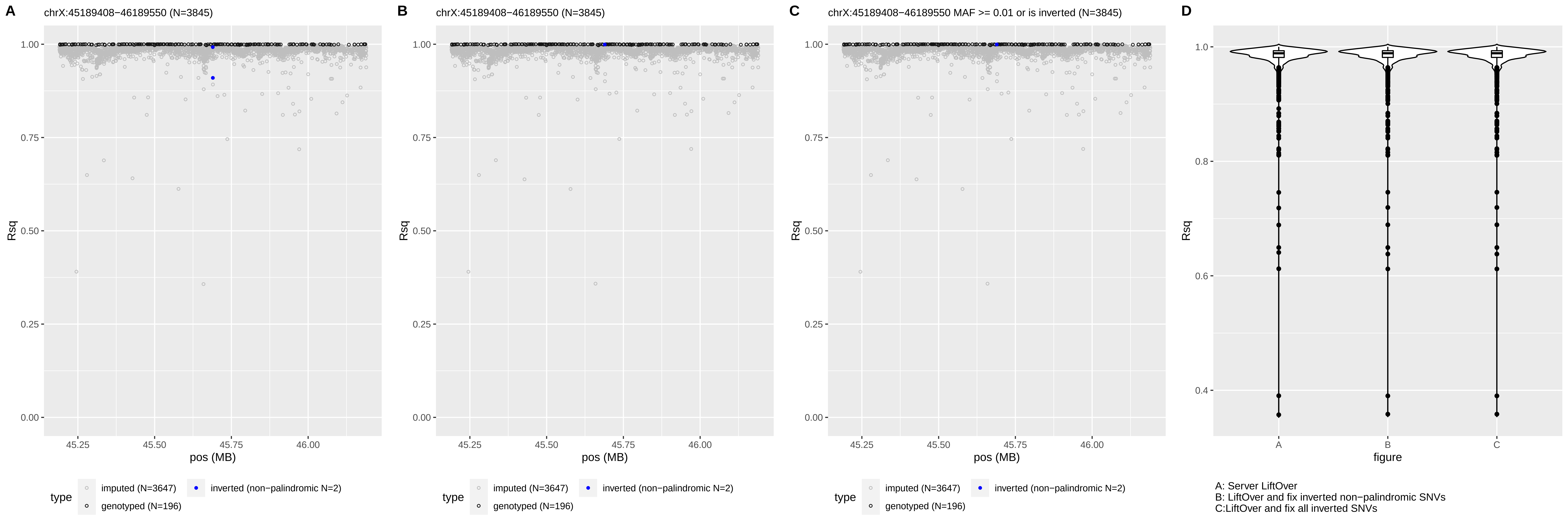

A
chrX:45189408−46189550 (N=3845)
B
chrX:45189408−46189550 (N=3845)
C
chrX:45189408−46189550 MAF >= 0.01 or is inverted (N=3845)
D
Rsq
1.00
0.75
0.50
0.25
0.00
45.25
45.50
45.75
46.00
pos (MB)
type
imputed (N=3647)
genotyped (N=196)
inverted (non−palindromic N=2)
1.0
0.8
0.6
0.4
A
B
C
figure
A: Server LiftOver
B: LiftOver and fix inverted non−palindromic SNVs
C:LiftOver and fix all inverted SNVs

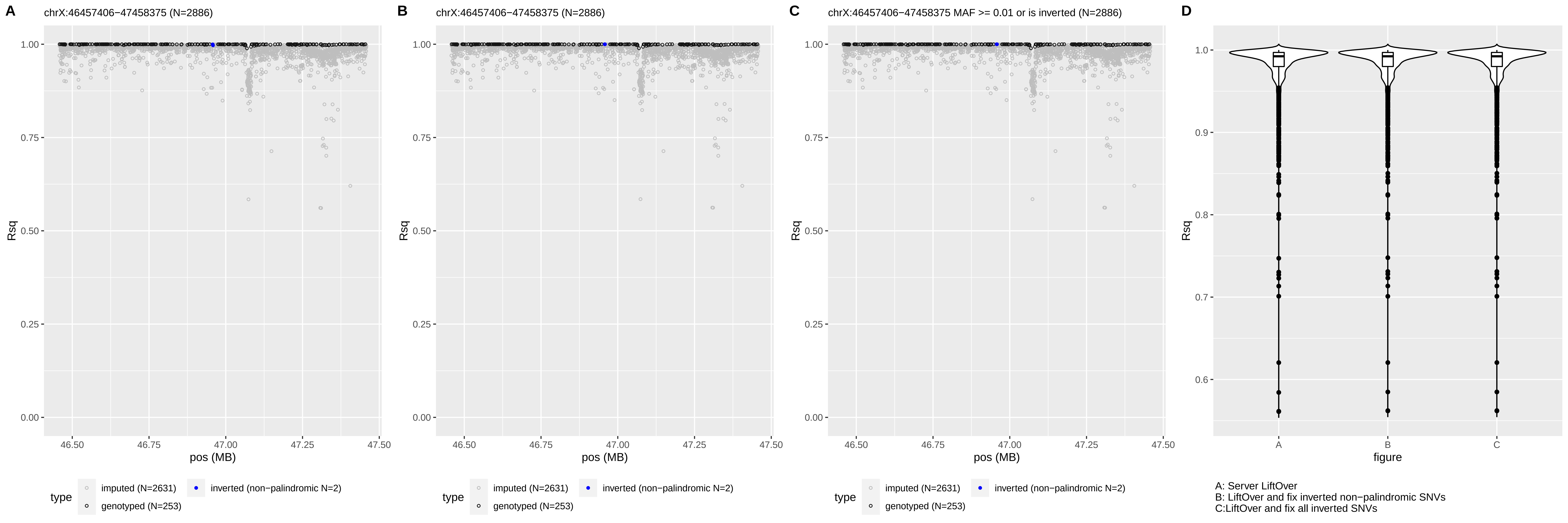

A
chrX:46457406−47458375 (N=2886)
B
chrX:46457406−47458375 (N=2886)
C
chrX:46457406−47458375 MAF >= 0.01 or is inverted (N=2886)
D
Rsq
pos (MB)
type
imputed (N=2631)
genotyped (N=253)
inverted (non−palindromic N=2)
figure
A: Server LiftOver
B: LiftOver and fix inverted non−palindromic SNVs
C:LiftOver and fix all inverted SNVs

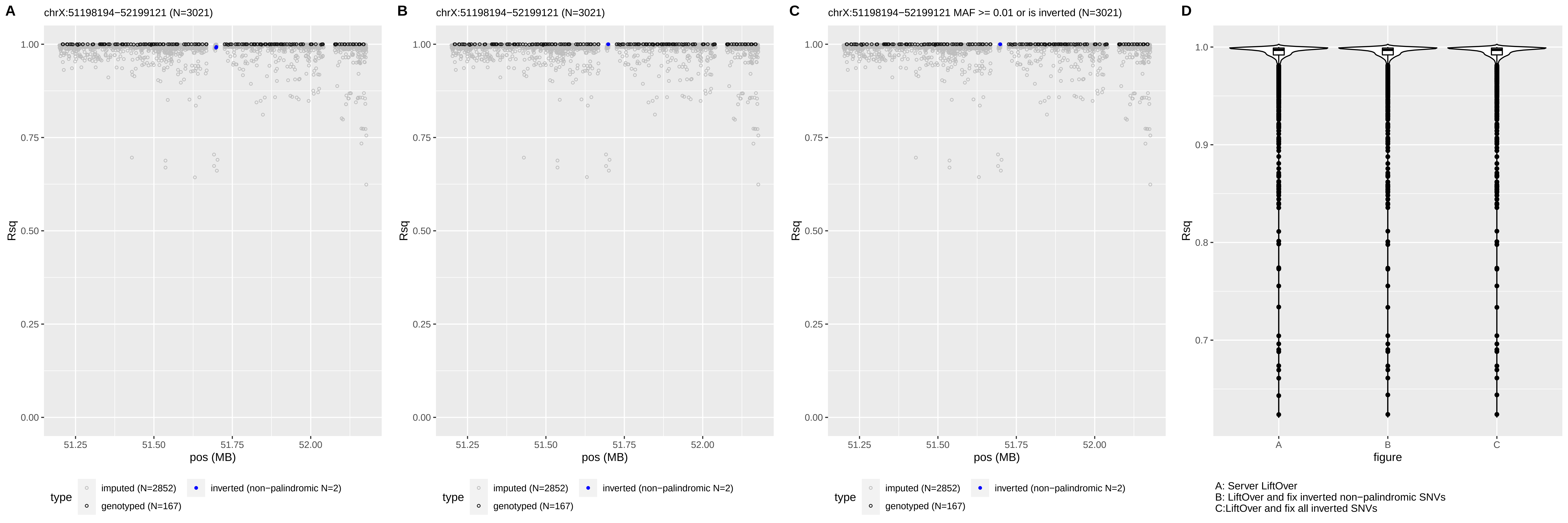

A
chrX:51198194−52199121 (N=3021)
B
chrX:51198194−52199121 (N=3021)
C
chrX:51198194−52199121 MAF >= 0.01 or is inverted (N=3021)
D
Rsq
pos (MB)
type
imputed (N=2852)
inverted (non−palindromic N=2)
genotyped (N=167)
figure
A: Server LiftOver
B: LiftOver and fix inverted non−palindromic SNVs
C:LiftOver and fix all inverted SNVs

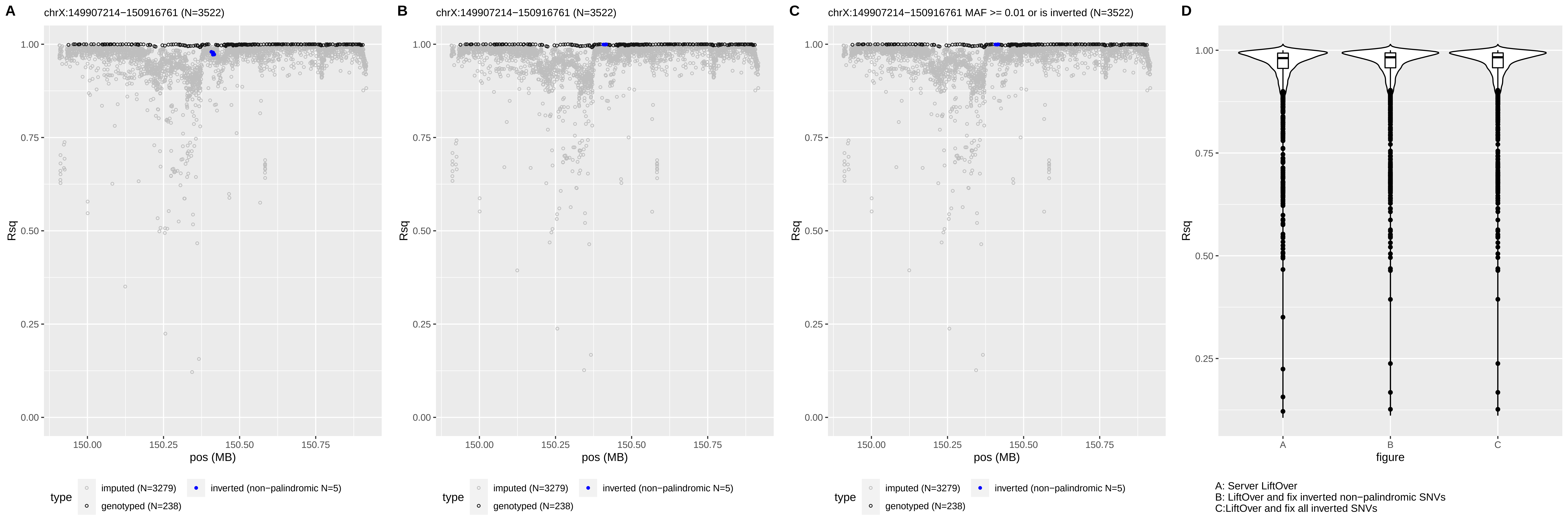
A
chrX:149907214−150916761 (N=3522)
B
chrX:149907214−150916761 (N=3522)
C
chrX:149907214−150916761 MAF >= 0.01 or is inverted (N=3522)
D
Rsq
pos (MB)
figure
type
imputed (N=3279)
genotyped (N=238)
inverted (non−palindromic N=5)
A: Server LiftOver
B: LiftOver and fix inverted non−palindromic SNVs
C:LiftOver and fix all inverted SNVs

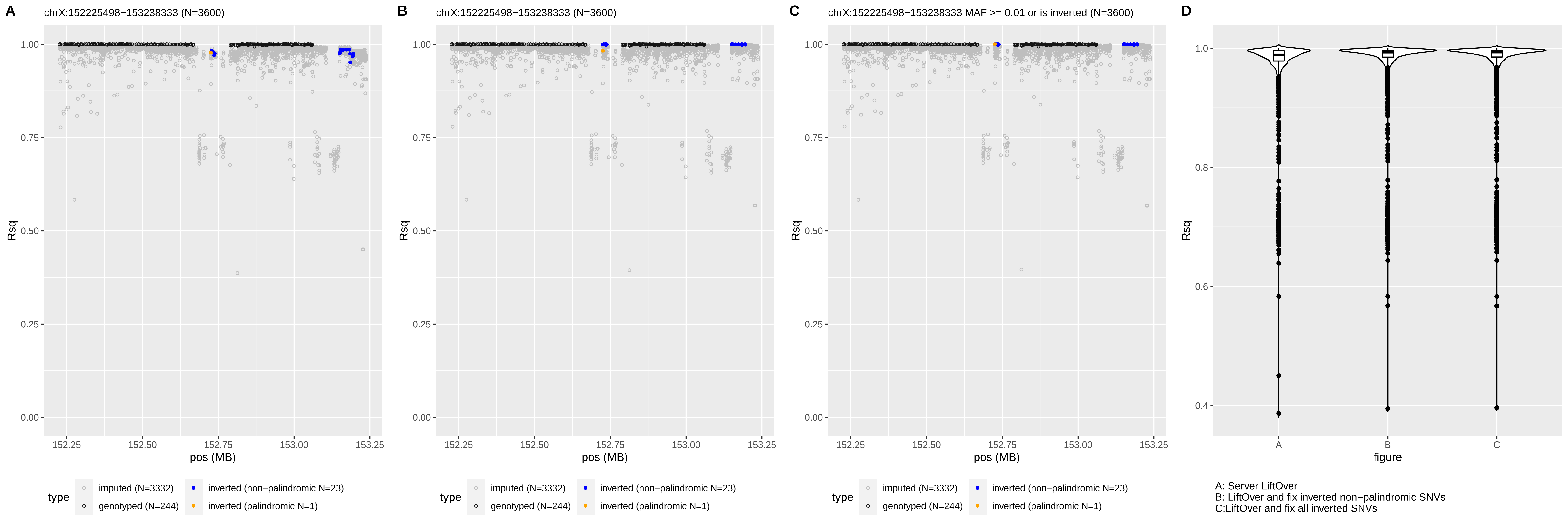

A
chrX:152225498−153238333 (N=3600)
Rsq
pos (MB)
type
imputed (N=3332)
genotyped (N=244)
inverted (non−palindromic N=23)
inverted (palindromic N=1)
B
chrX:152225498−153238333 (N=3600)
C
chrX:152225498−153238333 MAF >= 0.01 or is inverted (N=3600)
D
figure
A: Server LiftOver
B: LiftOver and fix inverted non−palindromic SNVs
C:LiftOver and fix all inverted SNVs

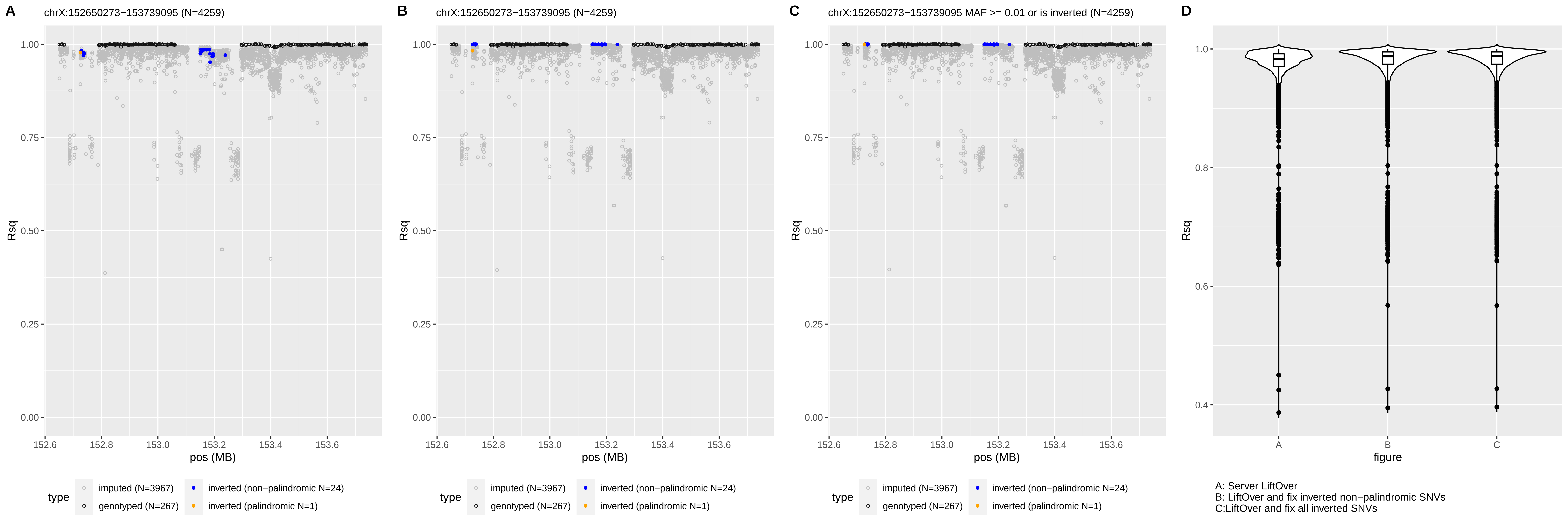
A
chrX:152650273−153739095 (N=4259)
B
chrX:152650273−153739095 (N=4259)
C
chrX:152650273−153739095 MAF >= 0.01 or is inverted (N=4259)
D
Rsq
1.00
0.75
0.50
0.25
0.00
pos (MB)
152.6
152.8
153.0
153.2
153.4
153.6
type
imputed (N=3967)
genotyped (N=267)
inverted (non−palindromic N=24)
inverted (palindromic N=1)
1.0
0.8
0.6
0.4
figure
A
B
C
A: Server LiftOver
B: LiftOver and fix inverted non−palindromic SNVs
C:LiftOver and fix all inverted SNVs
