## Supplemental Figure 4 for "Inverted genomic regions between reference genome builds in humans impact imputation accuracy and decrease the power of association testing"

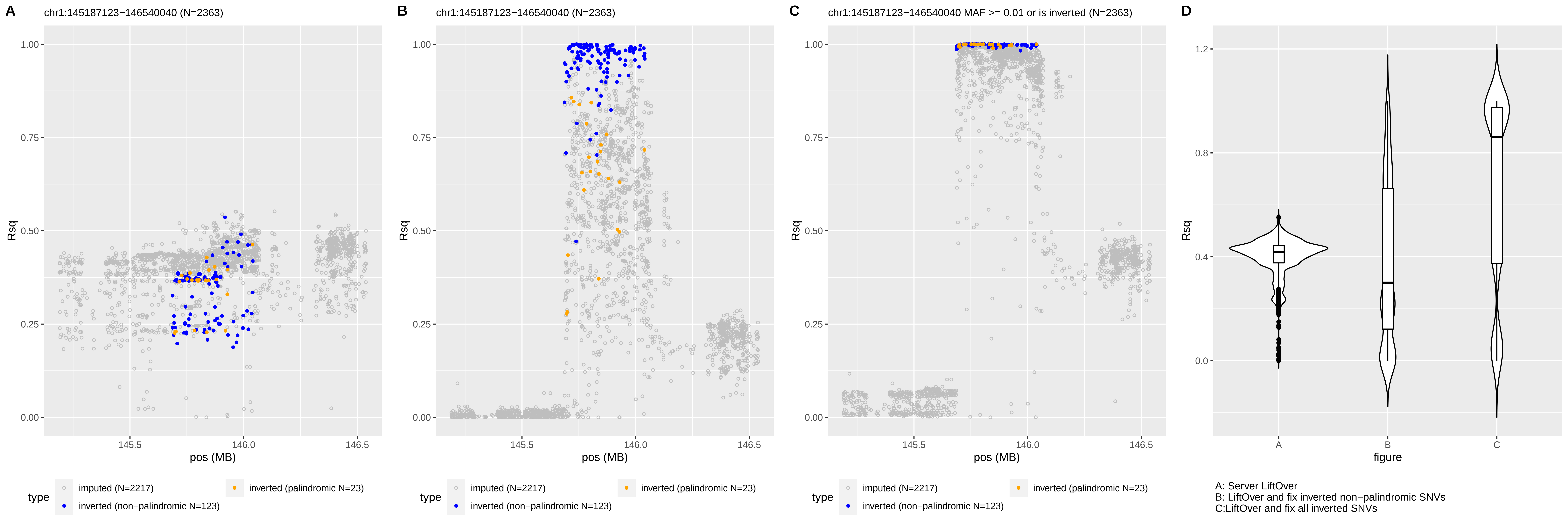

A
chr1:145187123−146540040 (N=2363)
B
chr1:145187123−146540040 (N=2363)
C
chr1:145187123−146540040 MAF >= 0.01 or is inverted (N=2363)
D
Rsq
pos (MB)
figure
type
imputed (N=2217)
inverted (non−palindromic N=123)
inverted (palindromic N=23)
A: Server LiftOver
B: LiftOver and fix inverted non−palindromic SNVs
C:LiftOver and fix all inverted SNVs

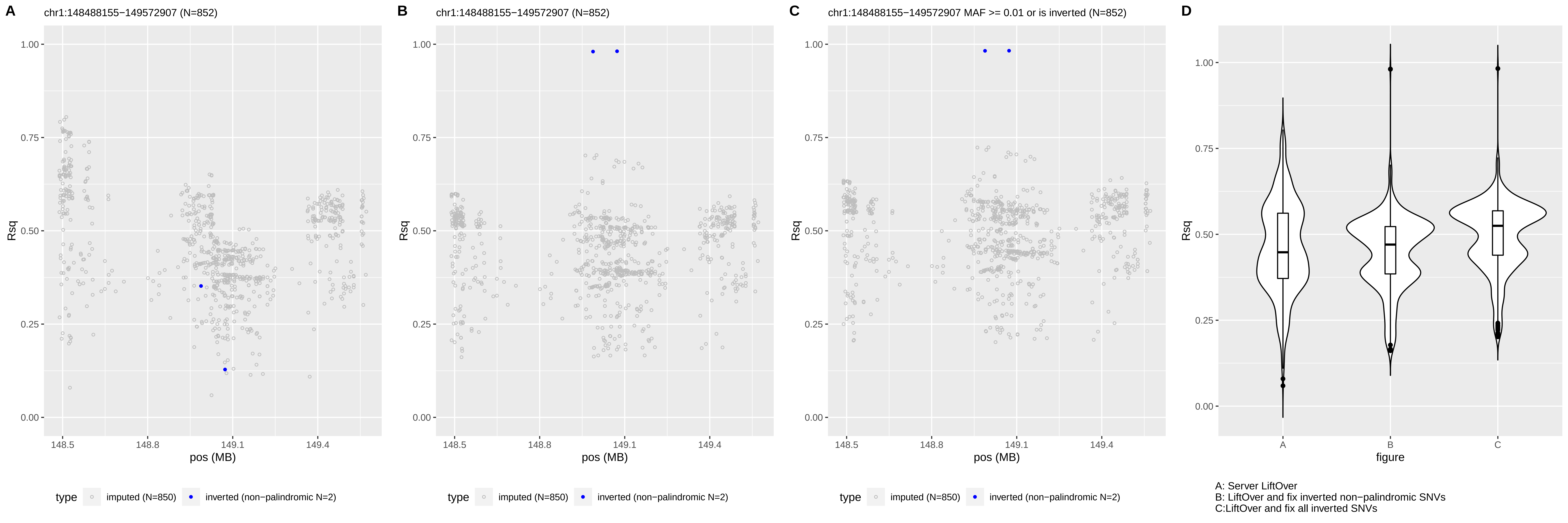
A
chr1:148488155−149572907 (N=852)
B
chr1:148488155−149572907 (N=852)
C
chr1:148488155−149572907 MAF >= 0.01 or is inverted (N=852)
D
Rsq
pos (MB)
figure
type
imputed (N=850)
inverted (non−palindromic N=2)
A: Server LiftOver
B: LiftOver and fix inverted non−palindromic SNVs
C:LiftOver and fix all inverted SNVs

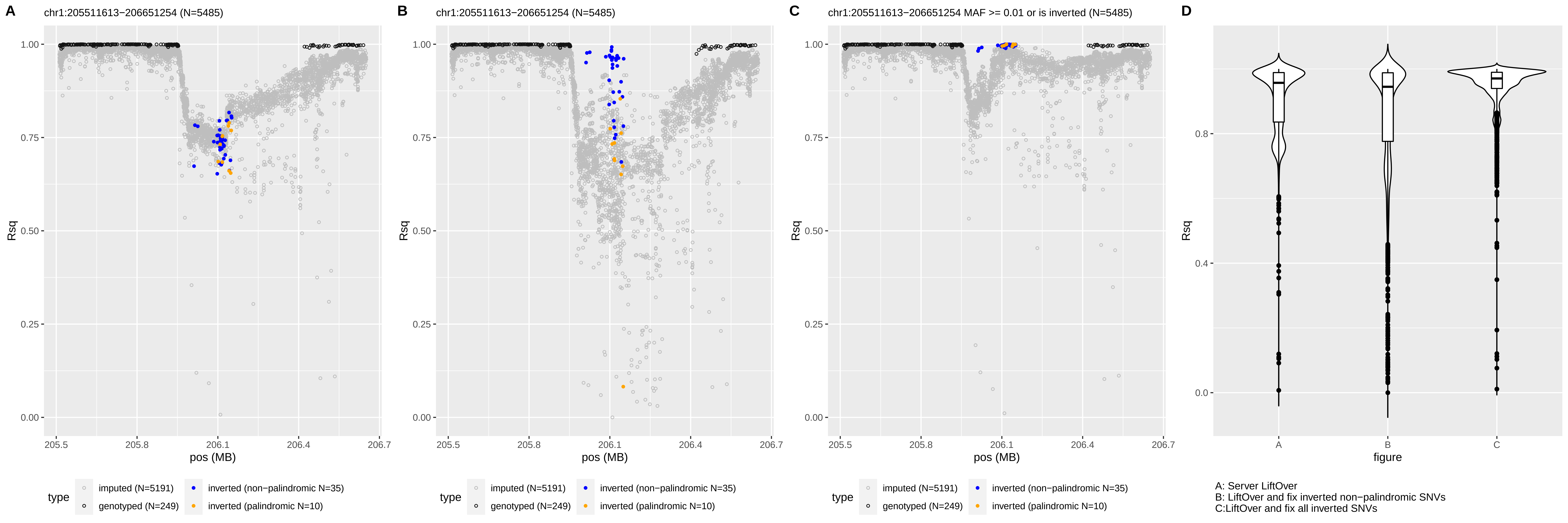
A
chr1:205511613−206651254 (N=5485)
B
chr1:205511613−206651254 (N=5485)
C
chr1:205511613−206651254 MAF >= 0.01 or is inverted (N=5485)
D
Rsq
pos (MB)
figure
type
imputed (N=5191)
genotyped (N=249)
inverted (non−palindromic N=35)
inverted (palindromic N=10)
A: Server LiftOver
B: LiftOver and fix inverted non−palindromic SNVs
C:LiftOver and fix all inverted SNVs

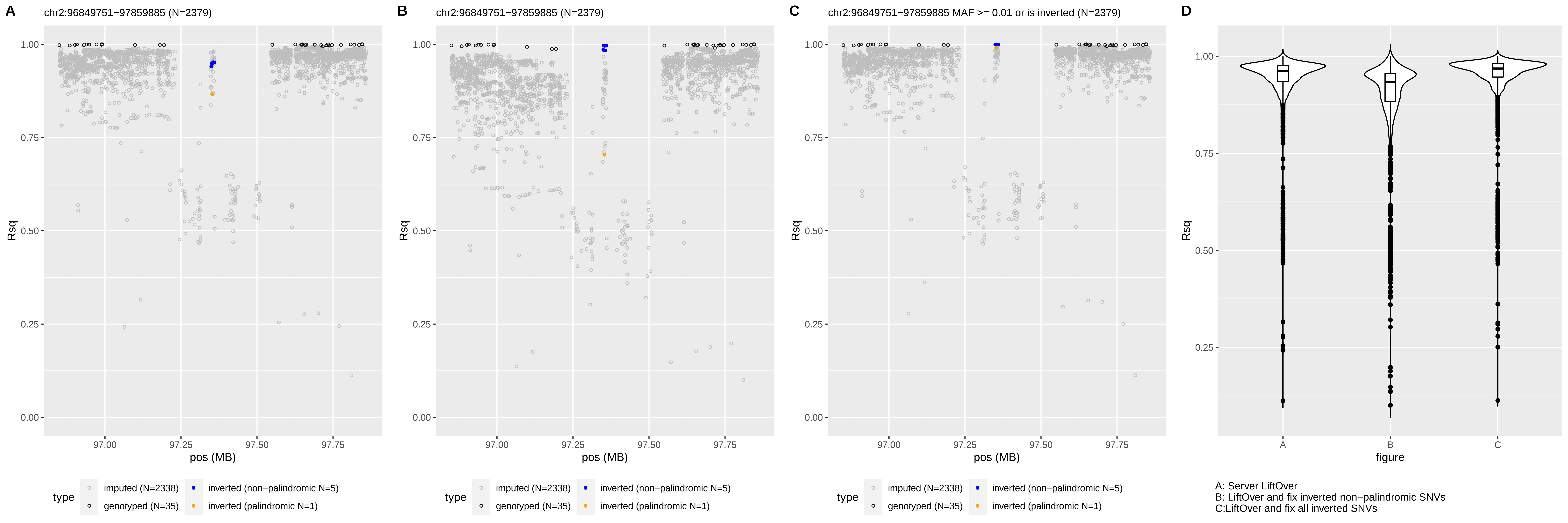
A
chr2:96849751−97859885 (N=2379)
B
chr2:96849751−97859885 (N=2379)
C
chr2:96849751−97859885 MAF >= 0.01 or is inverted (N=2379)
D
Rsq
pos (MB)
type
imputed (N=2338)
genotyped (N=35)
inverted (non−palindromic N=5)
inverted (palindromic N=1)
figure
A: Server LiftOver
B: LiftOver and fix inverted non−palindromic SNVs
C:LiftOver and fix all inverted SNVs

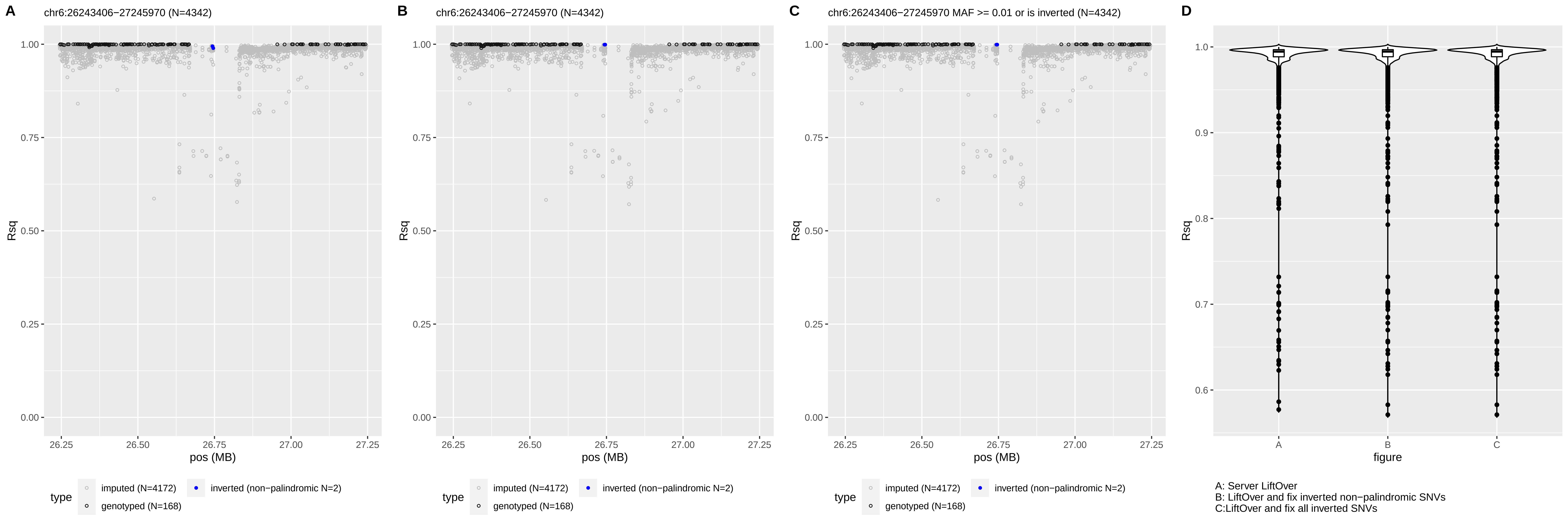
A
chr6:26243406−27245970 (N=4342)
B
chr6:26243406−27245970 (N=4342)
C
chr6:26243406−27245970 MAF >= 0.01 or is inverted (N=4342)
D
Rsq
pos (MB)
type
imputed (N=4172)
genotyped (N=168)
inverted (non−palindromic N=2)
figure
A: Server LiftOver
B: LiftOver and fix inverted non−palindromic SNVs
C:LiftOver and fix all inverted SNVs

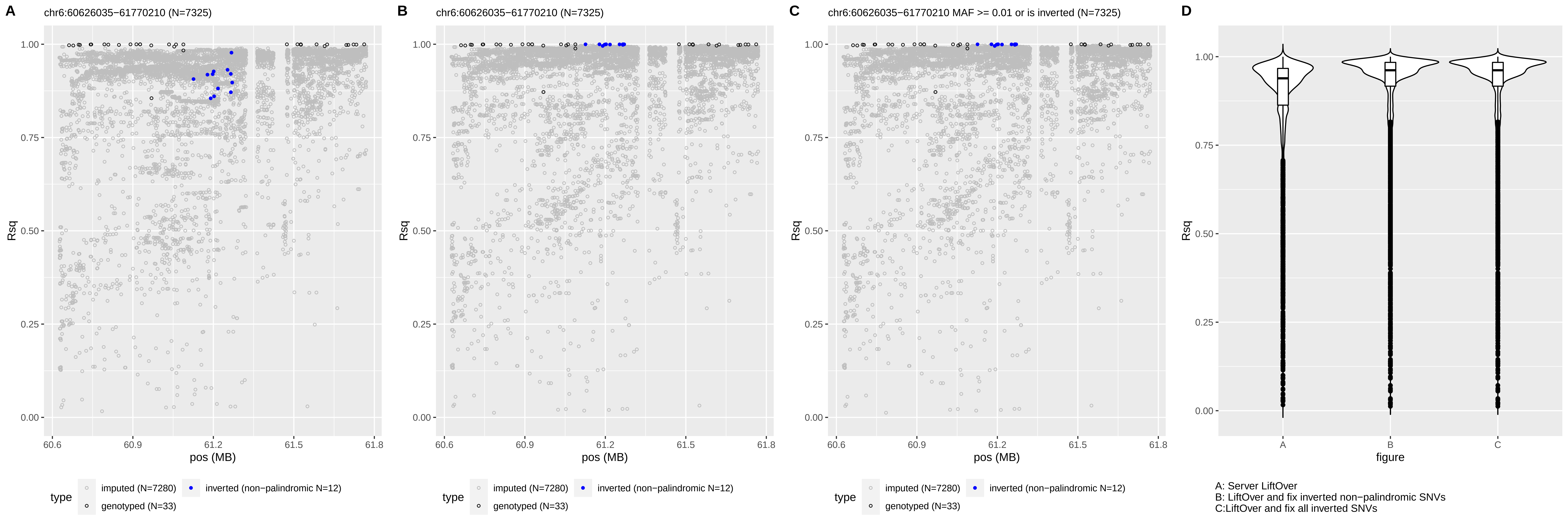
A
chr6:60626035−61770210 (N=7325)
B
chr6:60626035−61770210 (N=7325)
C
chr6:60626035−61770210 MAF >= 0.01 or is inverted (N=7325)
D
Rsq
pos (MB)
figure
type
imputed (N=7280)
inverted (non−palindromic N=12)
genotyped (N=33)
A: Server LiftOver
B: LiftOver and fix inverted non−palindromic SNVs
C:LiftOver and fix all inverted SNVs

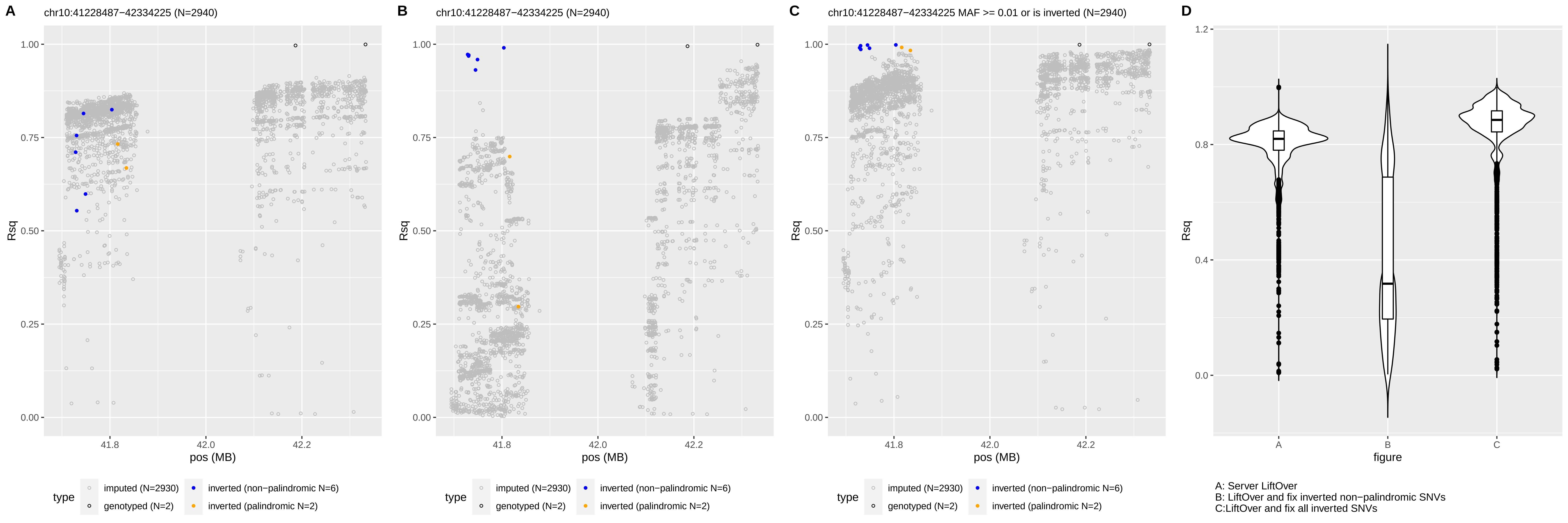
A
chr10:41228487−42334225 (N=2940)
B
chr10:41228487−42334225 (N=2940)
C
chr10:41228487−42334225 MAF >= 0.01 or is inverted (N=2940)
D
Rsq
pos (MB)
figure
type
imputed (N=2930)
genotyped (N=2)
inverted (non−palindromic N=6)
inverted (palindromic N=2)
A: Server LiftOver
B: LiftOver and fix inverted non−palindromic SNVs
C:LiftOver and fix all inverted SNVs

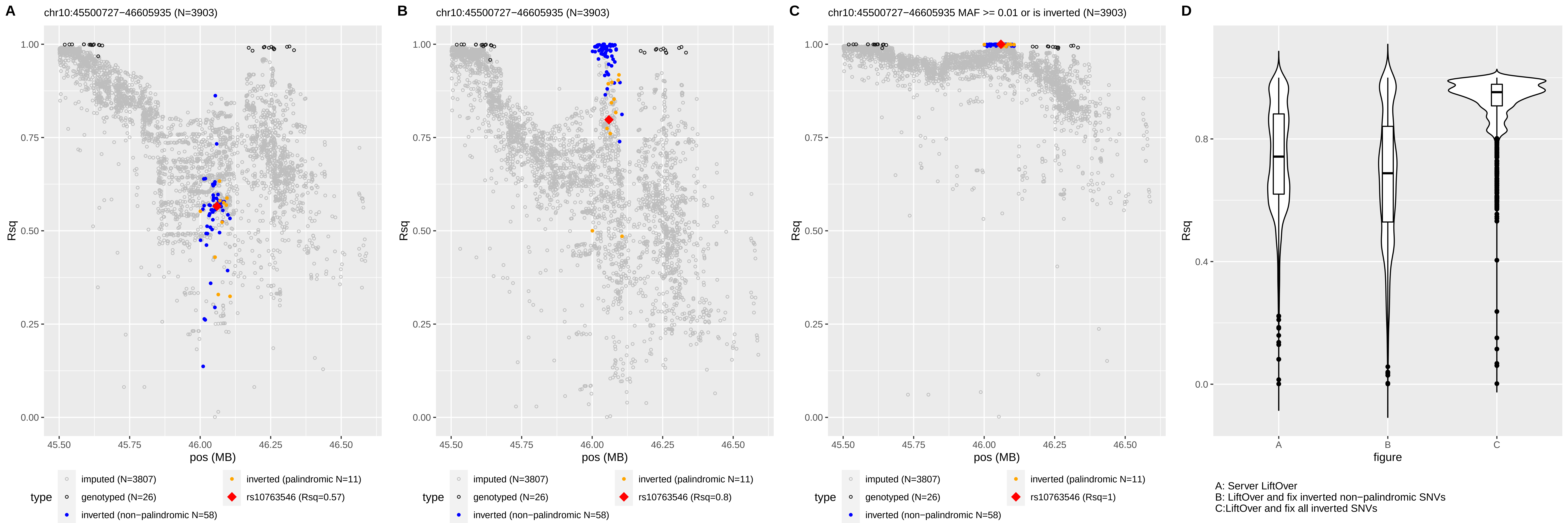
A
chr10:45500727−46605935 (N=3903)
B
chr10:45500727−46605935 (N=3903)
C
chr10:45500727−46605935 MAF >= 0.01 or is inverted (N=3903)
D
Rsq
1.00
0.75
0.50
0.25
0.00
45.50
45.75
46.00
46.25
46.50
pos (MB)
type
imputed (N=3807)
genotyped (N=26)
inverted (non−palindromic N=58)
inverted (palindromic N=11)
rs10763546 (Rsq=0.57)
rs10763546 (Rsq=0.8)
rs10763546 (Rsq=1)
0.8
0.4
0.0
A
B
C
figure
A: Server LiftOver
B: LiftOver and fix inverted non−palindromic SNVs
C:LiftOver and fix all inverted SNVs

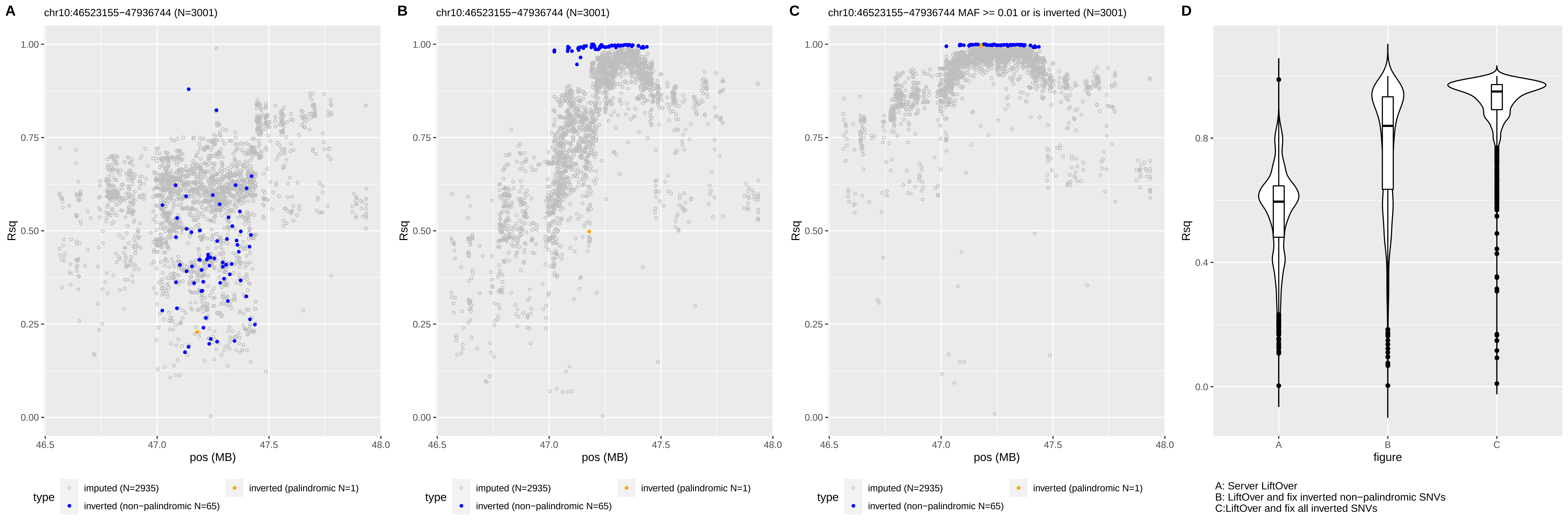
A
chr10:46523155−47936744 (N=3001)
B
chr10:46523155−47936744 (N=3001)
C
chr10:46523155−47936744 MAF >= 0.01 or is inverted (N=3001)
D
Rsq
1.00
0.75
0.50
0.25
0.00
46.5
47.0
47.5
48.0
pos (MB)
0.8
0.4
0.0
A
B
C
figure
type
imputed (N=2935)
inverted (palindromic N=1)
inverted (non−palindromic N=65)
A: Server LiftOver
B: LiftOver and fix inverted non−palindromic SNVs
C:LiftOver and fix all inverted SNVs

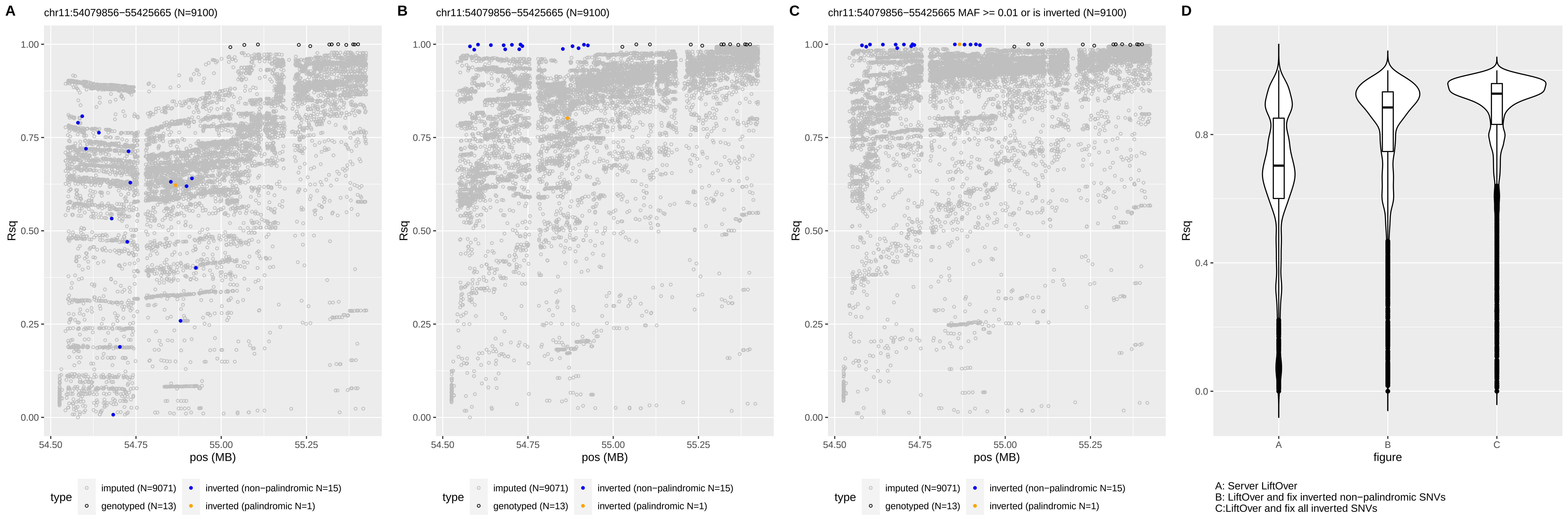
A
chr11:54079856−55425665 (N=9100)
B
chr11:54079856−55425665 (N=9100)
C
chr11:54079856−55425665 MAF >= 0.01 or is inverted (N=9100)
D
Rsq
pos (MB)
figure
type
imputed (N=9071)
genotyped (N=13)
inverted (non−palindromic N=15)
inverted (palindromic N=1)
A: Server LiftOver
B: LiftOver and fix inverted non−palindromic SNVs
C:LiftOver and fix all inverted SNVs

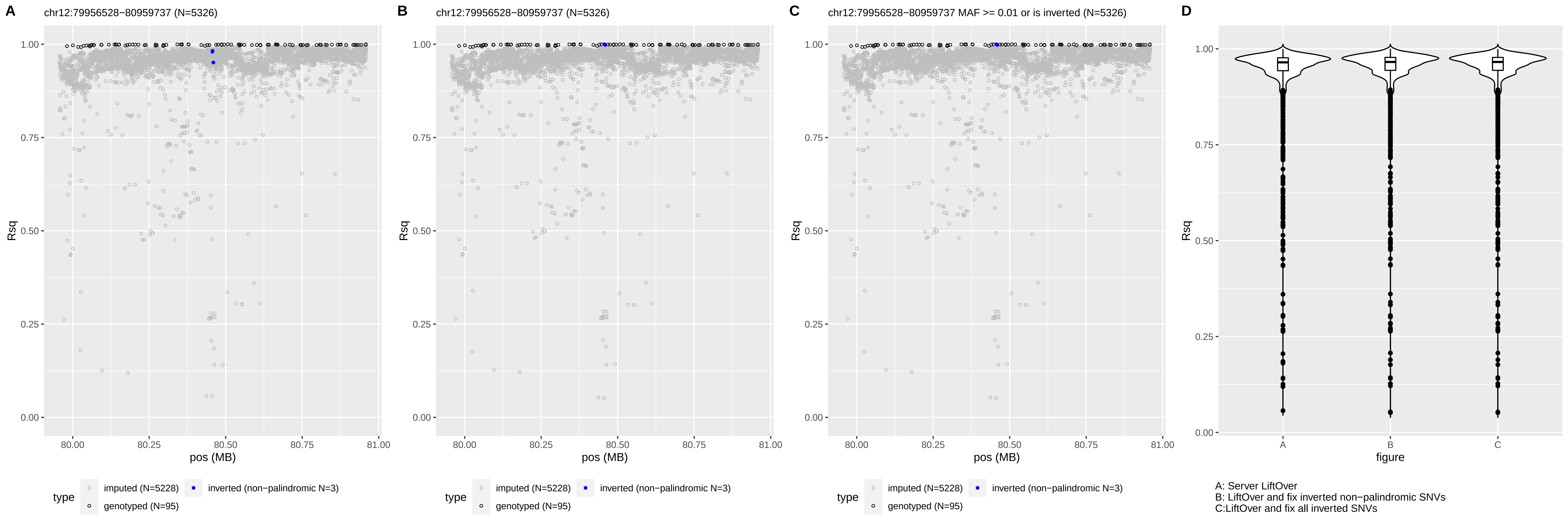

A
chr12:79956528−80959737 (N=5326)
B
chr12:79956528−80959737 (N=5326)
C
chr12:79956528−80959737 MAF >= 0.01 or is inverted (N=5326)
D
Rsq
pos (MB)
type
imputed (N=5228)
inverted (non−palindromic N=3)
genotyped (N=95)
figure
A: Server LiftOver
B: LiftOver and fix inverted non−palindromic SNVs
C:LiftOver and fix all inverted SNVs

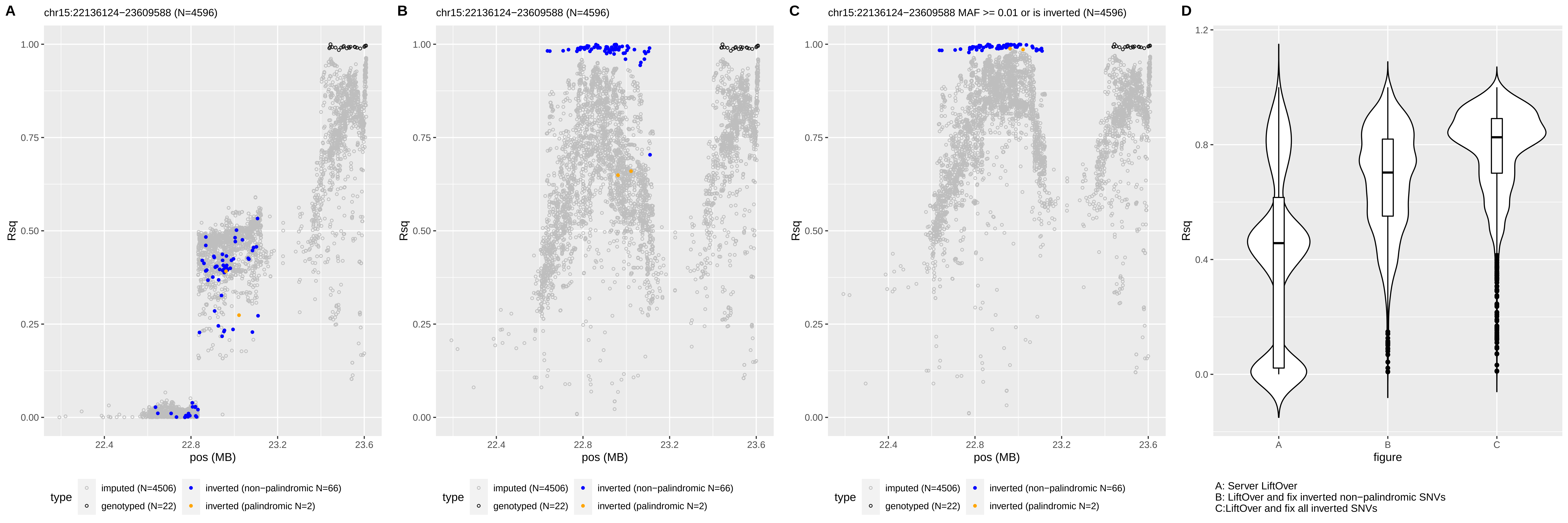
A
chr15:22136124−23609588 (N=4596)
B
chr15:22136124−23609588 (N=4596)
C
chr15:22136124−23609588 MAF >= 0.01 or is inverted (N=4596)
D
Rsq
pos (MB)
figure
type
imputed (N=4506)
genotyped (N=22)
inverted (non−palindromic N=66)
inverted (palindromic N=2)
A: Server LiftOver
B: LiftOver and fix inverted non−palindromic SNVs
C:LiftOver and fix all inverted SNVs

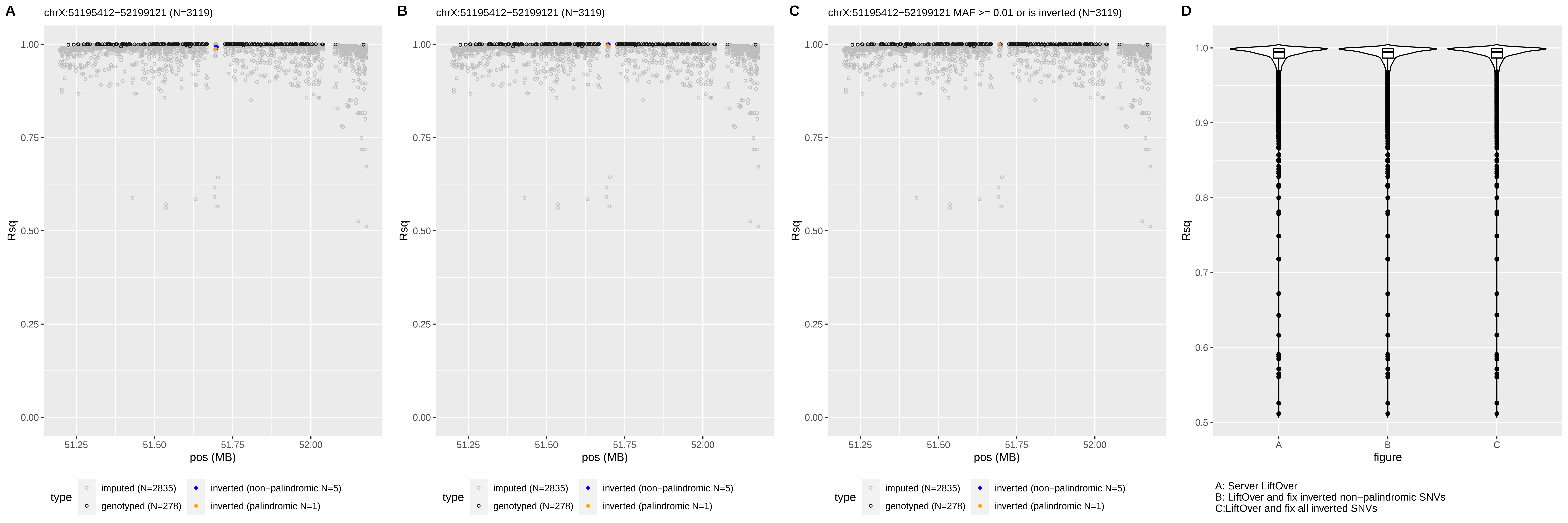
A
chrX:51195412−52199121 (N=3119)
B
chrX:51195412−52199121 (N=3119)
C
chrX:51195412−52199121 MAF >= 0.01 or is inverted (N=3119)
D
Rsq
pos (MB)
1.00
0.75
0.50
0.25
0.00
51.25
51.50
51.75
52.00
type
imputed (N=2835)
genotyped (N=278)
inverted (non−palindromic N=5)
inverted (palindromic N=1)
1.0
0.9
0.8
0.7
0.6
0.5
A
B
C
figure
A: Server LiftOver
B: LiftOver and fix inverted non−palindromic SNVs
C:LiftOver and fix all inverted SNVs

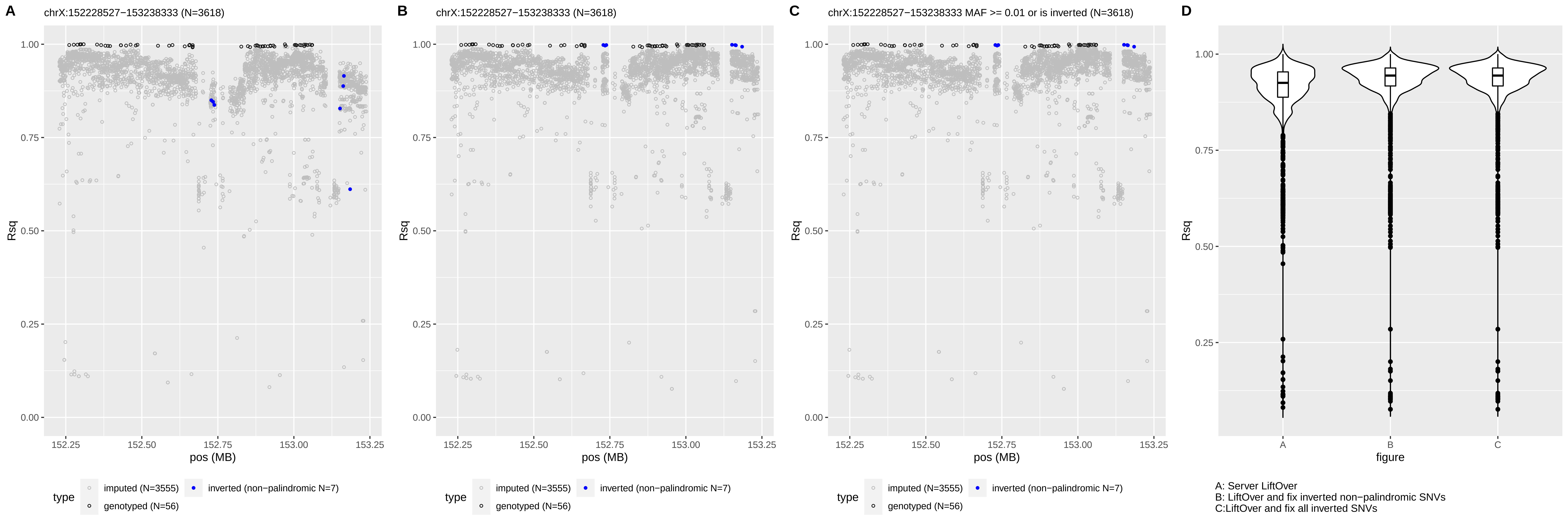

A
chrX:152228527−153238333 (N=3618)
B
chrX:152228527−153238333 (N=3618)
C
chrX:152228527−153238333 MAF >= 0.01 or is inverted (N=3618)
D
Rsq
pos (MB)
figure
type
imputed (N=3555)
inverted (non−palindromic N=7)
genotyped (N=56)
A: Server LiftOver
B: LiftOver and fix inverted non−palindromic SNVs
C:LiftOver and fix all inverted SNVs

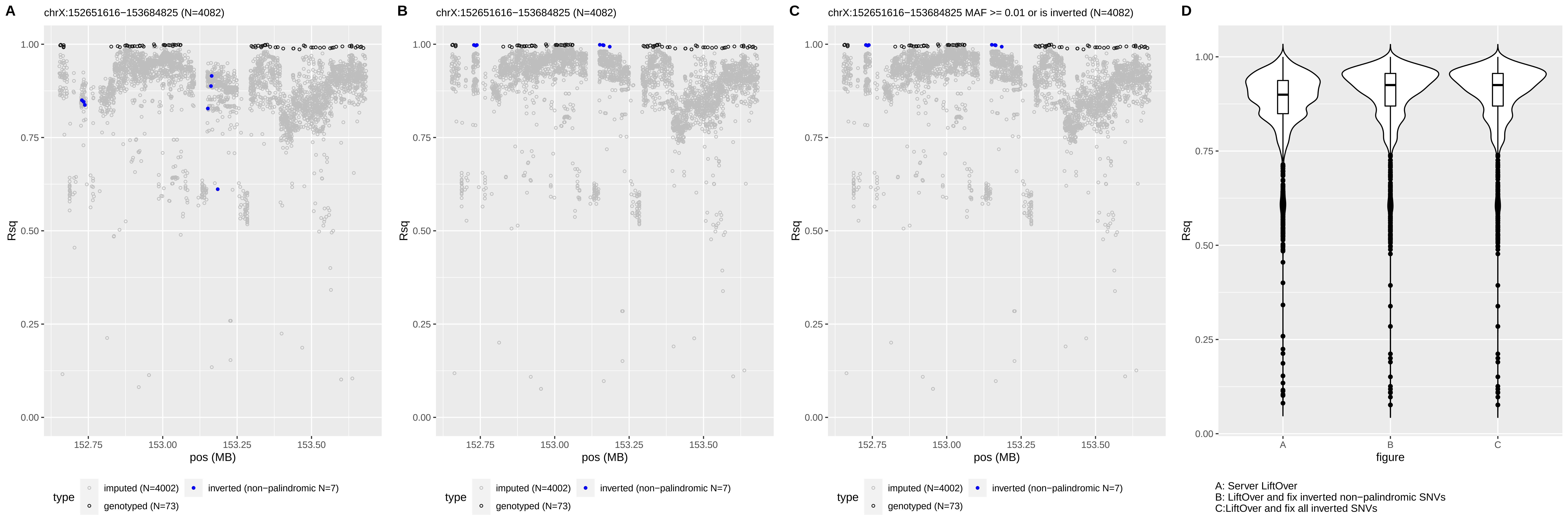
A
chrX:152651616−153684825 (N=4082)
B
chrX:152651616−153684825 (N=4082)
C
chrX:152651616−153684825 MAF >= 0.01 or is inverted (N=4082)
D
Rsq
pos (MB)
figure
type
imputed (N=4002)
inverted (non−palindromic N=7)
genotyped (N=73)
A: Server LiftOver
B: LiftOver and fix inverted non−palindromic SNVs
C:LiftOver and fix all inverted SNVs
