## Supplemental Figure 5 for "Inverted genomic regions between reference genome builds in humans impact imputation accuracy and decrease the power of association testing"

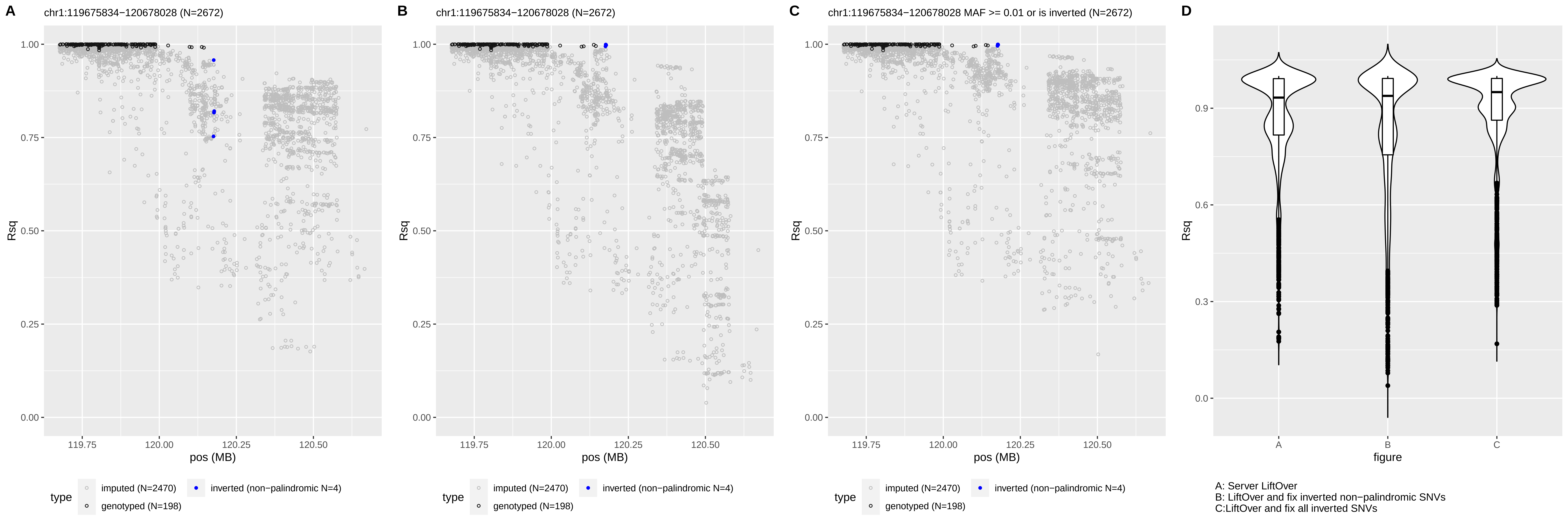

A
chr1:119675834−120678028 (N=2672)
B
chr1:119675834−120678028 (N=2672)
C
chr1:119675834−120678028 MAF >= 0.01 or is inverted (N=2672)
D
Rsq
pos (MB)
figure
type
imputed (N=2470)
genotyped (N=198)
inverted (non−palindromic N=4)
A: Server LiftOver
B: LiftOver and fix inverted non−palindromic SNVs
C:LiftOver and fix all inverted SNVs

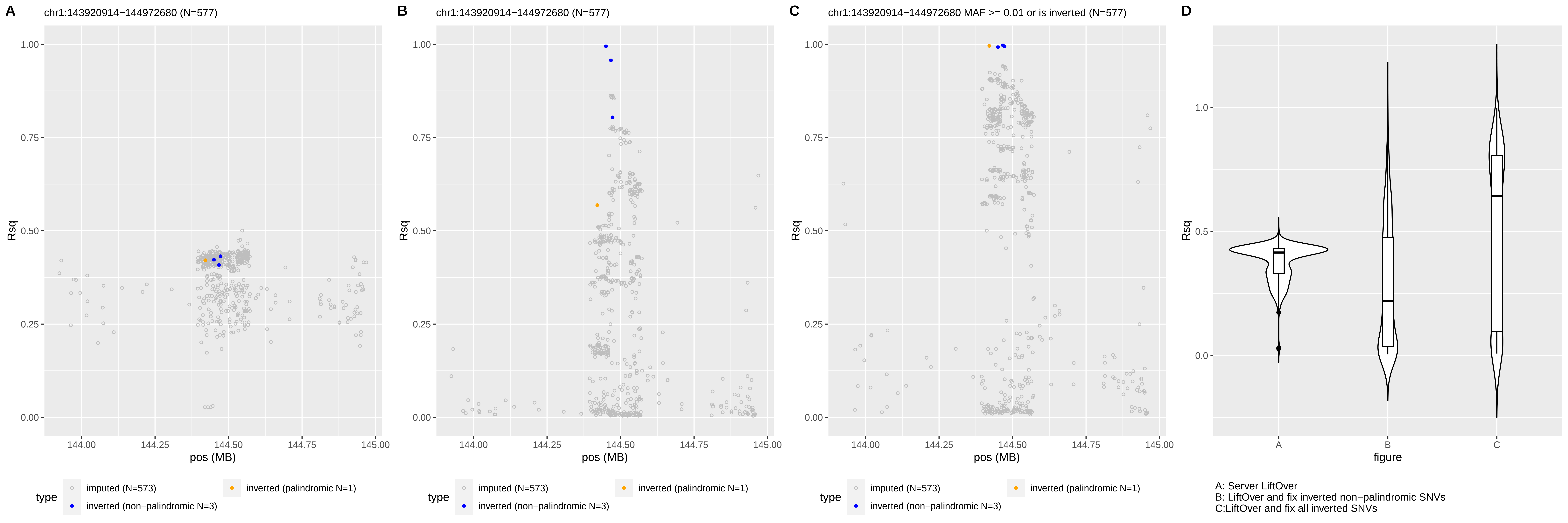
A
chr1:143920914−144972680 (N=577)
B
chr1:143920914−144972680 (N=577)
C
chr1:143920914−144972680 MAF >= 0.01 or is inverted (N=577)
D
Rsq
pos (MB)
figure
type
imputed (N=573)
inverted (palindromic N=1)
inverted (non−palindromic N=3)
A: Server LiftOver
B: LiftOver and fix inverted non−palindromic SNVs
C:LiftOver and fix all inverted SNVs

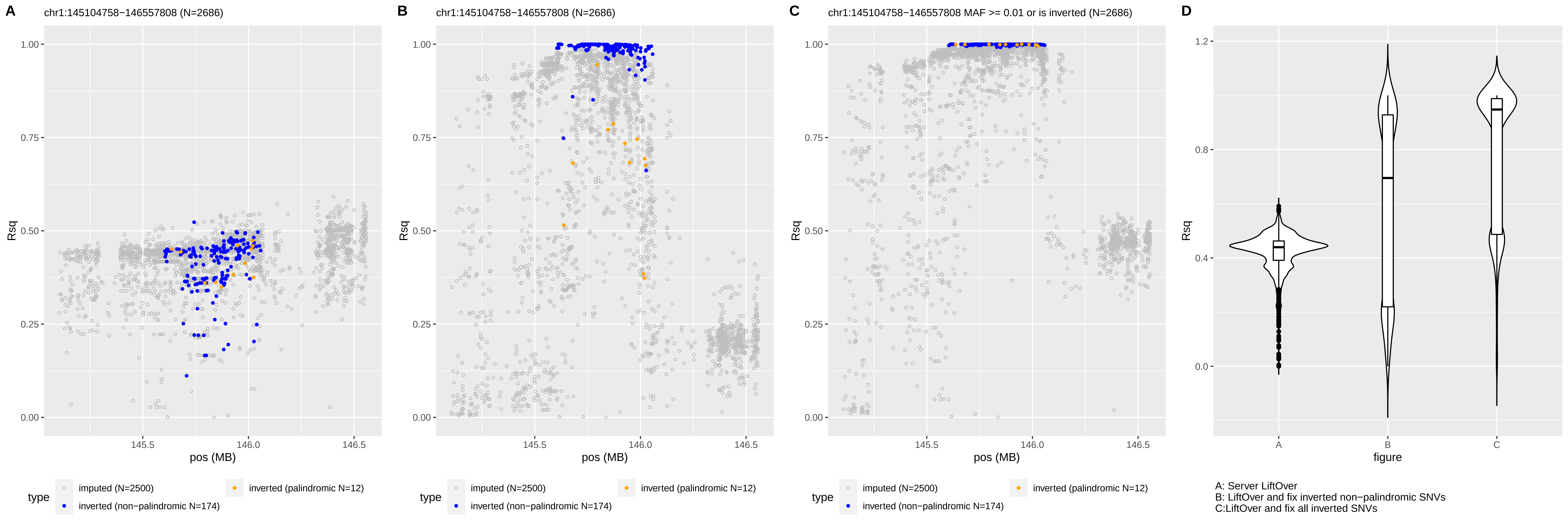
A
chr1:145104758−146557808 (N=2686)
B
chr1:145104758−146557808 (N=2686)
C
chr1:145104758−146557808 MAF >= 0.01 or is inverted (N=2686)
D
Rsq
pos (MB)
figure
type
imputed (N=2500)
inverted (non−palindromic N=174)
inverted (palindromic N=12)
A: Server LiftOver
B: LiftOver and fix inverted non−palindromic SNVs
C:LiftOver and fix all inverted SNVs

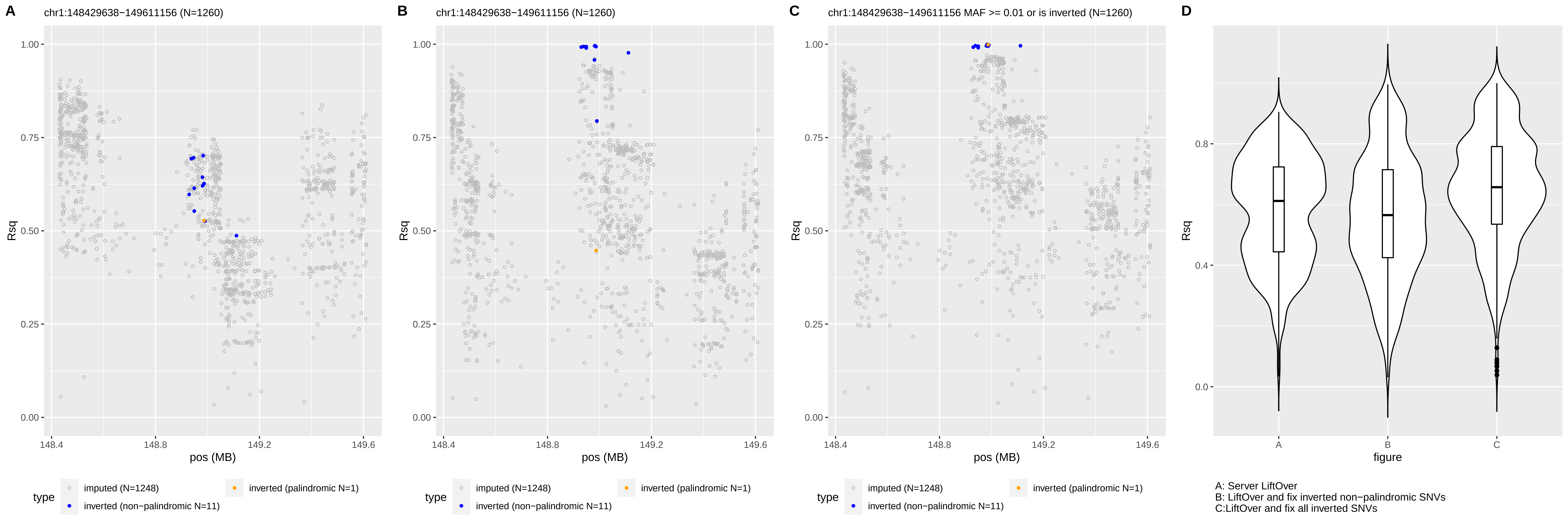
A
chr1:148429638−149611156 (N=1260)
B
chr1:148429638−149611156 (N=1260)
C
chr1:148429638−149611156 MAF >= 0.01 or is inverted (N=1260)
D
Rsq
pos (MB)
figure
type
imputed (N=1248)
inverted (non−palindromic N=11)
inverted (palindromic N=1)
A: Server LiftOver
B: LiftOver and fix inverted non−palindromic SNVs
C:LiftOver and fix all inverted SNVs

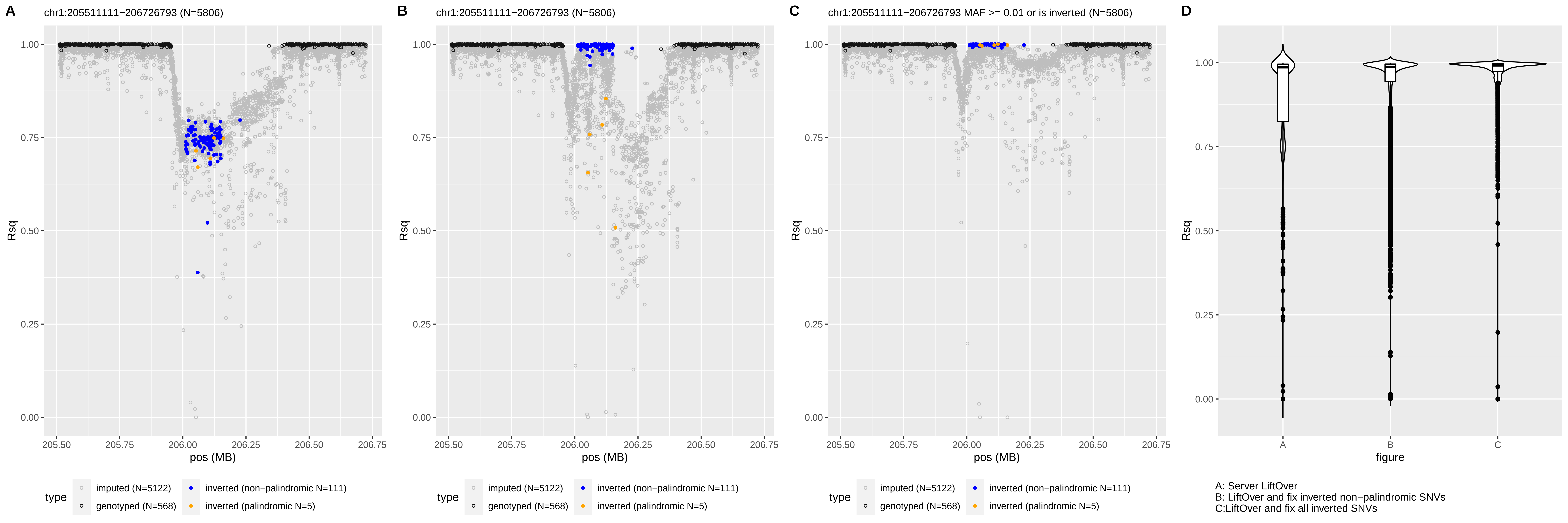
A
chr1:205511111−206726793 (N=5806)
B
chr1:205511111−206726793 (N=5806)
C
chr1:205511111−206726793 MAF >= 0.01 or is inverted (N=5806)
D
Rsq
pos (MB)
figure
type
imputed (N=5122)
genotyped (N=568)
inverted (non−palindromic N=111)
inverted (palindromic N=5)
A: Server LiftOver
B: LiftOver and fix inverted non−palindromic SNVs
C:LiftOver and fix all inverted SNVs

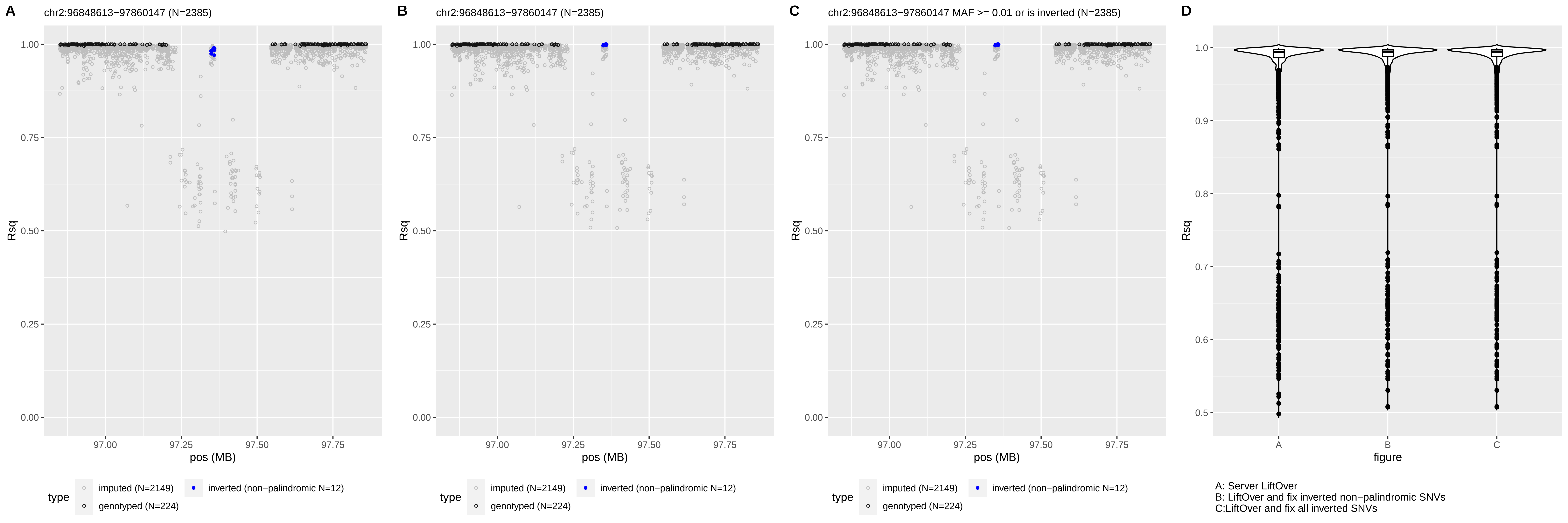

A
chr2:96848613−97860147 (N=2385)
B
chr2:96848613−97860147 (N=2385)
C
chr2:96848613−97860147 MAF >= 0.01 or is inverted (N=2385)
D
Rsq
pos (MB)
figure
type
imputed (N=2149)
inverted (non−palindromic N=12)
genotyped (N=224)
A: Server LiftOver
B: LiftOver and fix inverted non−palindromic SNVs
C:LiftOver and fix all inverted SNVs

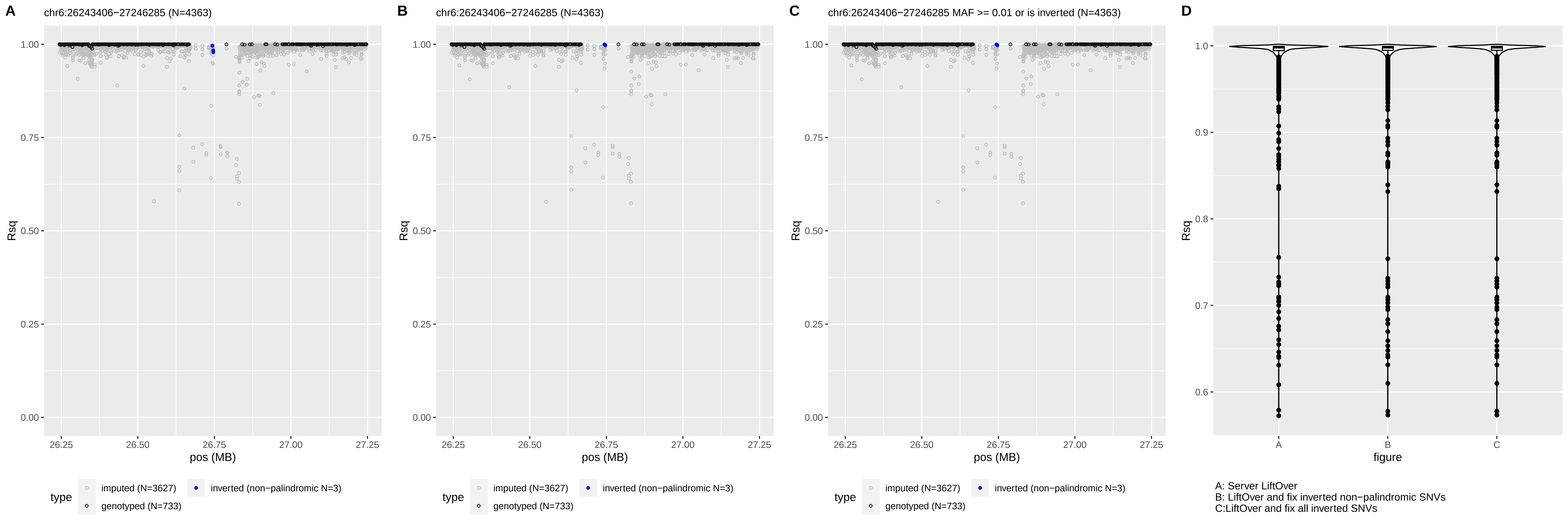
A
chr6:26243406−27246285 (N=4363)
B
chr6:26243406−27246285 (N=4363)
C
chr6:26243406−27246285 MAF >= 0.01 or is inverted (N=4363)
D
Rsq
pos (MB)
type
imputed (N=3627)
genotyped (N=733)
inverted (non−palindromic N=3)
figure
A: Server LiftOver
B: LiftOver and fix inverted non−palindromic SNVs
C:LiftOver and fix all inverted SNVs

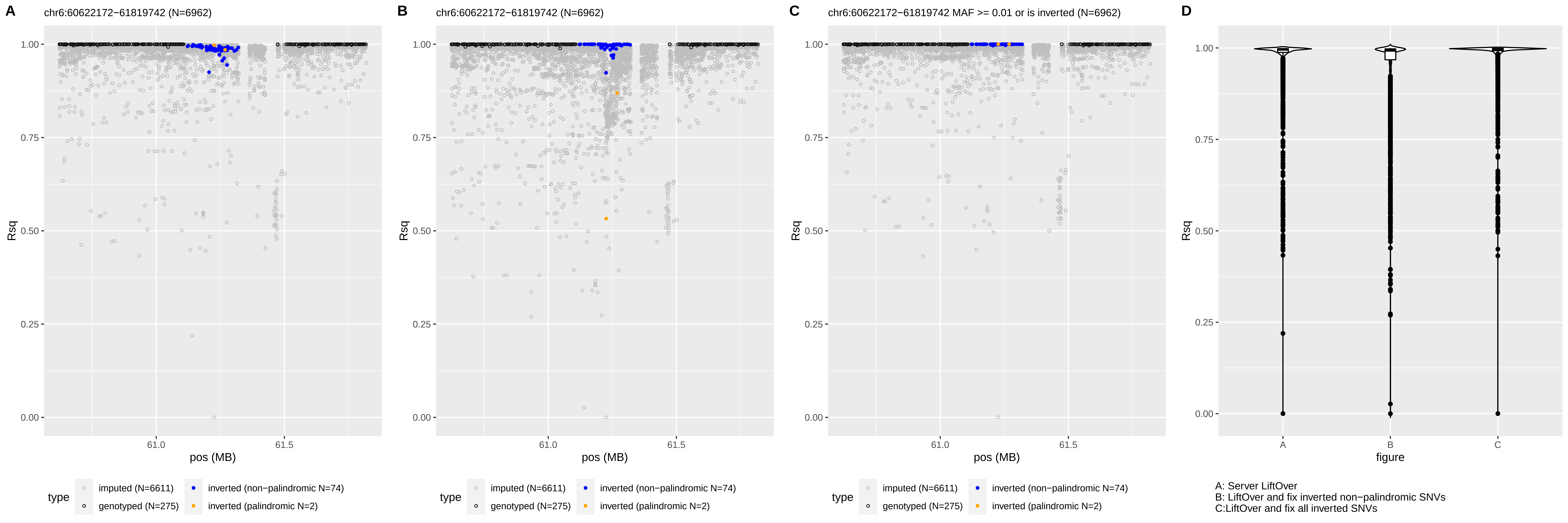
A
chr6:60622172−61819742 (N=6962)
B
chr6:60622172−61819742 (N=6962)
C
chr6:60622172−61819742 MAF >= 0.01 or is inverted (N=6962)
D
Rsq
pos (MB)
figure
type
imputed (N=6611)
genotyped (N=275)
inverted (non−palindromic N=74)
inverted (palindromic N=2)
A: Server LiftOver
B: LiftOver and fix inverted non−palindromic SNVs
C:LiftOver and fix all inverted SNVs

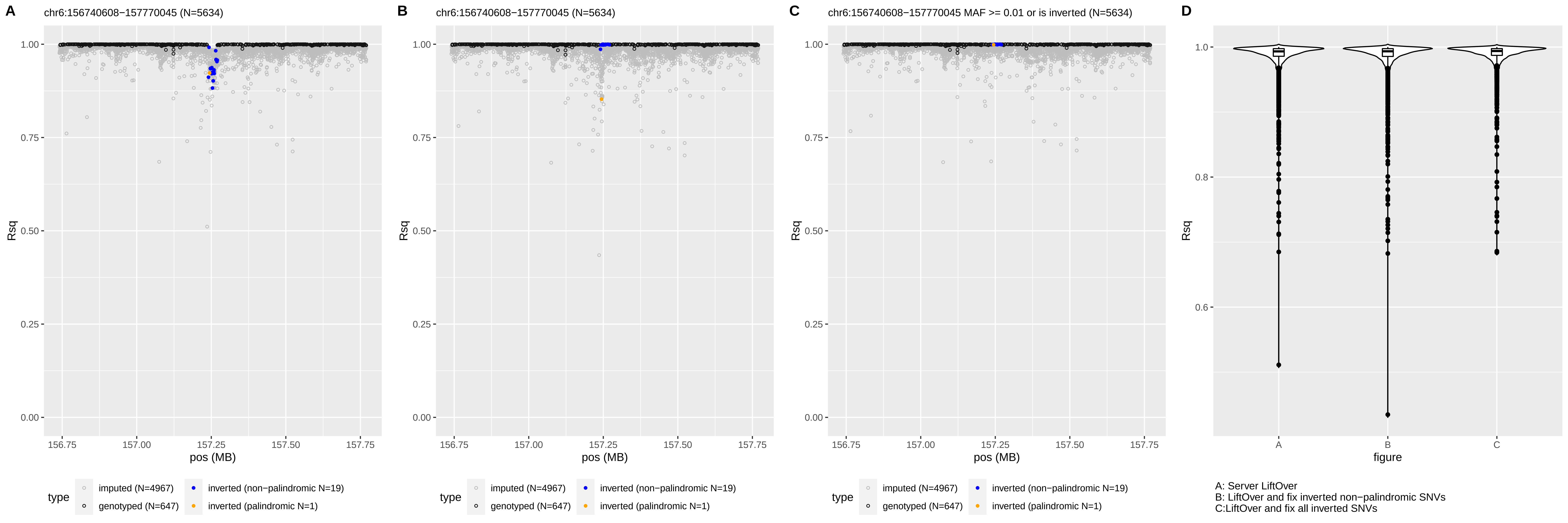

A
chr6:156740608−157770045 (N=5634)
B
chr6:156740608−157770045 (N=5634)
C
chr6:156740608−157770045 MAF >= 0.01 or is inverted (N=5634)
D
Rsq
pos (MB)
figure
type
imputed (N=4967)
genotyped (N=647)
inverted (non−palindromic N=19)
inverted (palindromic N=1)
A: Server LiftOver
B: LiftOver and fix inverted non−palindromic SNVs
C:LiftOver and fix all inverted SNVs

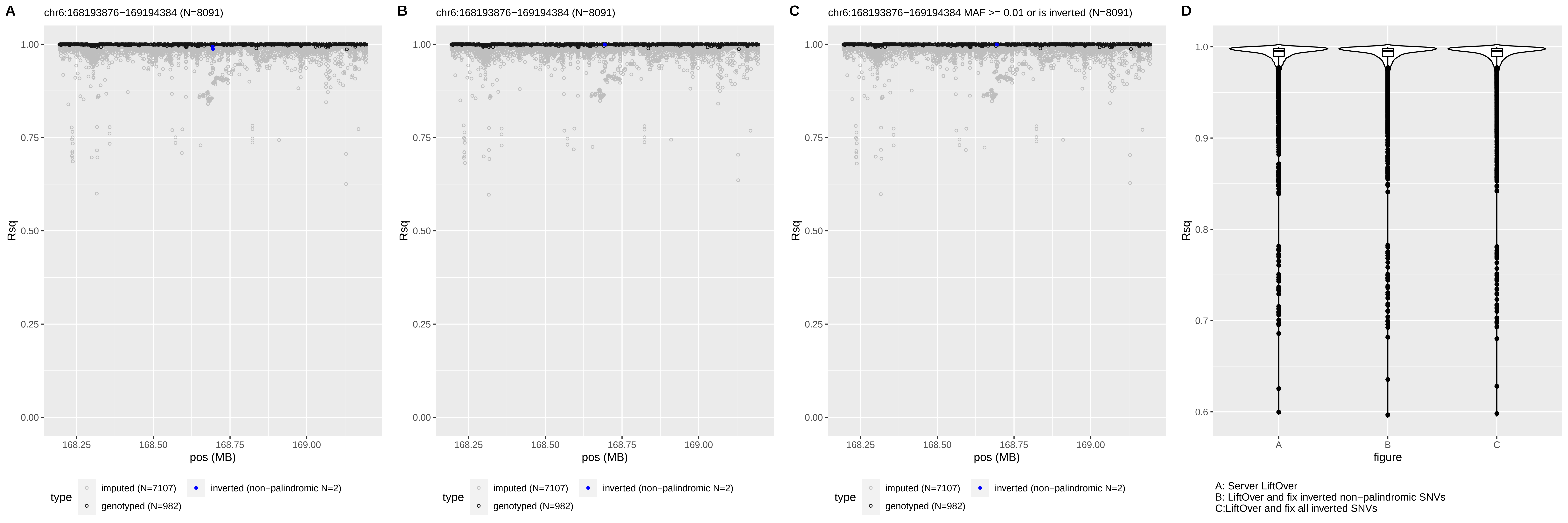

A
chr6:168193876−169194384 (N=8091)
B
chr6:168193876−169194384 (N=8091)
C
chr6:168193876−169194384 MAF >= 0.01 or is inverted (N=8091)
D
Rsq
pos (MB)
figure
type
imputed (N=7107)
genotyped (N=982)
inverted (non−palindromic N=2)
A: Server LiftOver
B: LiftOver and fix inverted non−palindromic SNVs
C:LiftOver and fix all inverted SNVs

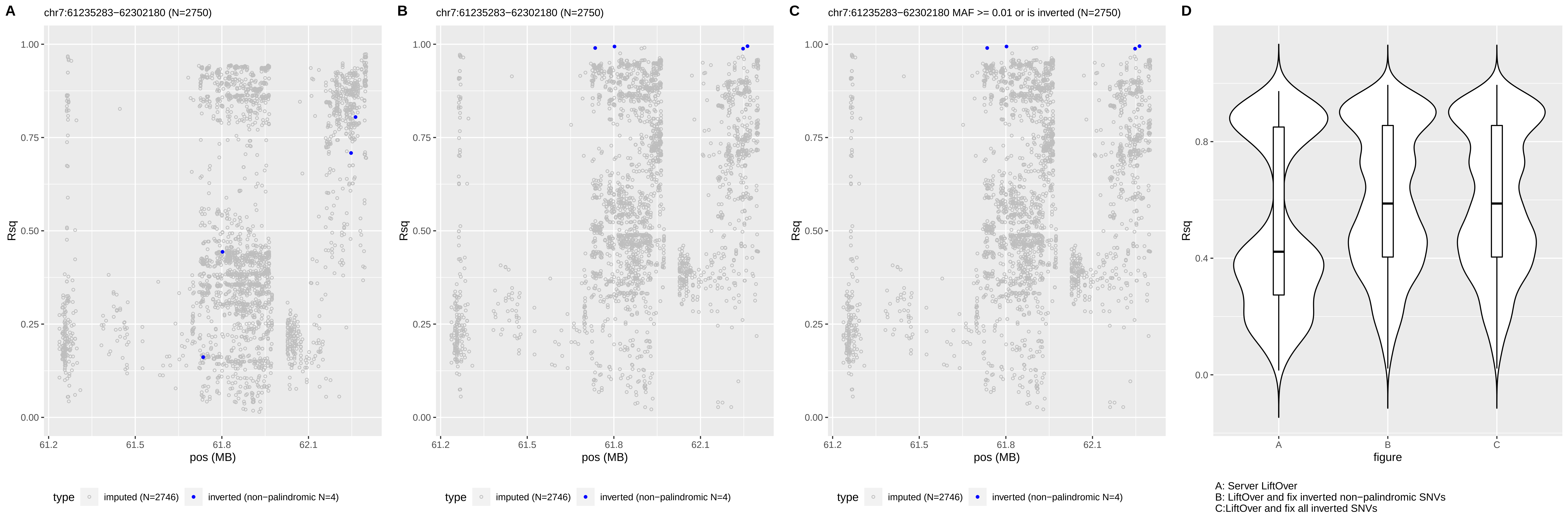
A
chr7:61235283−62302180 (N=2750)
B
chr7:61235283−62302180 (N=2750)
C
chr7:61235283−62302180 MAF >= 0.01 or is inverted (N=2750)
D
Rsq
pos (MB)
figure
type
imputed (N=2746)
inverted (non−palindromic N=4)
A: Server LiftOver
B: LiftOver and fix inverted non−palindromic SNVs
C:LiftOver and fix all inverted SNVs

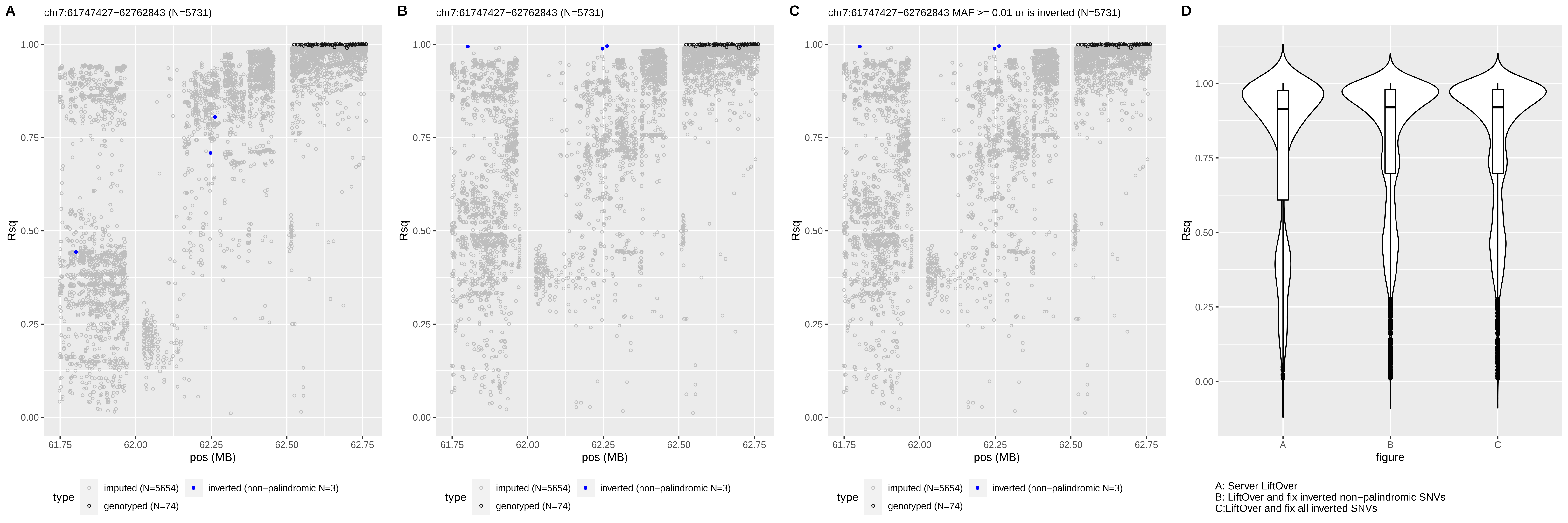
A
chr7:61747427−62763843 (N=5731)
B
chr7:61747427−62763843 (N=5731)
C
chr7:61747427−62763843 MAF >= 0.01 or is inverted (N=5731)
D
Rsq
pos (MB)
figure
type
imputed (N=5654)
genotyped (N=74)
inverted (non−palindromic N=3)
A: Server LiftOver
B: LiftOver and fix inverted non−palindromic SNVs
C:LiftOver and fix all inverted SNVs

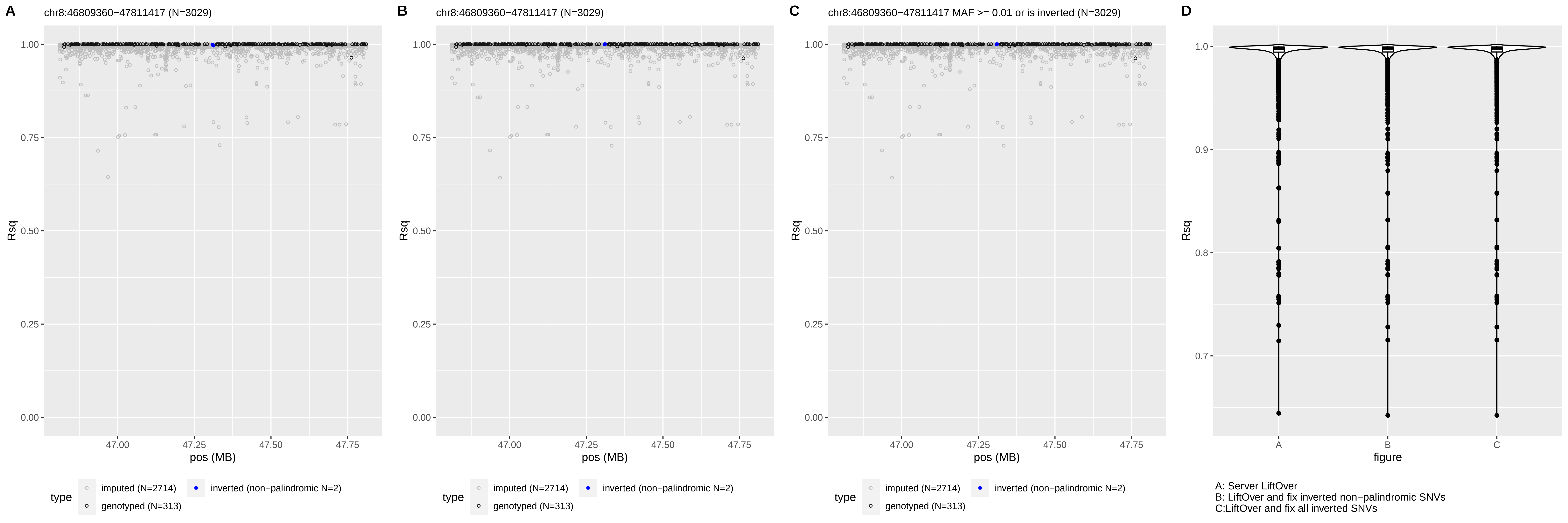
A
chr8:46809360−47811417 (N=3029)
B
chr8:46809360−47811417 (N=3029)
C
chr8:46809360−47811417 MAF >= 0.01 or is inverted (N=3029)
D
Rsq
pos (MB)
figure
type
imputed (N=2714)
genotyped (N=313)
inverted (non−palindromic N=2)
A: Server LiftOver
B: LiftOver and fix inverted non−palindromic SNVs
C:LiftOver and fix all inverted SNVs

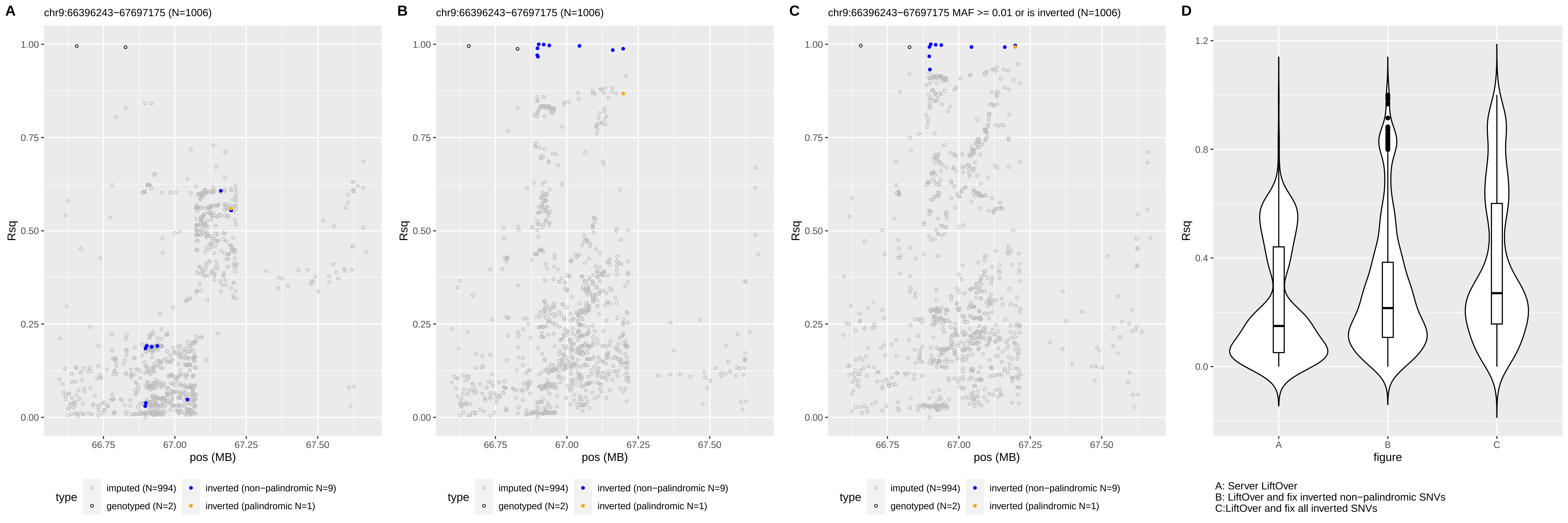

A
chr9:66396243−67697175 (N=1006)
B
chr9:66396243−67697175 (N=1006)
C
chr9:66396243−67697175 MAF >= 0.01 or is inverted (N=1006)
D
Rsq
pos (MB)
figure
type
imputed (N=994)
genotyped (N=2)
inverted (non−palindromic N=9)
inverted (palindromic N=1)
A: Server LiftOver
B: LiftOver and fix inverted non−palindromic SNVs
C:LiftOver and fix all inverted SNVs

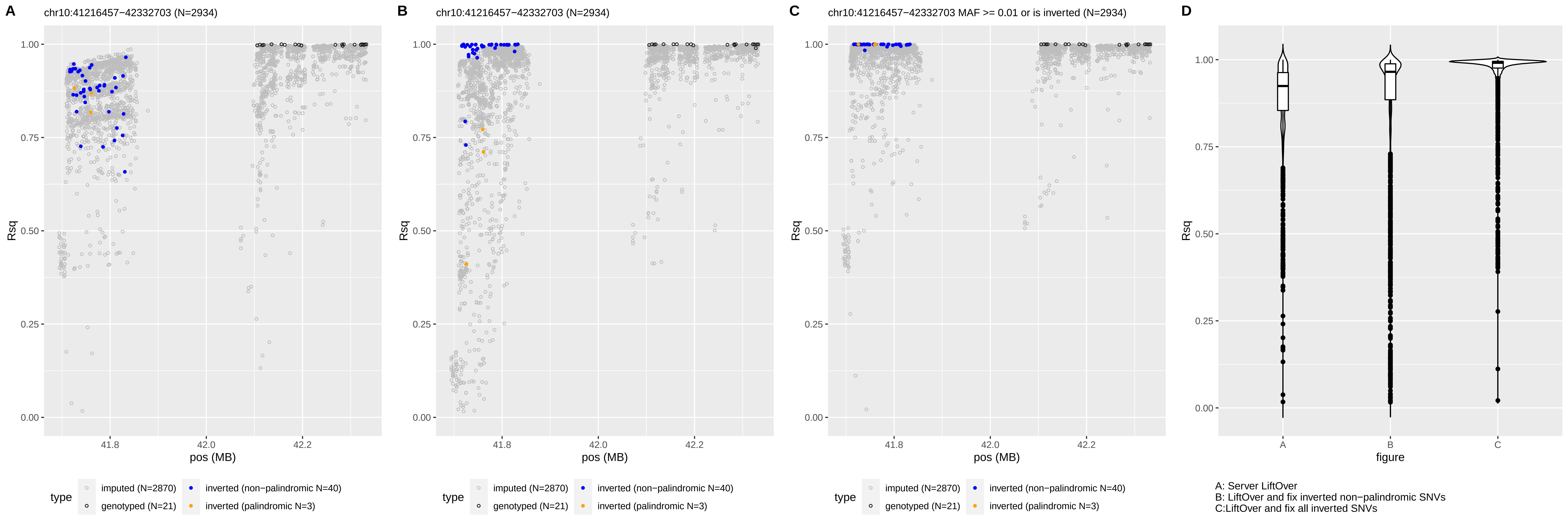
A
chr10:41216457−42332703 (N=2934)
B
chr10:41216457−42332703 (N=2934)
C
chr10:41216457−42332703 MAF >= 0.01 or is inverted (N=2934)
D
Rsq
pos (MB)
41.8
42.0
42.2
0.00
0.25
0.50
0.75
1.00
type
imputed (N=2870)
genotyped (N=21)
inverted (non−palindromic N=40)
inverted (palindromic N=3)
figure
A
B
C
A: Server LiftOver
B: LiftOver and fix inverted non−palindromic SNVs
C:LiftOver and fix all inverted SNVs

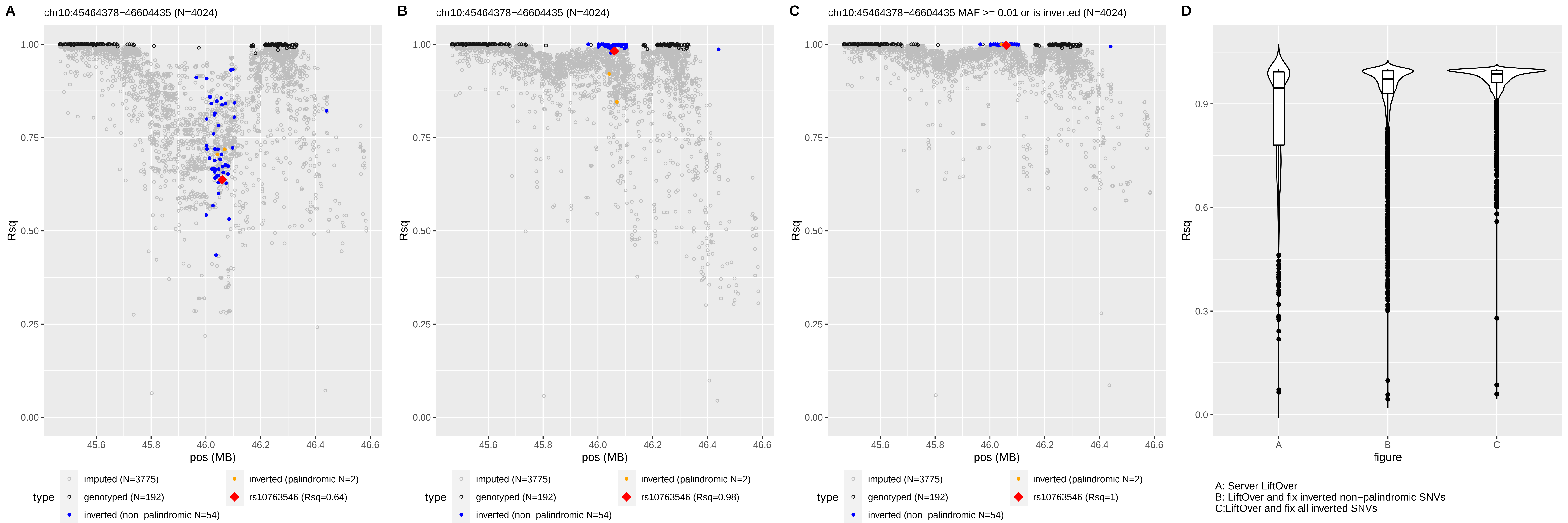
A
chr10:45464378−46604435 (N=4024)
B
chr10:45464378−46604435 (N=4024)
C
chr10:45464378−46604435 MAF >= 0.01 or is inverted (N=4024)
D
Rsq
pos (MB)
figure
type
imputed (N=3775)
genotyped (N=192)
inverted (non−palindromic N=54)
inverted (palindromic N=2)
rs10763546 (Rsq=0.64)
rs10763546 (Rsq=0.98)
rs10763546 (Rsq=1)
A: Server LiftOver
B: LiftOver and fix inverted non−palindromic SNVs
C:LiftOver and fix all inverted SNVs

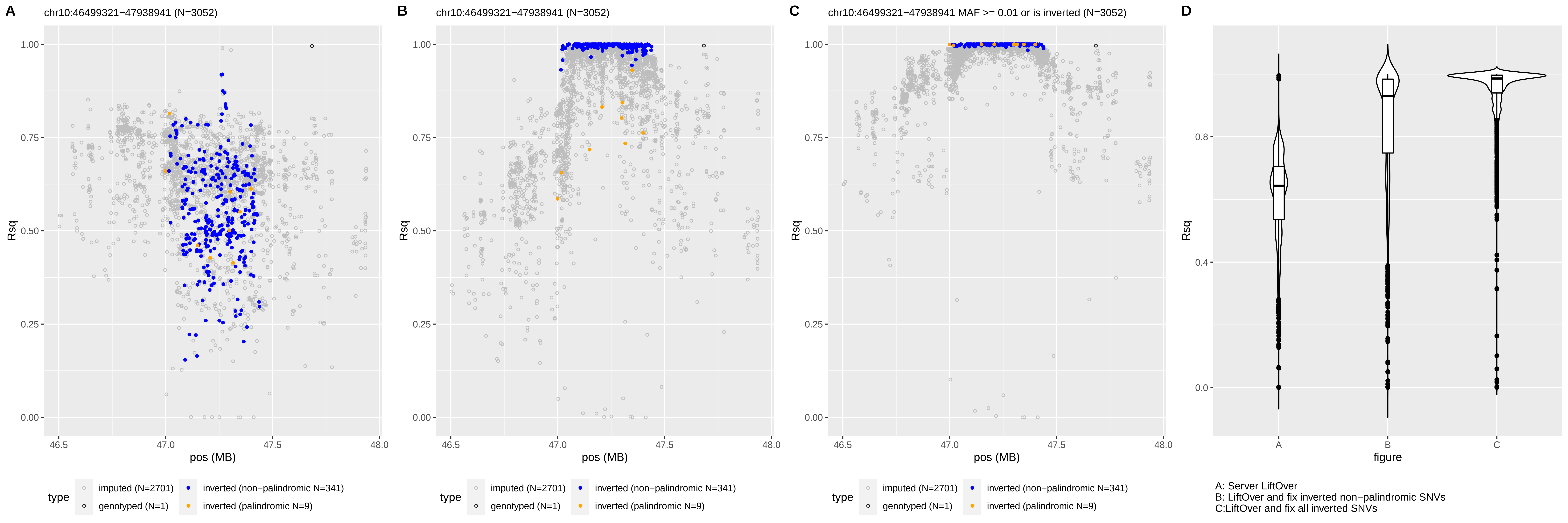
A
chr10:46499321−47938941 (N=3052)
B
chr10:46499321−47938941 (N=3052)
C
chr10:46499321−47938941 MAF >= 0.01 or is inverted (N=3052)
D
Rsq
pos (MB)
figure
type
imputed (N=2701)
genotyped (N=1)
inverted (non−palindromic N=341)
inverted (palindromic N=9)
A: Server LiftOver
B: LiftOver and fix inverted non−palindromic SNVs
C:LiftOver and fix all inverted SNVs

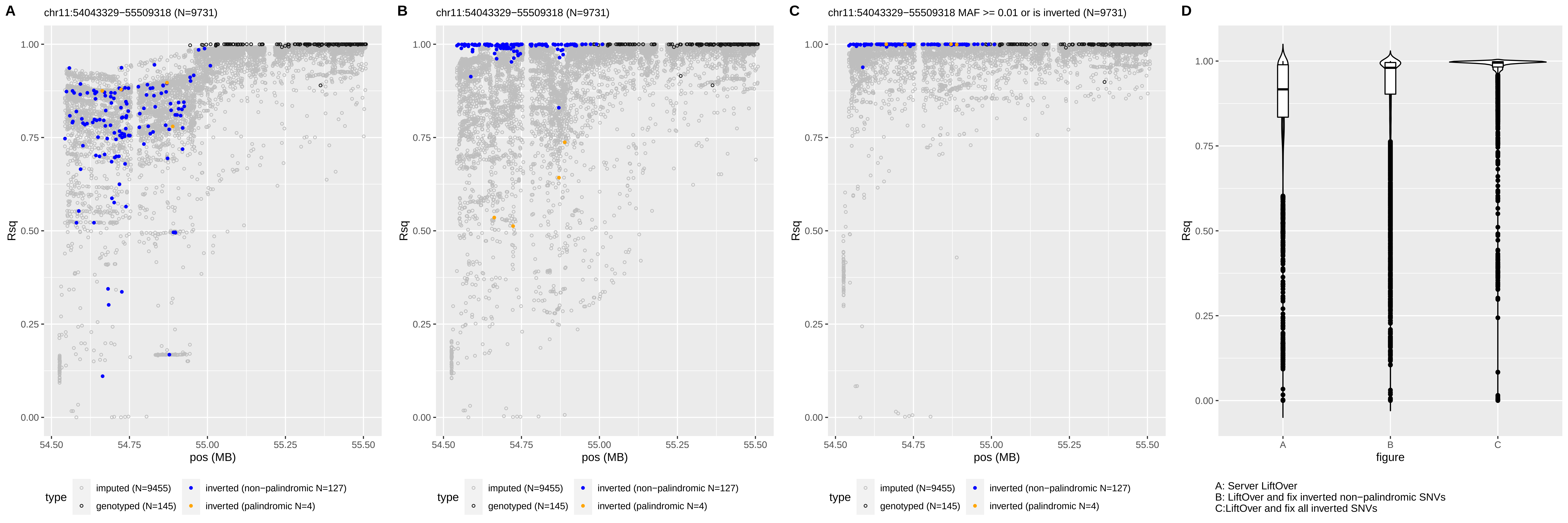
A
chr11:54043329−55509318 (N=9731)
B
chr11:54043329−55509318 (N=9731)
C
chr11:54043329−55509318 MAF >= 0.01 or is inverted (N=9731)
D
Rsq
pos (MB)
figure
type
imputed (N=9455)
genotyped (N=145)
inverted (non−palindromic N=127)
inverted (palindromic N=4)
A: Server LiftOver
B: LiftOver and fix inverted non−palindromic SNVs
C:LiftOver and fix all inverted SNVs

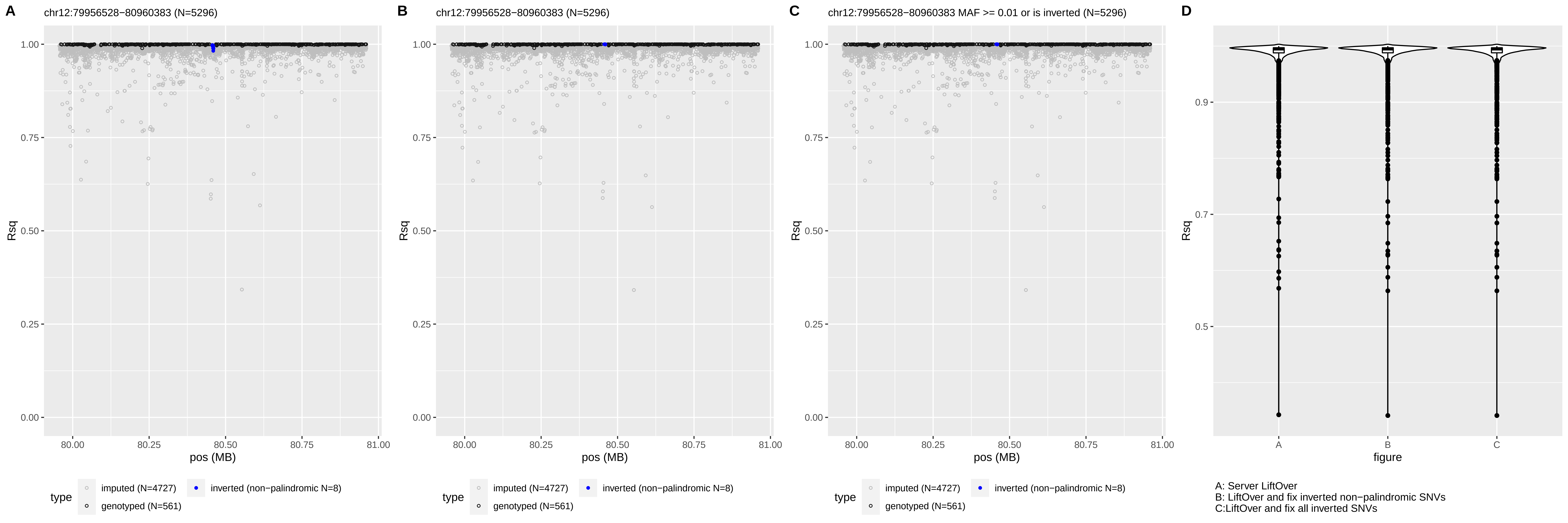

A
chr12:79956528−80960383 (N=5296)
B
chr12:79956528−80960383 (N=5296)
C
chr12:79956528−80960383 MAF >= 0.01 or is inverted (N=5296)
D
Rsq
pos (MB)
type
imputed (N=4727)
genotyped (N=561)
inverted (non−palindromic N=8)
figure
A: Server LiftOver
B: LiftOver and fix inverted non−palindromic SNVs
C:LiftOver and fix all inverted SNVs

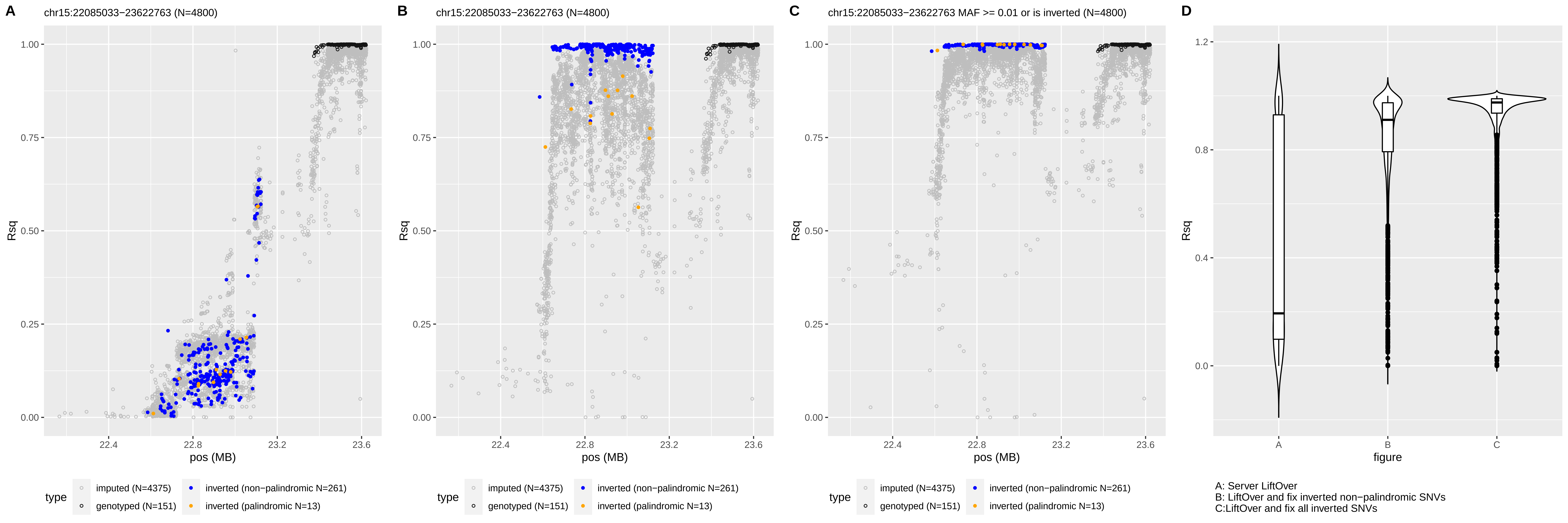
A
chr15:22085033−23622763 (N=4800)
B
chr15:22085033−23622763 (N=4800)
C
chr15:22085033−23622763 MAF >= 0.01 or is inverted (N=4800)
D
Rsq
pos (MB)
figure
type
imputed (N=4375)
genotyped (N=151)
inverted (non−palindromic N=261)
inverted (palindromic N=13)
A: Server LiftOver
B: LiftOver and fix inverted non−palindromic SNVs
C:LiftOver and fix all inverted SNVs

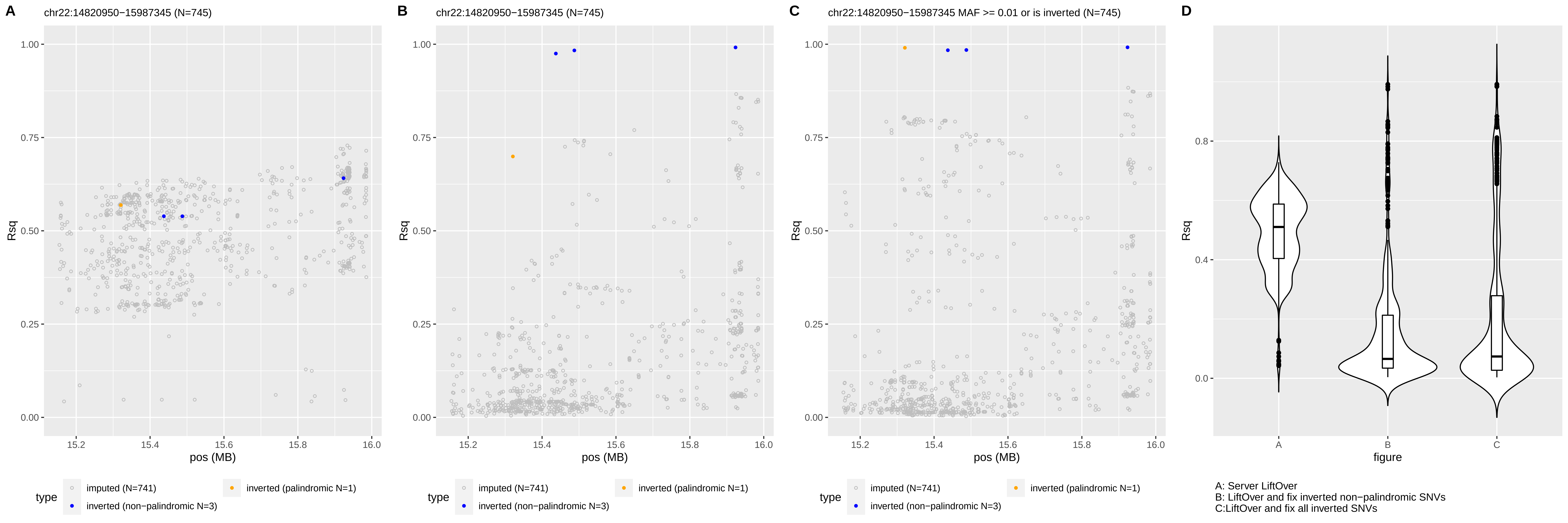
A
chr22:14820950−15987345 (N=745)
B
chr22:14820950−15987345 (N=745)
C
chr22:14820950−15987345 MAF >= 0.01 or is inverted (N=745)
D
Rsq
pos (MB)
figure
type
imputed (N=741)
inverted (palindromic N=1)
inverted (non−palindromic N=3)
A: Server LiftOver
B: LiftOver and fix inverted non−palindromic SNVs
C:LiftOver and fix all inverted SNVs

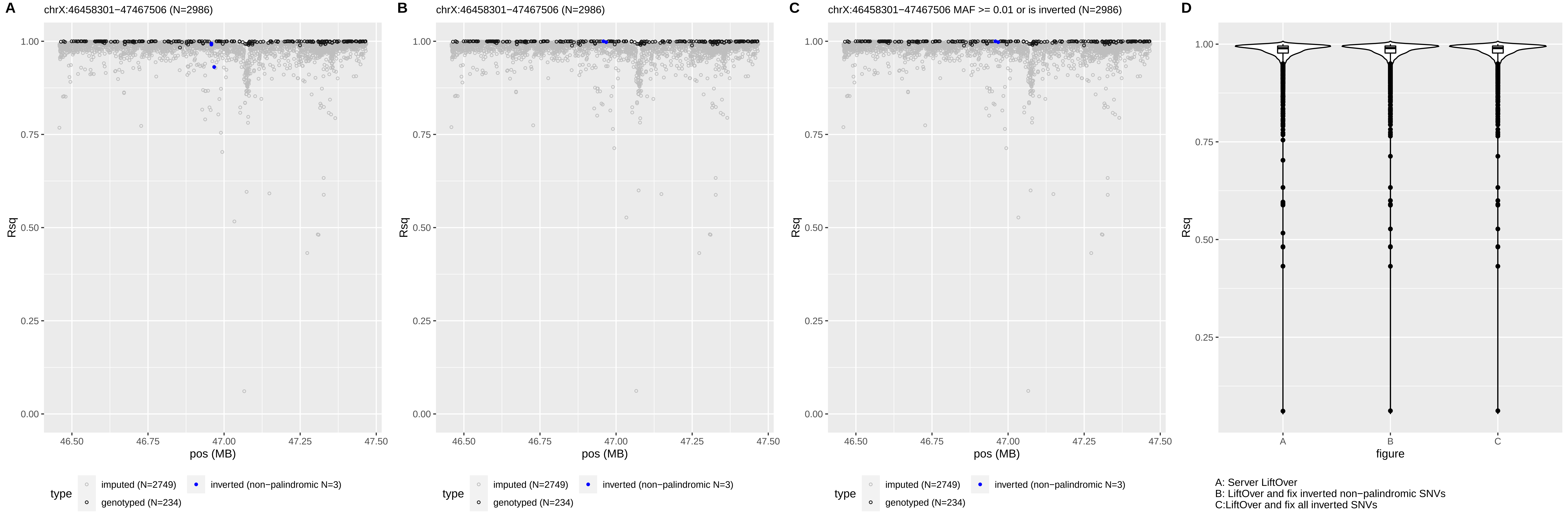
A
chrX:46458301−47467506 (N=2986)
B
chrX:46458301−47467506 (N=2986)
C
chrX:46458301−47467506 MAF >= 0.01 or is inverted (N=2986)
D
Rsq
pos (MB)
figure
type
imputed (N=2749)
genotyped (N=234)
inverted (non−palindromic N=3)
A: Server LiftOver
B: LiftOver and fix inverted non−palindromic SNVs
C:LiftOver and fix all inverted SNVs

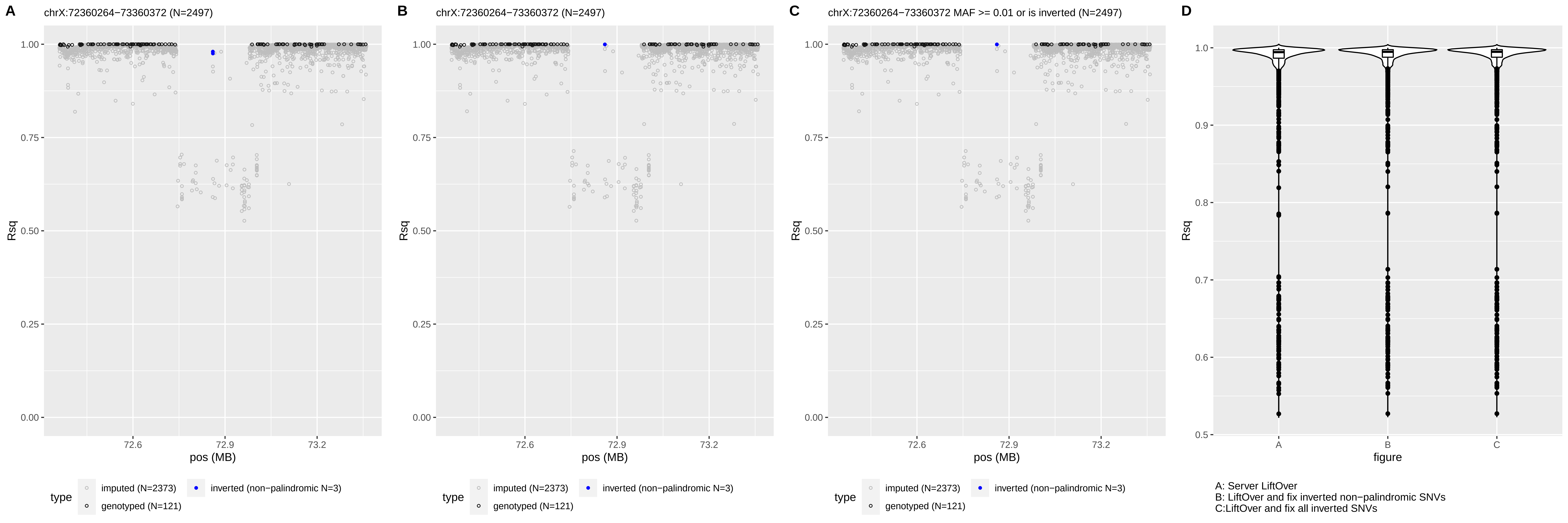
A
chrX:72360264−73360372 (N=2497)
B
chrX:72360264−73360372 (N=2497)
C
chrX:72360264−73360372 MAF >= 0.01 or is inverted (N=2497)
D
Rsq
pos (MB)
figure
type
imputed (N=2373)
genotyped (N=121)
inverted (non−palindromic N=3)
A: Server LiftOver
B: LiftOver and fix inverted non−palindromic SNVs
C:LiftOver and fix all inverted SNVs

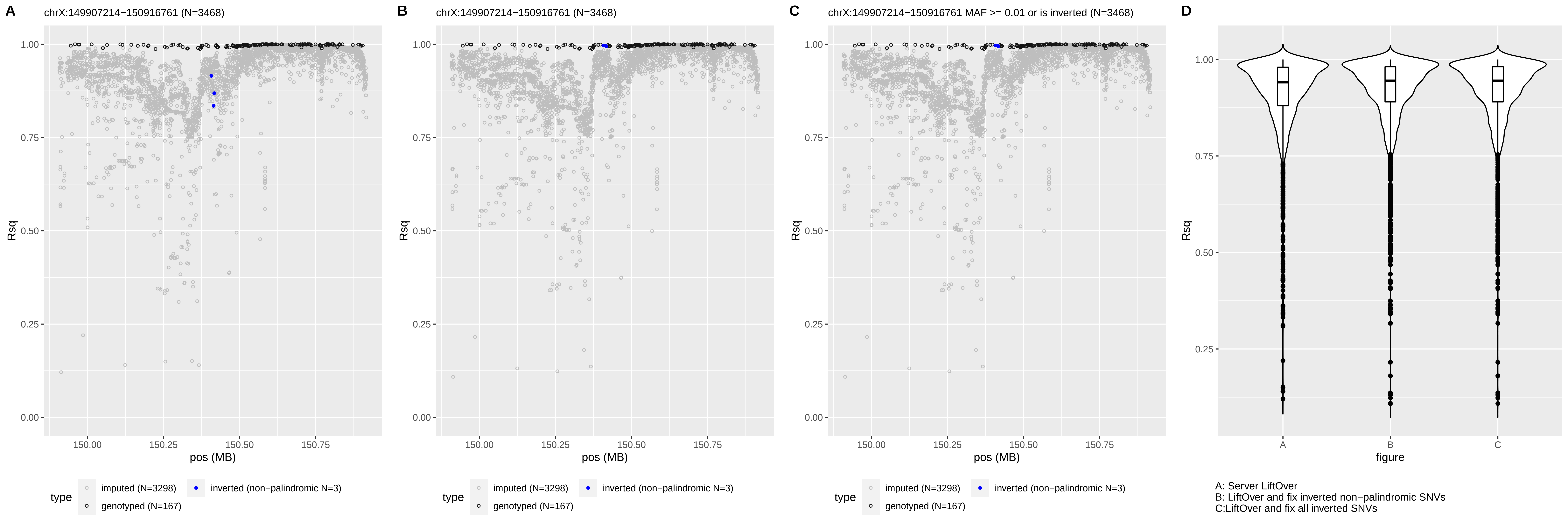

A
chrX:149907214−150916761 (N=3468)
B
chrX:149907214−150916761 (N=3468)
C
chrX:149907214−150916761 MAF >= 0.01 or is inverted (N=3468)
D
Rsq
pos (MB)
figure
type
imputed (N=3298)
inverted (non−palindromic N=3)
genotyped (N=167)
A: Server LiftOver
B: LiftOver and fix inverted non−palindromic SNVs
C:LiftOver and fix all inverted SNVs

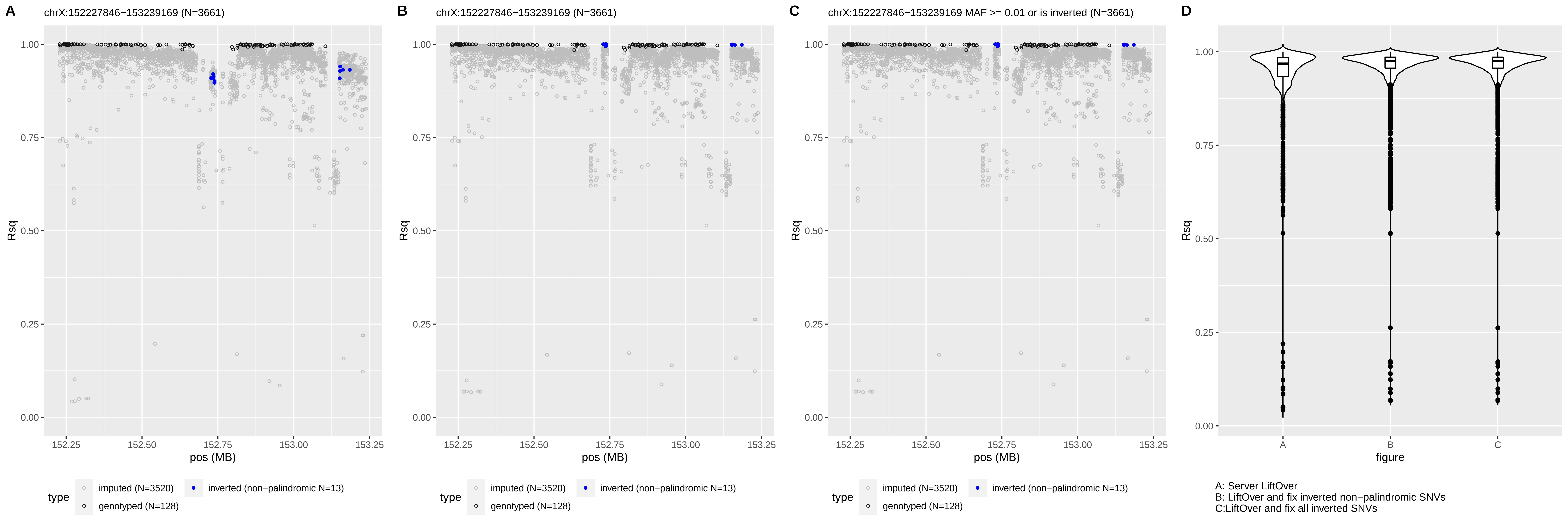
A
chrX:152227846−153239169 (N=3661)
B
chrX:152227846−153239169 (N=3661)
C
chrX:152227846−153239169 MAF >= 0.01 or is inverted (N=3661)
D
Rsq
1.00
0.75
0.50
0.25
0.00
pos (MB)
152.25
152.50
152.75
153.00
153.25
type
imputed (N=3520)
genotyped (N=128)
inverted (non−palindromic N=13)
figure
A
B
C
A: Server LiftOver
B: LiftOver and fix inverted non−palindromic SNVs
C:LiftOver and fix all inverted SNVs

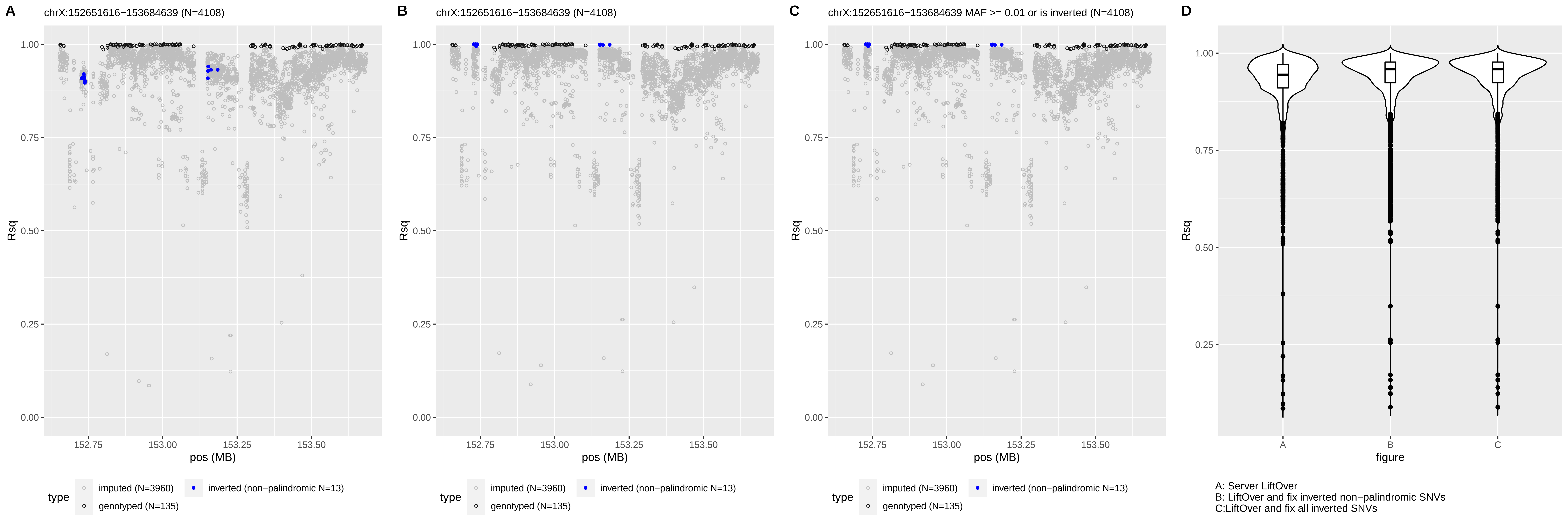
A
chrX:152651616−153684639 (N=4108)
B
chrX:152651616−153684639 (N=4108)
C
chrX:152651616−153684639 MAF >= 0.01 or is inverted (N=4108)
D
Rsq
pos (MB)
type
imputed (N=3960)
inverted (non−palindromic N=13)
genotyped (N=135)
figure
A: Server LiftOver
B: LiftOver and fix inverted non−palindromic SNVs
C:LiftOver and fix all inverted SNVs
