## Supplemental Figure 6 for "Inverted genomic regions between reference genome builds in humans impact imputation accuracy and decrease the power of association testing"

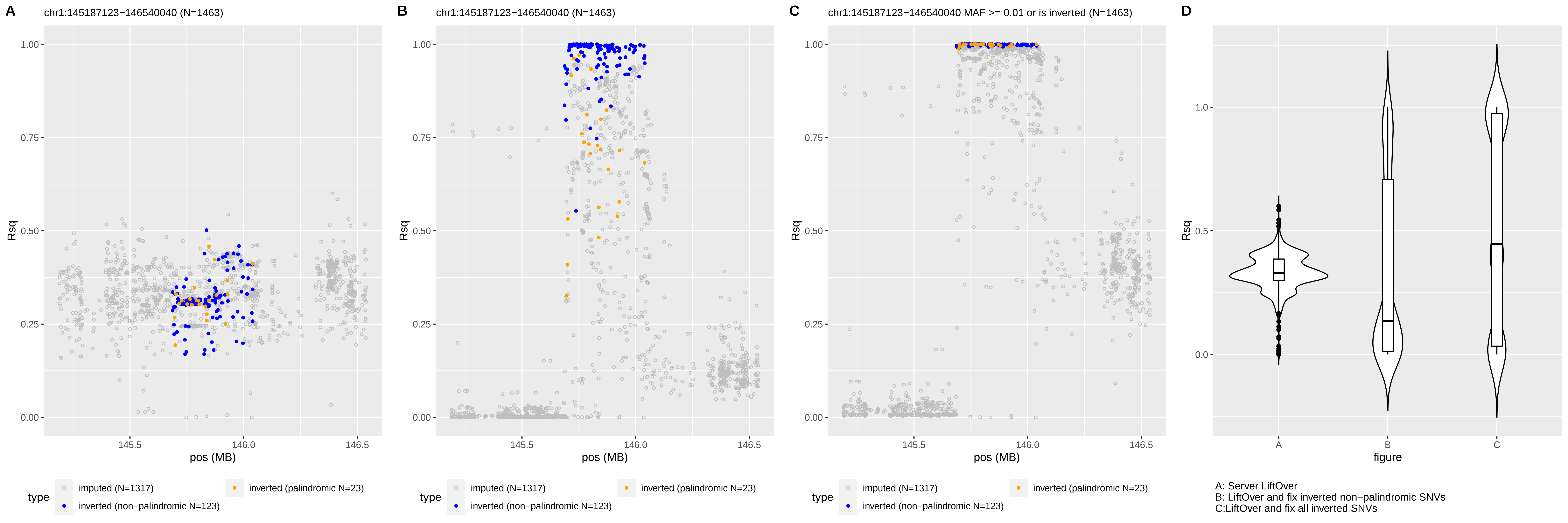
A
chr1:145187123−146540040 (N=1463)
B
chr1:145187123−146540040 (N=1463)
C
chr1:145187123−146540040 MAF >= 0.01 or is inverted (N=1463)
D
Rsq
pos (MB)
type
imputed (N=1317)
inverted (non−palindromic N=123)
inverted (palindromic N=23)
figure
A: Server LiftOver
B: LiftOver and fix inverted non−palindromic SNVs
C:LiftOver and fix all inverted SNVs

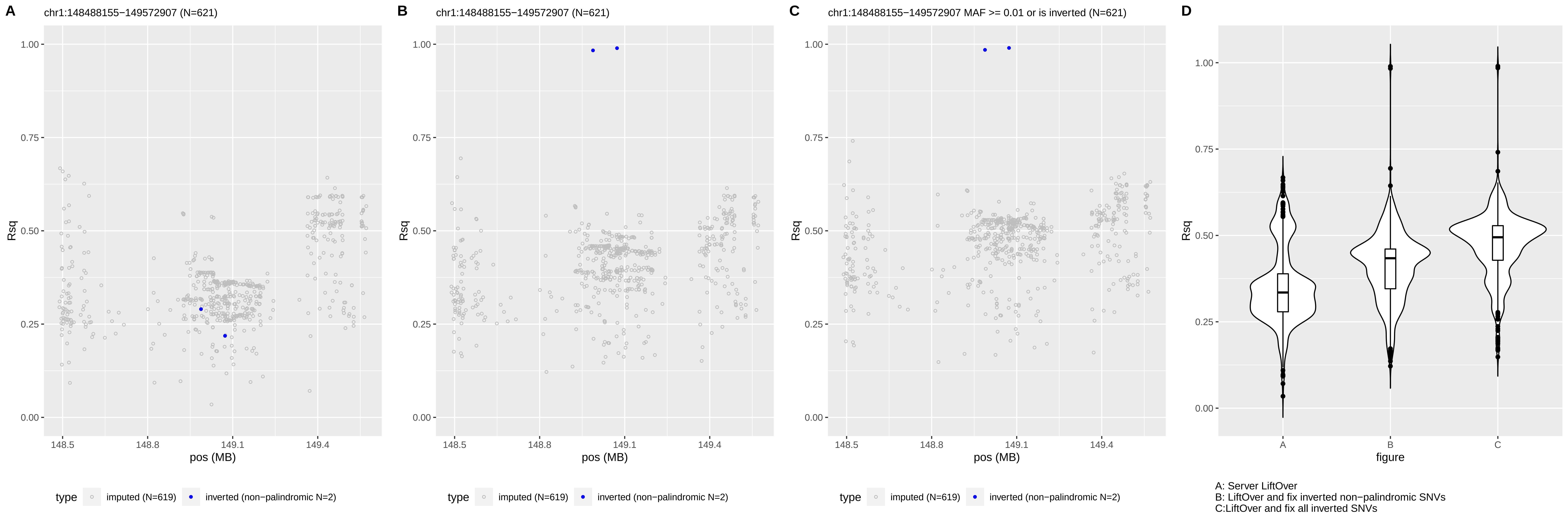

A
chr1:148488155−149572907 (N=621)
B
chr1:148488155−149572907 (N=621)
C
chr1:148488155−149572907 MAF >= 0.01 or is inverted (N=621)
D
Rsq
pos (MB)
figure
type
imputed (N=619)
inverted (non−palindromic N=2)
A: Server LiftOver
B: LiftOver and fix inverted non−palindromic SNVs
C:LiftOver and fix all inverted SNVs

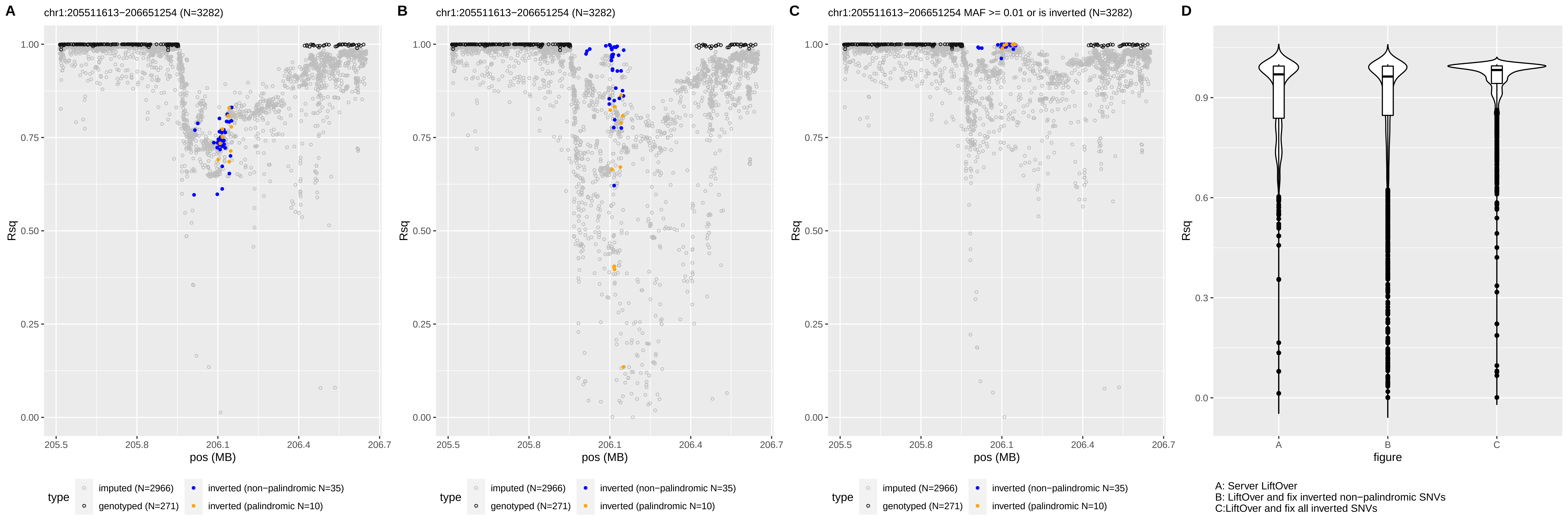

A
chr1:205511613−206651254 (N=3282)
B
chr1:205511613−206651254 (N=3282)
C
chr1:205511613−206651254 MAF >= 0.01 or is inverted (N=3282)
D
Rsq
pos (MB)
figure
type
imputed (N=2966)
genotyped (N=271)
inverted (non−palindromic N=35)
inverted (palindromic N=10)
A: Server LiftOver
B: LiftOver and fix inverted non−palindromic SNVs
C:LiftOver and fix all inverted SNVs

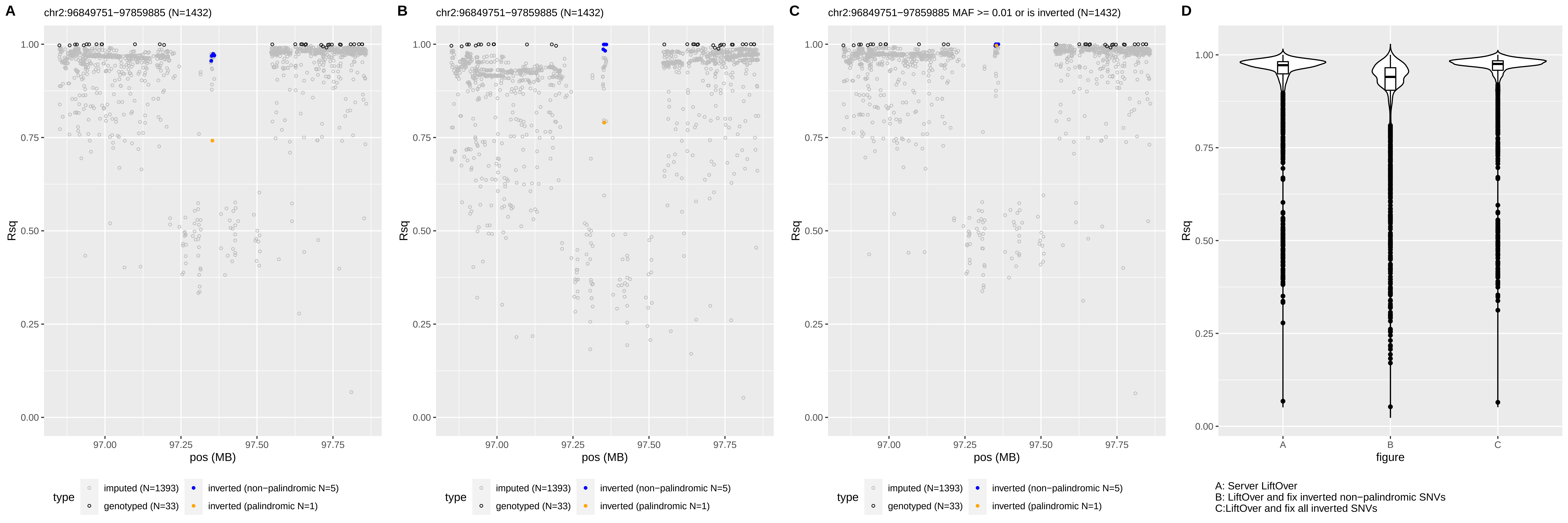

A
chr2:96849751−97859885 (N=1432)
B
chr2:96849751−97859885 (N=1432)
C
chr2:96849751−97859885 MAF >= 0.01 or is inverted (N=1432)
D
Rsq
pos (MB)
97.00
97.25
97.50
97.75
0.00
0.25
0.50
0.75
1.00
type
imputed (N=1393)
genotyped (N=33)
inverted (non−palindromic N=5)
inverted (palindromic N=1)
figure
A
B
C
A: Server LiftOver
B: LiftOver and fix inverted non−palindromic SNVs
C:LiftOver and fix all inverted SNVs

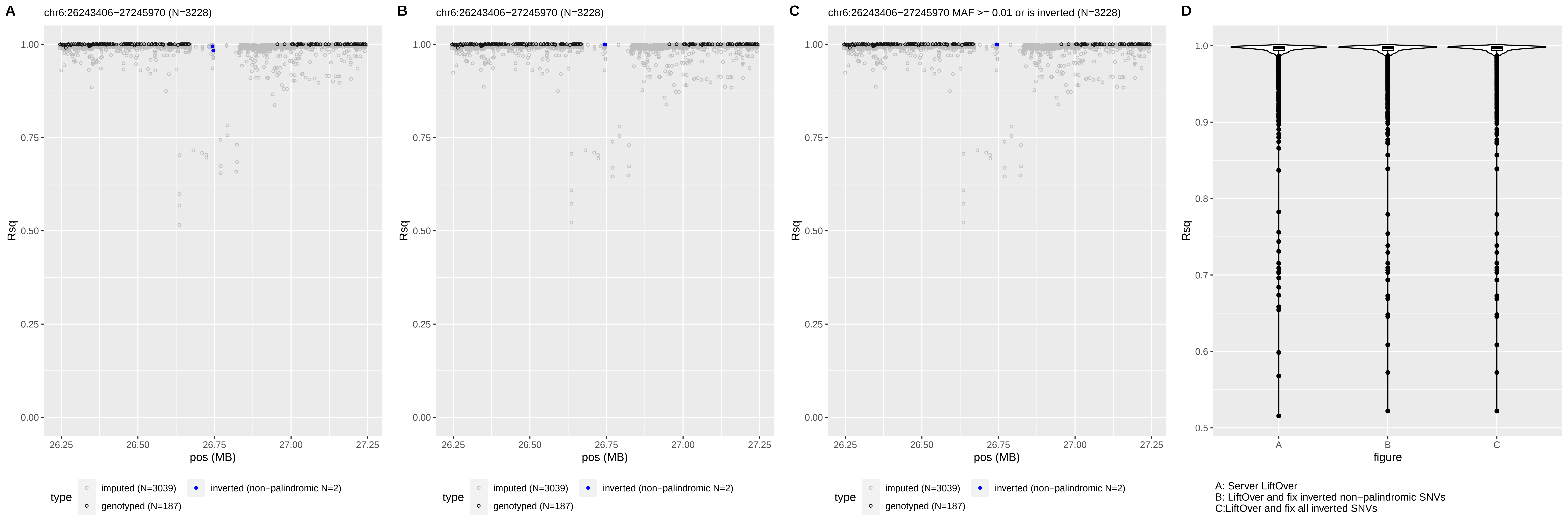

A
chr6:26243406−27245970 (N=3228)
B
chr6:26243406−27245970 (N=3228)
C
chr6:26243406−27245970 MAF >= 0.01 or is inverted (N=3228)
D
Rsq
pos (MB)
type
imputed (N=3039)
genotyped (N=187)
inverted (non−palindromic N=2)
figure
A: Server LiftOver
B: LiftOver and fix inverted non−palindromic SNVs
C:LiftOver and fix all inverted SNVs

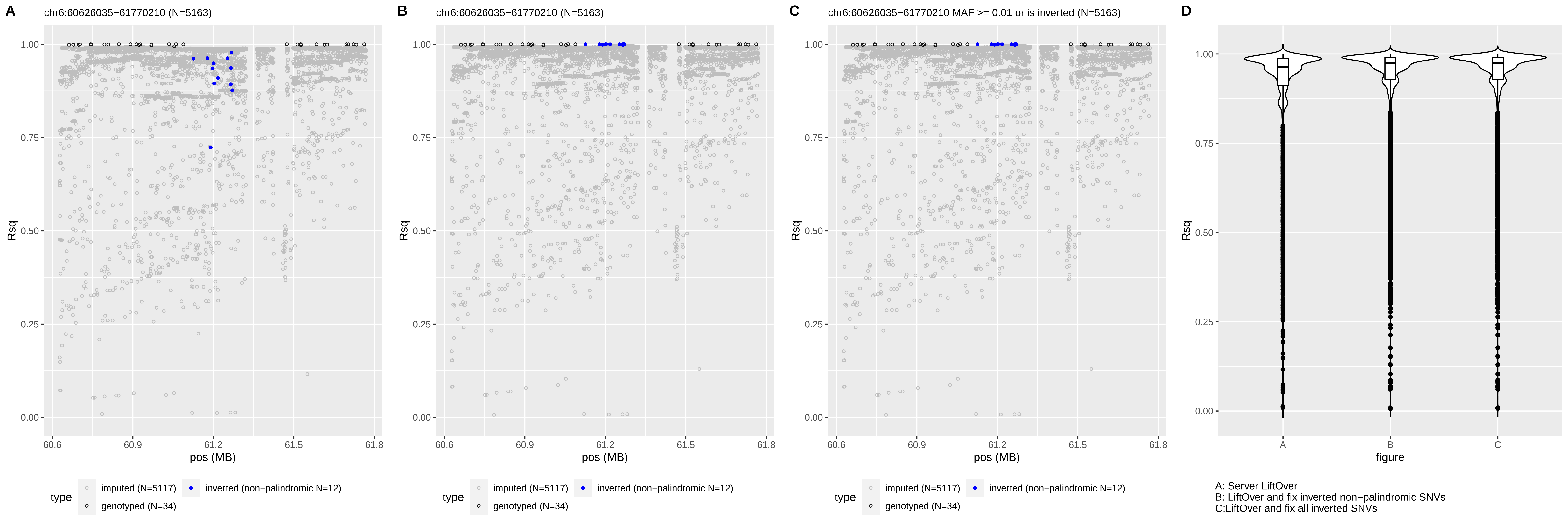
A
chr6:60626035−61770210 (N=5163)
B
chr6:60626035−61770210 (N=5163)
C
chr6:60626035−61770210 MAF >= 0.01 or is inverted (N=5163)
D
Rsq
pos (MB)
type
imputed (N=5117)
inverted (non−palindromic N=12)
genotyped (N=34)
figure
A: Server LiftOver
B: LiftOver and fix inverted non−palindromic SNVs
C:LiftOver and fix all inverted SNVs

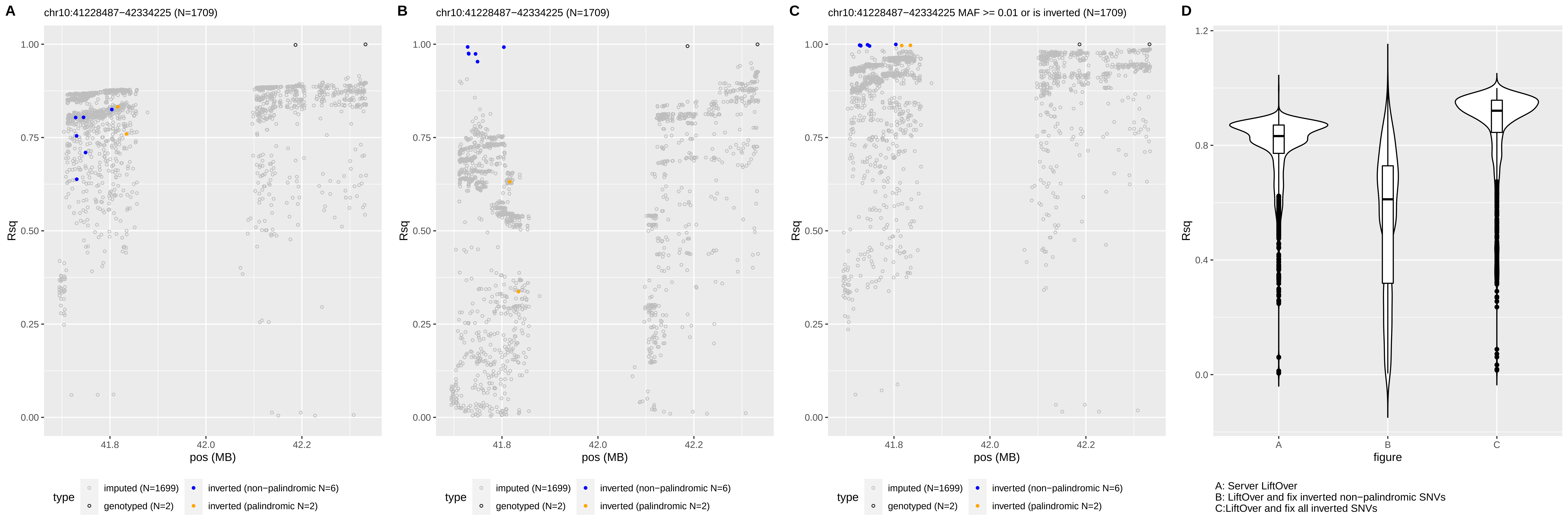

A
chr10:41228487−42334225 (N=1709)
B
chr10:41228487−42334225 (N=1709)
C
chr10:41228487−42334225 MAF >= 0.01 or is inverted (N=1709)
D
Rsq
pos (MB)
figure
type
imputed (N=1699)
genotyped (N=2)
inverted (non−palindromic N=6)
inverted (palindromic N=2)
A: Server LiftOver
B: LiftOver and fix inverted non−palindromic SNVs
C:LiftOver and fix all inverted SNVs

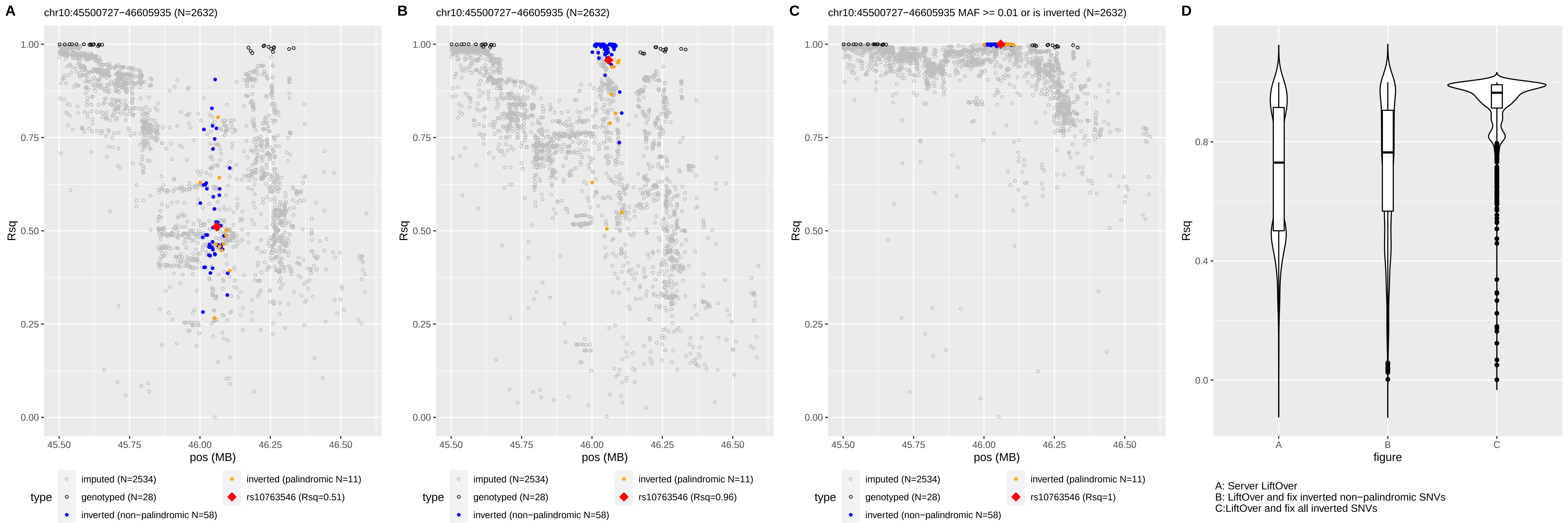
A
chr10:45500727−46605935 (N=2632)
B
chr10:45500727−46605935 (N=2632)
C
chr10:45500727−46605935 MAF >= 0.01 or is inverted (N=2632)
D
Rsq
pos (MB)
type
imputed (N=2534)
genotyped (N=28)
inverted (non−palindromic N=58)
inverted (palindromic N=11)
rs10763546 (Rsq=0.51)
rs10763546 (Rsq=0.96)
rs10763546 (Rsq=1)
figure
A: Server LiftOver
B: LiftOver and fix inverted non−palindromic SNVs
C:LiftOver and fix all inverted SNVs

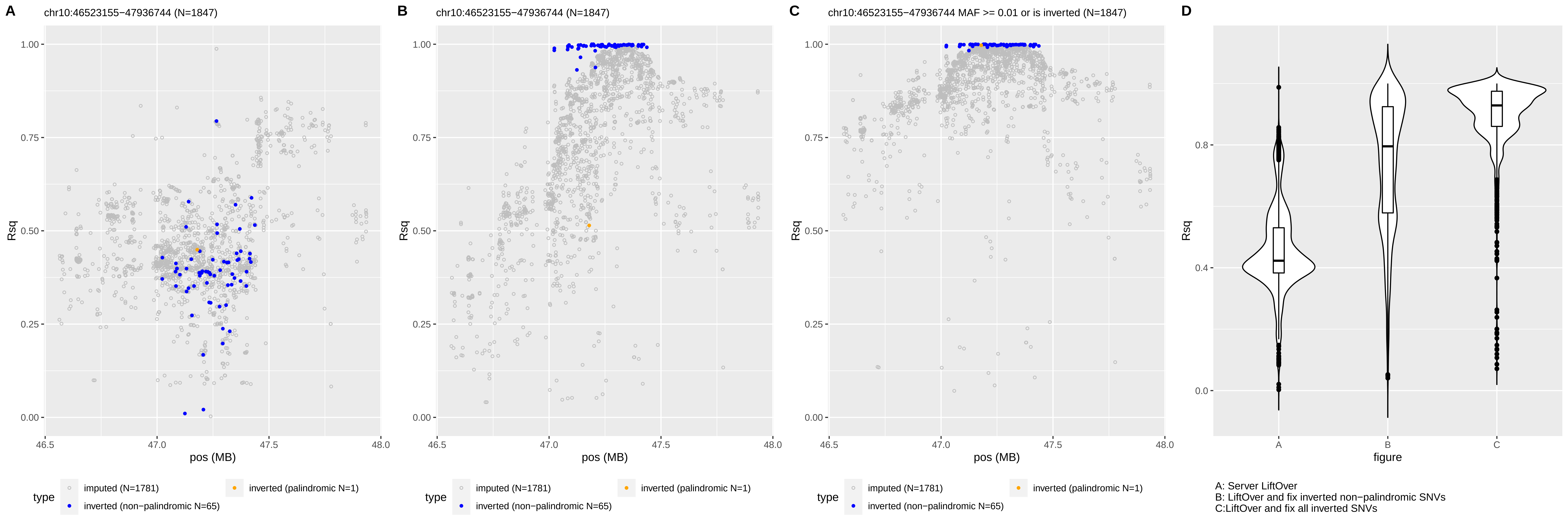
A
chr10:46523155−47936744 (N=1847)
B
chr10:46523155−47936744 (N=1847)
C
chr10:46523155−47936744 MAF >= 0.01 or is inverted (N=1847)
D
Rsq
pos (MB)
type
imputed (N=1781)
inverted (non−palindromic N=65)
inverted (palindromic N=1)
figure
A: Server LiftOver
B: LiftOver and fix inverted non−palindromic SNVs
C:LiftOver and fix all inverted SNVs

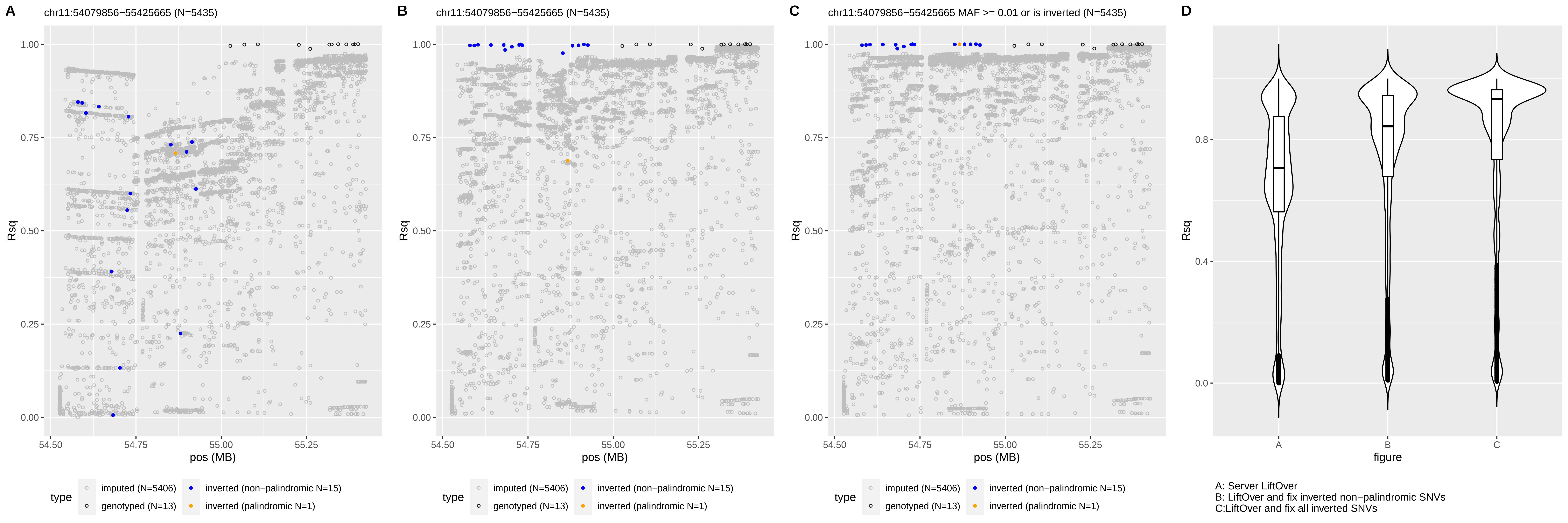

A
chr11:54079856−55425665 (N=5435)
B
chr11:54079856−55425665 (N=5435)
C
chr11:54079856−55425665 MAF >= 0.01 or is inverted (N=5435)
D
Rsq
pos (MB)
figure
type
imputed (N=5406)
genotyped (N=13)
inverted (non−palindromic N=15)
inverted (palindromic N=1)
A: Server LiftOver
B: LiftOver and fix inverted non−palindromic SNVs
C:LiftOver and fix all inverted SNVs

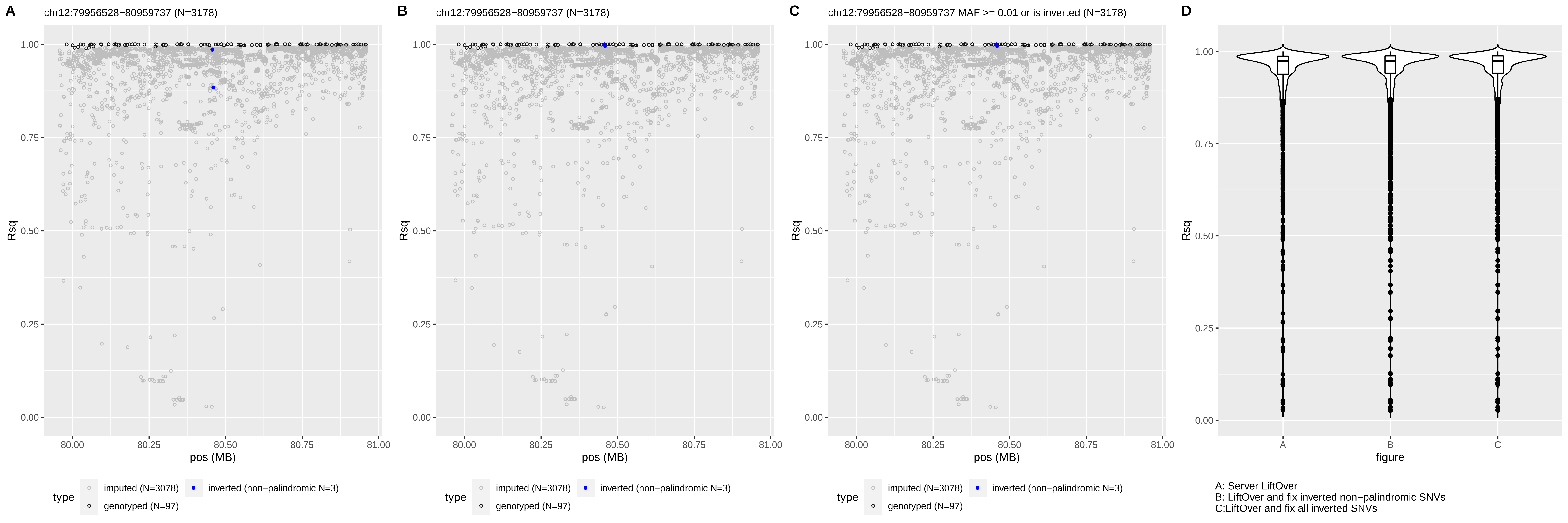

A
chr12:79956528−80959737 (N=3178)
B
chr12:79956528−80959737 (N=3178)
C
chr12:79956528−80959737 MAF >= 0.01 or is inverted (N=3178)
D
Rsq
pos (MB)
figure
type
imputed (N=3078)
inverted (non−palindromic N=3)
genotyped (N=97)
A: Server LiftOver
B: LiftOver and fix inverted non−palindromic SNVs
C:LiftOver and fix all inverted SNVs

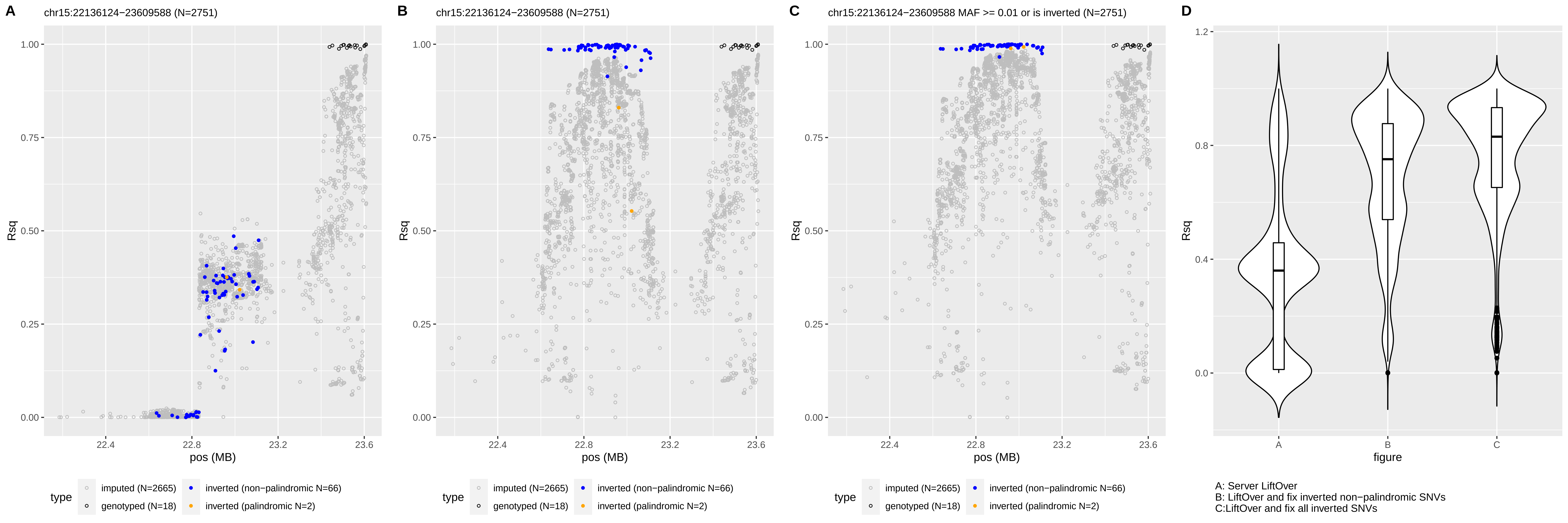
A
chr15:22136124−23609588 (N=2751)
B
chr15:22136124−23609588 (N=2751)
C
chr15:22136124−23609588 MAF >= 0.01 or is inverted (N=2751)
D
Rsq
pos (MB)
figure
type
imputed (N=2665)
genotyped (N=18)
inverted (non−palindromic N=66)
inverted (palindromic N=2)
A: Server LiftOver
B: LiftOver and fix inverted non−palindromic SNVs
C:LiftOver and fix all inverted SNVs

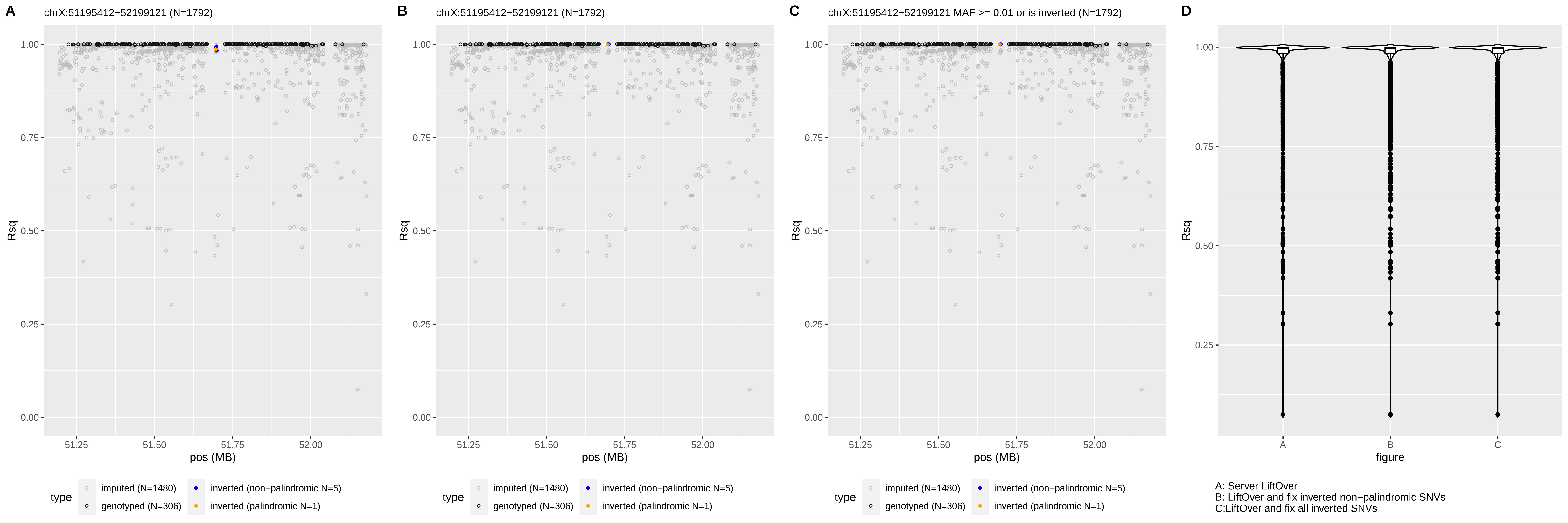

A
chrX:51195412−52199121 (N=1792)
B
chrX:51195412−52199121 (N=1792)
C
chrX:51195412−52199121 MAF >= 0.01 or is inverted (N=1792)
D
Rsq
1.00
0.75
0.50
0.25
0.00
pos (MB)
51.25
51.50
51.75
52.00
type
imputed (N=1480)
genotyped (N=306)
inverted (non−palindromic N=5)
inverted (palindromic N=1)
figure
A
B
C
A: Server LiftOver
B: LiftOver and fix inverted non−palindromic SNVs
C:LiftOver and fix all inverted SNVs

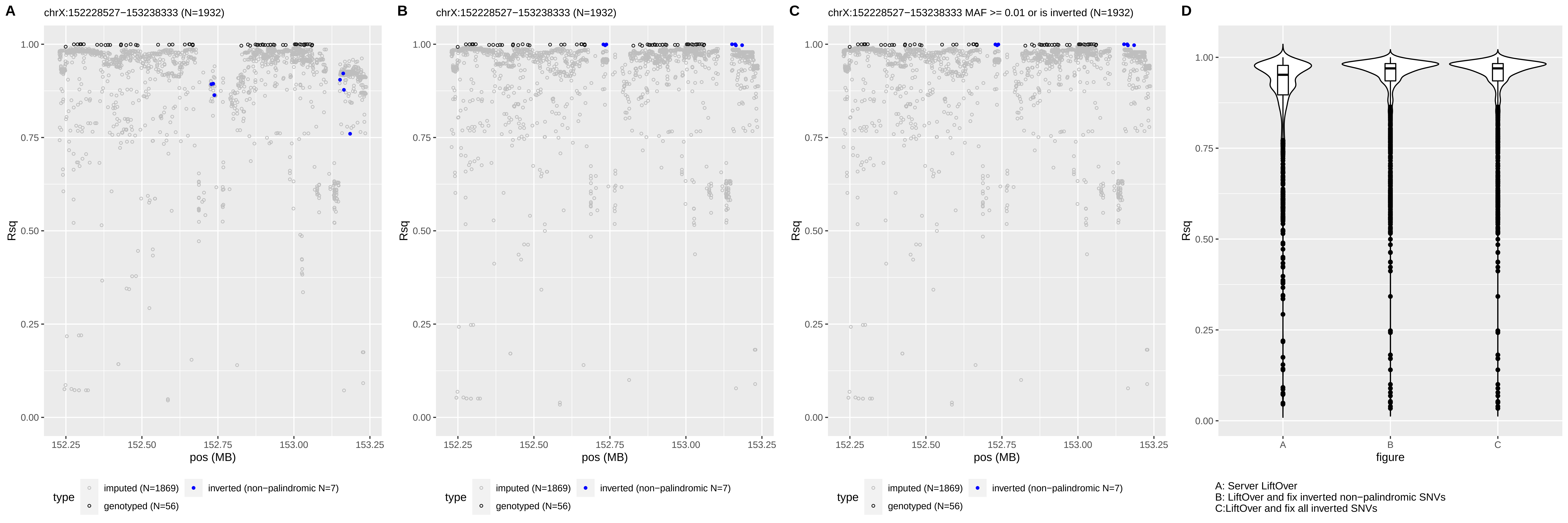

A
chrX:152228527−153238333 (N=1932)
B
chrX:152228527−153238333 (N=1932)
C
chrX:152228527−153238333 MAF >= 0.01 or is inverted (N=1932)
D
Rsq
pos (MB)
figure
type
imputed (N=1869)
inverted (non−palindromic N=7)
genotyped (N=56)
A: Server LiftOver
B: LiftOver and fix inverted non−palindromic SNVs
C:LiftOver and fix all inverted SNVs

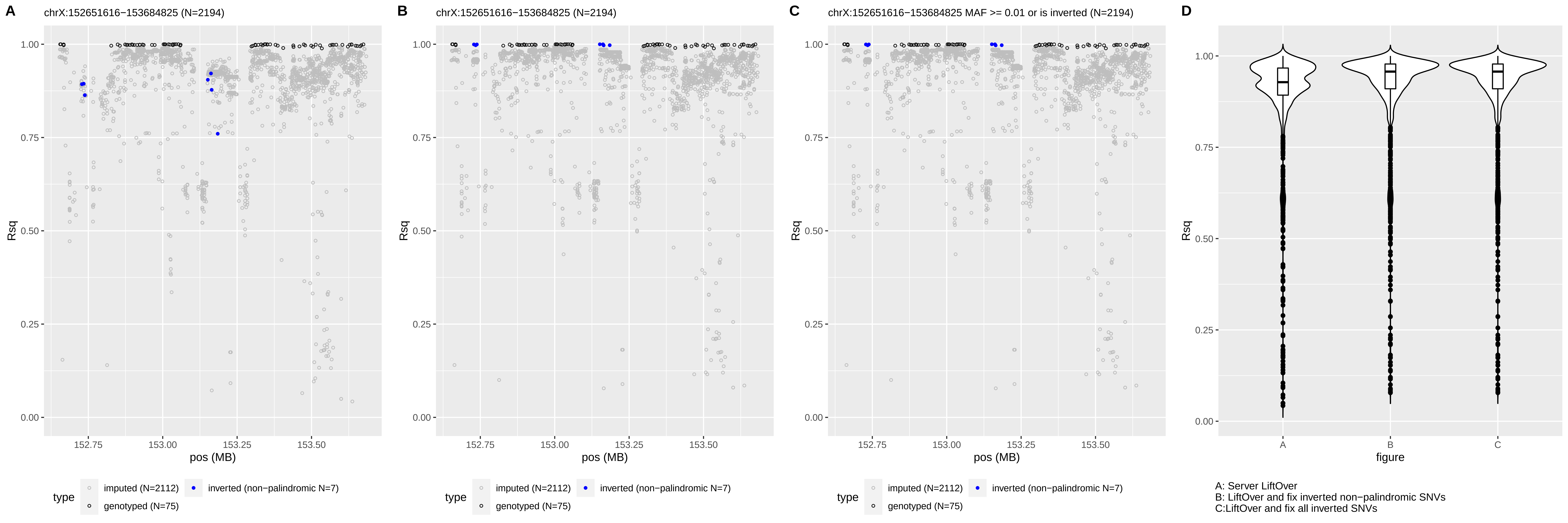

A
chrX:152651616−153684825 (N=2194)
B
chrX:152651616−153684825 (N=2194)
C
chrX:152651616−153684825 MAF >= 0.01 or is inverted (N=2194)
D
Rsq
pos (MB)
figure
type
imputed (N=2112)
inverted (non−palindromic N=7)
genotyped (N=75)
A: Server LiftOver
B: LiftOver and fix inverted non−palindromic SNVs
C:LiftOver and fix all inverted SNVs
